# Deterministic dynamics of distributional multi-agent reinforcement learning

**DOI:** 10.64898/2025.12.22.696014

**Authors:** Clémence Bergerot, Pawel Romanczuk, Wolfram Barfuss

## Abstract

Understanding how cognition shapes behavior across contexts remains a fundamental challenge for many disciplines. In particular, for the optimism heuristic–i.e., the tendency to overweight positive (relative to negative) information–knowledge remains fragmented, with models developed in specific domains in isolation. Here, we present a unifying computational framework by deriving the deterministic dynamics of distributional multi-agent reinforcement learning. Our approach discretizes reward distributions through a finite set of neurons, consistent with recent empirical findings on distributional coding in the brain. We validate our framework by reproducing established results across three iconic domains spanning individual exploration versus exploitation, social coordination, and risky choice. Beyond validation, we uncover novel interactions among optimism, reward discretization, and temporal discounting. Specifically, we identify conditions under which choice hysteresis and path-dependent strategies emerge, suggesting that perseveration results from neural reward discretization rather than constituting an independent heuristic. We further reveal “individual dilemmas”: circumstances where agents gravitate toward suboptimal yet stable strategies, offering a mechanistic explanation for incoherent choice patterns. Our framework bridges neuroscience, psychology, and collective behavior, enabling empirically testable hypotheses about how cognitive biases propagate from individual cognition to social outcomes in complex environments.

## 1 Introduction

To advance fields such as biology, economics, artificial intelligence (AI), and sustainability, we need an adequate model of how behavior arises from biological and cognitive first principles. In biology, the study of collective behavior began to incorporate the latter [1, 2]. Likewise, evolutionary game theorists were called upon to include cognitive mechanisms [3, 4]. In economics, non-standard choice patterns are increasingly explained by foundational aspects of cognitive information processing [5]. In AI, scientists have advocated for the interdisciplinary study of machine behavior and machine culture, analyzing machine behavior in society similarly to that of animals [6, 7]. Additionally, sustainability scientists argued that insights into human behavior and cognition, embedded in a complex systems approach, are key to developing policies towards a sustainable future [8, 9].

At the behavioral level, there are many biases, heuristics, and behavioral theories– and it is not clear how they relate to each other. Knowledge about human behavior is fragmented into many different, context-specific, and often informal theories. For example, in an attempt to order and use this knowledge for sustainability science, Constantino and colleagues presented a selection of 32 distinct behavioral theories [9], some of which are specific biases or heuristics. Originating from Kahneman and Tversky’s pioneering work [10], biases are defined as systematic deviations from rationality in judgment or decision-making. Systemic, here, means that these deviations are non-random and, thus, predictable [11]. Kahneman and Tversky’s fertile program has led to a proliferation of identified biases; yet, an overarching explanation for their origin or adaptive character is lacking [12]. For example, Wikipedia’s “List of cognitive biases” [13] contains more than 100 entries. Prominent examples include the confirmation bias, the anchoring bias, and the availability bias. As a result of regarding them as deviations from rationality, biases are often seen as flaws or fallacies, which implicitly re-establishes neoclassical rationality as a norm to be achieved.

Another school of thought, pioneered by Gigerenzer, calls these biases “heuristics” to stress their ecological rationality [14]. Heuristics are considered as “fast and frugal” cognitive strategies that bring about good decisions in complex, uncertain environments. From this viewpoint, heuristics are not a bug, but rather a feature of human decisionmaking. Although ecological rationality is a unifying feature of heuristics, these are still considered multiple tools within an “adaptive toolbox” [15]. Prominent examples include the recognition heuristic, the fluency heuristic, and fast-and-frugal decision trees. Moreover, for a long time, psychologists and economists have studied heuristics with little regard for biological implementation and neural underpinnings. Therefore, it is unclear why heuristics are so pervasive among humans and animals, and which brain mechanisms support them [16].

At the cognitive level, Reinforcement Learning (RL) has emerged as a framework for integrating cognitive mechanisms–such as learning, representation, and decisionmaking–across the neuro, cognitive, behavioral and machine learning sciences [17, 18, 19]. RL is a trial-and-error method in which an agent aims to improve its behavior while receiving continuous feedback from its environment. Various concepts from behavioral theories, including biases and heuristics, have been implemented in RL approaches. For instance, in robotics, intrinsic motivations–such as curiosity, drive and open-ended development [20]–are incorporated into various RL algorithms for complex tasks such as tool use [21, 22, 23]. In machine learning, heuristic RL improves agent performance in different settings, with social influence enhancing coordination [24] and optimism promoting cooperation [25]. In cognitive psychology, modified RL algorithms help study behaviors like perseveration, optimism, and confirmation bias, shedding light on decision-making processes [26, 27].

In particular, optimism and pessimism are widely studied heuristics. An agent is defined as “optimistic” when it puts more weight on positive, than on negative, information. In cognitive psychology, optimism has been modeled via so-called asymmetric RL, which uses two different learning rates. A higher learning rate updates positive prediction errors, while a lower one updates negative ones. Several experimental studies have documented the presence of optimism in humans [28, 29], mice [30], and monkeys [31] performing different versions of a two-armed bandit task. Moreover, asymmetric RL has been shown to solve a variety of problems when applied to the design of decentralized Multi-Agent Reinforcement Learning (MARL) systems. In MARL, researchers study systems that are composed of two or more virtual agents; these agents learn and aim to solve a task together. The design of several decentralized algorithms with an optimism/pessimism heuristic, such as Lenient Learning [32], Hysteretic Q-learning [25], Win-or-Learn-Fast Policy Hill-Climbing [33], and variants of Frequency Maximization Q-learning [34, 35], has produced MARL systems with enhanced coordination and improved performance on various tasks. However, these models and algorithms must often treat optimism and pessimism as ad-hoc hyperparameters. Moreover, they cannot explain why specific levels of optimism or pessimism are helpful in a given situation.

Distributional RL provides the representational richness needed to meaningfully implement and reason about optimism/pessimism. In the past few years, optimism has been studied within a distributional framework. Here, agents do not compute the expected value of each option, but the full distributions of reward outcomes [36]. Optimism is thus not implemented by asymmetric learning rates, but by the selection of an upper quantile or expectile of the distribution instead of the mean [37, 38], or by a distortion of the shape of the distribution [39, 40]. Doing so allows for improved treatment of uncertainty [41]. Distributional Reinforcement Learning (RL) characterizes aleatoric (environmental) uncertainty by using value distributions rather than single-point estimates. This methodology enables the agent to differentiate between uncertainty stemming from environmental stochasticity and that due to insufficient exploration, which is crucial for deciding when to be optimistic (explore) vs. pessimistic (exploit safely) during learning. Interestingly, neural signatures in the brain have been shown to align with a distributional RL perspective [42], thereby adding biological plausibility to it.

However, all these algorithms and learning processes of optimism and pessimism have mostly been used in specific contexts only, such as uncertain, risky, or collective choices. On the one hand, experimental psychologists’ asymmetric RL focuses on individual subjects performing basic tasks. Only a few agent-based modeling works have begun to study the influence of optimism via asymmetric learning rates in social settings and complex environments [43, 44, 45]. On the other hand, machine learning scientists’ optimistic MARL and distributional RL handle higher levels of social and environmental complexity. However, they are not primarily aimed at studying human decision-making and tend to develop multiple algorithmic variants to solve specific machine-learning problems.

As a result, the current state of knowledge lacks a unified, biologically plausible framework for modeling heuristics in RL that is applicable across various choice domains. Combining multi-agent reinforcement learning with complexity science and evolutionary dynamics can help integrate cognitive, social, and environmental factors in human decision-making into a coherent, understandable framework [46, 47]. Two characteristics of this approach facilitate understandability. First, using deterministic RL dynamics, where all relevant quantities in the learning processes (such as rewards and values) are averaged over the agents’ strategies and the environment’s state transitions [48, 49, 50, 51]. Second, using idealized choice environments that aim to capture the phenomenon under investigation without introducing additional complexity, i.e., as simple as possible but not simpler. Together, these two characteristics offer a computationally efficient and understandable way to study how, under given environmental conditions, individual decisions give rise to collective behavior.

Thus, with this work, we aim to combine deterministic RL dynamics with a distributional approach to devise a broadly applicable framework for optimism/pessimism across different environments. We start by deriving distributional deterministic RL (DDRL) dynamics and propose a way to model the optimistic heuristic within this framework (Section 2. A key aspect of our model is the discretization of the continuous reward spectrum by employing stylized reward neurons, in line with empirical findings [42]. Second, we illustrate the broad applicability of DDRL across relevant scenarios by testing it in three choice environments: an exploration-exploitation challenge, a social coordination task, and an intertemporal risky choice problem (Section 3). On the one hand, our findings replicate previous results, indicating the unifying power of our model: The impact of optimism on the exploration-exploitation challenge is modulated by resource scarcity; optimism enhances coordination; and optimism increases risk-seeking in our intertemporal risky choice problem. On the other hand, we show how our model gives rise to previously unknown effects, stemming from the interaction between optimism/pessimism, reward discretization, and intertemporal choice: the emergence of bistable choice regimes; and a form of incoherent behavior we call “individual dilemma”. In such a dilemma, a single agent values another choice more than their current one, yet cannot perform a behavioral change. We hope to pave the way for more biologically grounded models of multi-agent decision-making and collective learning. Moreover, we believe our framework can be helpful to cognitive scientists and experimental psychologists for two main reasons. First, because the deterministic aspect of our model can help understand the role of noise in the learning process [52]; second, because the discretized and distributional reward spectrum gives rise to unique psychological phenomena.

## 2 Methods

Our framework combines deterministic RL dynamics [50] with distributional reinforcement learning [36]. We start with presenting the relevant background of our framework: stochastic games, reinforcement learning (RL), distributional RL (DistRL), and deterministic RL dynamics (DetRL). From there, we derive the deterministic distributional RL (DDRL) dynamics. Finally, we introduce the weighing algorithm we use to bias the learning dynamics towards optimism or pessimism.

### 2.1 Background

#### 2.1.1 Stochastic games

Stochastic games, also known as Markov games, are defined by the elements ⟨*N*, 𝒮, ***𝒜***, *T*, ***R***⟩*N* ∈ ℕ agents reside in an environment of *Z* ∈ ℕ states 𝒮 = (*S*_1_, …, *S*_*Z*_). In each state *s*, each agent *i* ∈ 𝒩 = {1, …, *N* } has *M* ∈ ℕ available actions 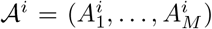 to choose from. ***𝒜*** = ⊗_*i*_ 𝒜^*i*^ is the joint-action set where ⊗_*i*_ denotes the cartesian product over the sets indexed by *i*. Time advances in discrete steps, and agents choose their actions simultaneously. A joint action is denoted by ***a*** = (*a*^1^, …, *a*^*N*^ ) ∈ ***𝒜***. With ***a***^*−i*^ = (*a*^1^, …, *a*^*i−*1^, *a*^*i*+1^, …, *a*^*N*^ ) we denote the joint action except agent *i*’s, and we write the joint action in which agent *i* chooses *a*^*i*^ and all other agents choose ***a***^*−i*^ as *a*^*i*^***a***^*−i*^. We chose an equal number of actions for all states and agents out of notational convenience.

The **transition** tensor ***T*** : 𝒮 × ***𝒜*** × 𝒮 ⟶ [0, 1] determines the probabilistic state change. *T* ^*s*,***a***,*ś*^ is the transition probability from current state *s* to next state *ś* under joint action ***a***.

The **reward** tensor ***R*** : 𝒩 × 𝒮 × ***𝒜***× 𝒮 ⟶ ℛ maps the triple of current state *s*, joint action ***a*** and next state *ś* to an immediate reward scalar for each agent. *R*^*i,s*,***a***,*ś*^ is the reward agent *i* receives.

At each time step, actions are chosen from the joint **strategy** tensor ***X*** : 𝒩 × 𝒮 ×*𝒜* ⟶ [0, 1]. In state *s*, agent *i* chooses action *a*^*i*^ with probability 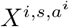 .

Each agent *i* aims for a strategy 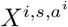to maximize their **gain**,

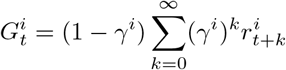

where 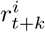 denotes the reward agent *i* obtained at time step *t* + *k*. The discount factor, *γ*^*i*^ ∈ [0, 1), denotes how much agent *i* cares for future rewards.

#### 2.1.2 Reinforcement learning

Reinforcement learning is a trial-and-error method of mapping situations to actions to maximize a numerical reward signal [18]. When rewards are a delayed consequence of current actions, so-called temporal-difference or reward-prediction learning has been particularly influential. This type of learning summarizes the difference between value estimates from past and present experiences into a reward-prediction error, which is then used to adapt the current behavior to gain more rewards over time. There also exists remarkable similarities between computational reinforcement learning and the results of neuroscientific experiments [53]. Dopamine conveys reward-prediction errors to brain structures where learning and decision-making occur [54].

At time step *t*, in state *s*, agent *i* evaluates action *a*^*i*^ to be of quality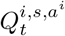. Learning then means updating these state-action value estimates, 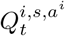, after selecting action 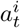 and having observed state *s*_*t*_ according to

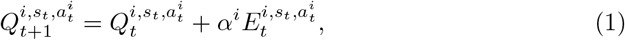

where *α*^*i*^ ∈ [0, 1] denotes agent *i*’s learning rate, which regulates how much new information the agent uses for the update, and 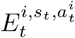 the current temporal-difference or reward-prediction error. The expression of the temporal-difference error depends on the chosen RL algorithm.

In this work, we focus on two different RL variants: SARSA learning, and Actor-Critic learning. SARSA learning takes into account the five elements of the current State, current Action, Reward, next State, and next Action for one learning update step. In Actor-Critic learning, an approximation of the State-Action value gets “criticized” by (i.e., compared against) an approximation of the state value. Another well-known variant is Q-learning. A comparison of these three variants can be found in [50].

In SARSA learning, the temporal-difference error is expressed as:

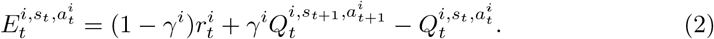

In Actor-Critic learning, by contrast, the temporal-difference error is a comparison between the state-action value at time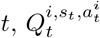, and the state value at time *t*,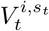 . Given that 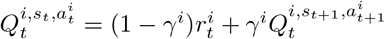,

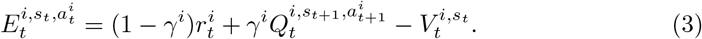

The state value at time 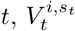, quantifies the overall quality of state *s* at time *t*, regardless of the chosen action *a*_*t*_. In Actor-Critic learning, it is separately updated according to:

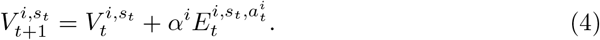

In both learning variants, the quality of action *a*^*i*^ is also described by the Bellman equation:

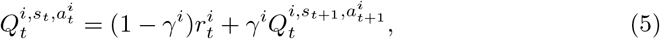

which forms the first part of the expression of the prediction error in Eq. 2 and Eq. 3.

Agents select actions based on the current state-action value beliefs 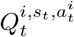, balancing exploitation (i.e., choosing the action of maximum quality) and exploration (i.e., selecting lower-quality actions to learn more about the environment). We employ the widely used Boltzmann policy function. The probability of choosing action *a*^*i*^ under observing state *s* is

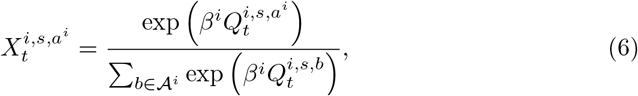

where the *intensity-of-choice* parameters, *β*^*i*^ ∈ ℝ^+^, regulate the exploration-exploitation trade-off. For high *β*^*i*^, agents exploit their learned knowledge about the environment, leaning toward actions with high estimated state-action values. For low *β*^*i*^, agents are more likely to deviate from these high-value actions to explore the environment further with the chance of finding actions that eventually lead to even higher values.

#### 2.1.3 Distributional reinforcement learning

Distributional approaches to RL (DistRL) recently became popular in artificial intelligence and machine learning research. DistRL diverges from the classical approach insofar as learning agents do not compute expected values, but rather the full distribution of outcomes. Therefore, the goal is not to compute an expected value or quality, but rather a **value distribution**, which is described by a distributional Bellman equation [36]:

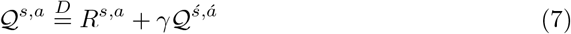

where *R*^*s,a*^ is the reward for action *a* in state *s*, (*ś, á*) is the next-state action, and Q is its random return.

Over the past years, DistRL researchers have used various supports for discretizing the distributions. In categorical DistRL, the discretization support is a set {*z*_*i*_ = *V*_*min*_ + *i*Δ*z* : 0 ≤ *i < N* } of *N* categorical “atoms”, which are fixed and evenly spaced (within 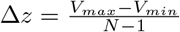) [36]. Later versions of DistRL algorithms rely on the distributions’ statistics as discretization support: the atoms are not fixed anymore, but are updated so as to converge towards quantiles or expectiles of the target distribution [55].

In this work, we will use a categorical discretization support for the sake of simplicity.

#### 2.1.4 Deterministic learning dynamics

Deterministic approximations of RL remove noise from the learning process. Hence, the agents’ strategies over time can be computed and visualized as deterministic trajectories in phase space. To this end, we use temporal-difference RL in the deterministic limit (DetRL), which was developed by Barfuss *et al*., [50]. This method uses strategy-averages as a way to remove the noise from the learning dynamics: the prediction error is averaged over all strategies and state transitions.

First, we compute the different quantities we need to calculate the average temporal-difference error. As explained in Section 2.1.1, 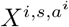 denotes the probability that agent *i* chooses action *a*^*i*^ in state *s*; ***X*** : 𝒩 × 𝒮 × 𝒜 ⟶ [0, 1] denotes the joint strategy tensor.

We call 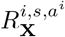 the average reward given the joint policy **X** that agent *i* obtains by taking action *a*^*i*^ in state *s*. More precisely, the average is taken over state transitions from *s* to *ś*, and over other agents’ actions ***a***^***−i***^:

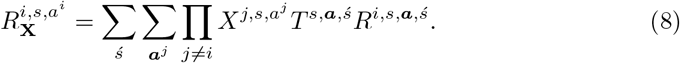

Similarly, we call 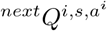 the average next-state Q-value given the joint policy **X** for agent *i* taking action *a*^*i*^ in state *s*:

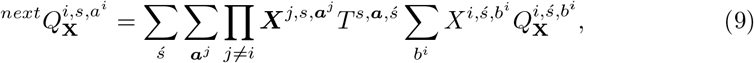

where 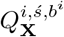denotes the average Q-value of action *b*^*i*^ taken by agent *i* in state *ś*, described by the recursive Bellman equation:

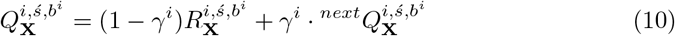

Therefore, for a SARSA agent, the average prediction error for agent *i* taking action *a*^*i*^ in state *s* at time *t* is

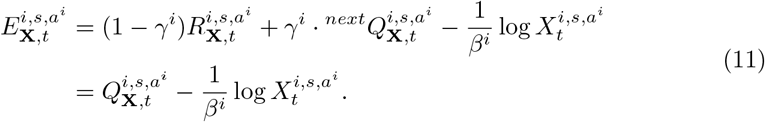

where *β*^*i*^ is the choice intensity, or inverse temperature, of agent *i*.

For an Actor-Critic agent, by contrast, the average Q-value 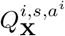 is compared with the average state value 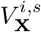 (the “critic”). Since this term is constant in action, we can omit it from the formula [50]. Thus, for an Actor-Critic agent, the average prediction error for agent *i* taking action *a*^*i*^ in state *s* at time *t* is:

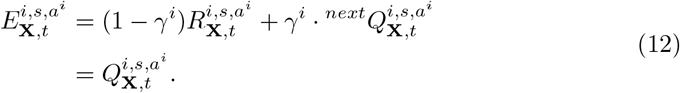

The combination of the Q-value update (eq. 1) and the Boltzmann formula (eq. 6) yields the deterministic update of strategy 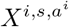 :

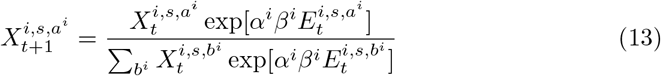

### 2.2 Deriving deterministic distributional reinforcement learning dynamics

In order to model the optimism-pessimism heuristic, we apply deterministic learning dynamics to distributional RL. By doing so, we combine the best of both worlds: on the one hand, DetRL provides a transparent, reproducible, and computationally fast approach to RL; on the other hand, DistRL allows us to bias the value distribution of the learning dynamics and devise a simple model at the neuronal level.

Our approach is presented in Fig 1. The agent-environment interface is formalized as a Markov game ⟨*N*, 𝒮, ***𝒜***, *T*, ***R***⟩ comprised of *N* agents, a set 𝒮 of different states, a set ***𝒜*** of available actions, a transition tensor *T*, and a reward tensor ***R***. In DDRL, rewards and qualities are not computed as expected values, but as full distributions. These distributions are discretized according to a categorical support and a smoothing procedure. Then, the Q-value distribution is distorted according to a weighing algorithm, implementing the optimism heuristic.

**Figure 1:**
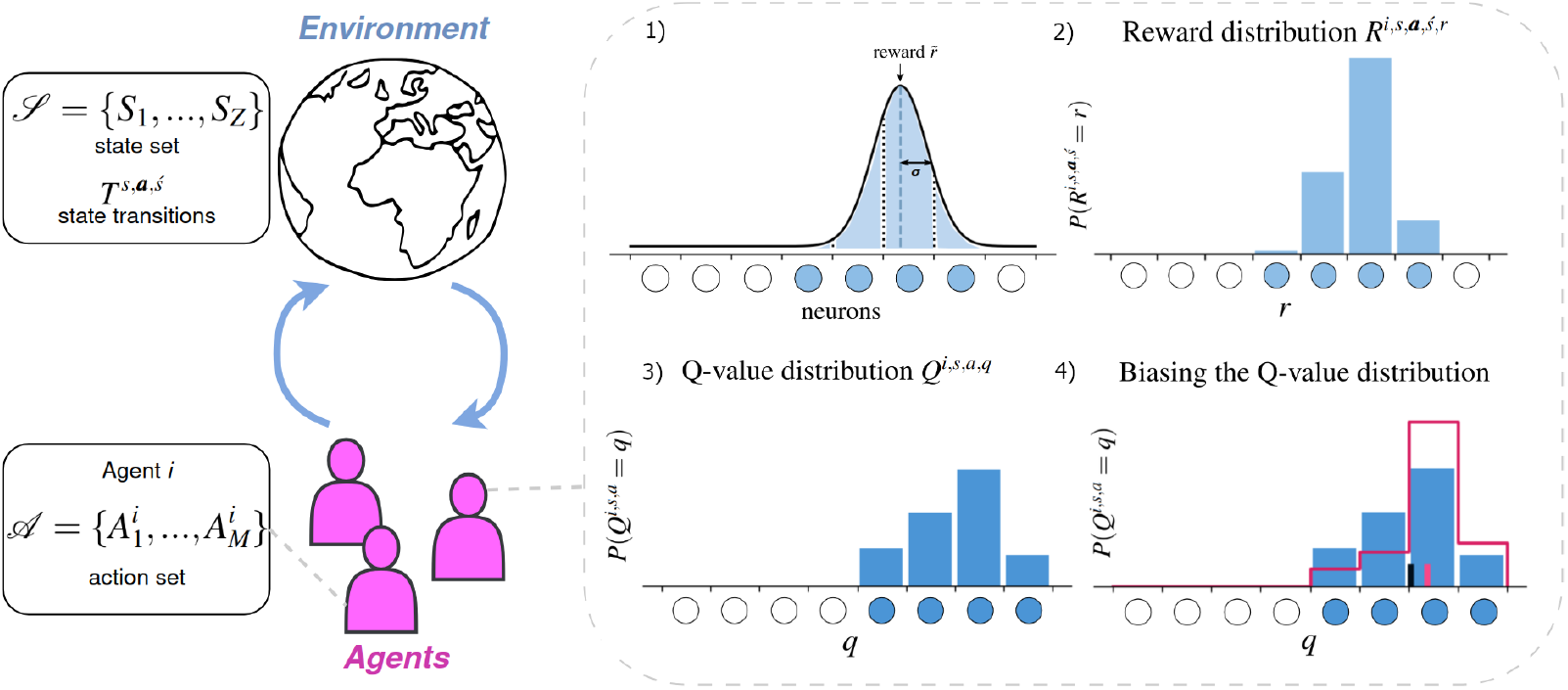
The agent-environment interface. *N* agents play in an environment formalized as a set of *Z* states 𝒮 = {*S*_1_, …, *S*_*Z*_}. In each state, agent *i* can choose between *M* actions from the action set 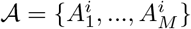. The probability that a transition from state *s* to state *ś* occurs when agents play the joint action ***a*** is *T* ^*s*,***a***,*ś*^. The right panel represents reward discretization and Q-value estimation for one agent *i*. 1) An environmental reward quantity, 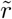, is detected with sensitivity *σ* by several neurons (in light blue). Each neuron *k* represents the probability that the environmental reward equals a quantity *r*_*k*_ ± Δ*r*; this probability is the area, taken between *r*_*k*_ − Δ*r* and *r*_*k*_ + Δ*r*, under the Gaussian curve centered around 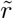 with standard deviation *σ*. 2) This yields the discretized reward distribution *R*^*i,s*,***a***,*ś,r*^. In DDRL, all quantities are no longer calculated as average estimates, but rather as discretized distributions. 3) Example Q-value distribution, which is calculated according to our DDRL dynamics (Section 2.2). 4) The optimistic heuristic is applied to the Q-value distribution (in blue). It transforms the original distribution into a distorted one (outlined in magenta). The mean of this new distribution (magenta bar) is an optimistic estimate of the average Q-value (black bar).

In the following subsections, we explain how we discretize and compute the relevant distributions.

#### 2.2.1 Discretization and reward distribution

We assume an agent has *n* ∈ ℕ reward “neurons”, each neuron being sensitive to a given range of amount of reward. We define *r*_1_ ∈ ℝ (*resp. r*_*n*_ ∈ ℝ) the minimal (*resp*. maximal) amount of reward that can be detected. Similarly to categorical DistRL [55], we assume a regular grid (*r*_*k*_)_*k∈*[1,*n*]_ such that *r*_*k*_ = *r*_1_ + (*k* − 1)Δ*r*, where 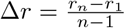 denotes the width separating two consecutive grid points.

Fig. 1 describes the process of discretization by reward neurons. In particular, it shows how an arbitrary reward amount 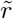 gets represented as a discretized distribution by defining a Gaussian kernel centered around 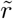 To smooth out the effects of discretization, we assume that each neuron *k* is maximally sensitive to the reward amount *r*_*k*_, quantified by a discretization standard deviation, *σ*. Each reward neuron *r*_*k*_ “fires” with intensity 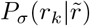, which quantifies the probability that 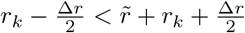. More precisely, 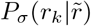 is calculated as the area between 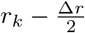 and 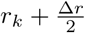 under the Gaussian bell curve centered around 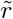 and with standard deviation *σ*,

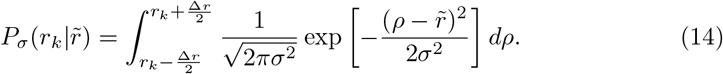

Therefore, this smoothing procedure captures some uncertainty about the categorization into different bins. This uncertainty is also modulated by the discretization standard deviation, *σ*. Ultimately, this discretization method allows us to transform any stochastic game’s reward tensor into a reward distribution. For all *r* ∈ {*r*_1_, …, *r*_*n*_}, we define the reward distribution *R*^*i,s*,***a***,*ś,r*^ for agent *i*, current state *s*, joint action ***a***, and next state *ś*, as:

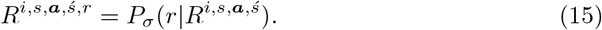

Note that each *R*^*i,s*,***a***,*ś*^ is an entry of the original reward tensor ***R***. In other terms, *R*^*i,s*,***a***,*ś,r*^ is the area between 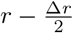 and 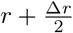under the bell curve centered around *R*^*i,s*,***a***,*ś*^.

#### 2.2.2 Reward distributions 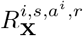 and 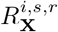

The previous reward distribution can be averaged over joint actions ***a*** and next states *ś*. We call the 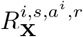 reward distribution for agent *i* taking action *a*^*i*^ in state *s*, averaged over transitions to the next state *ś*, and over other agents’ actions ***a***^*j*^:

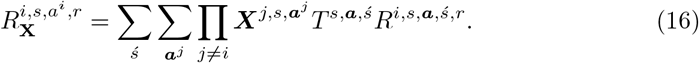

The reward distribution 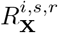 denotes the probability that agent *i* receives reward *r* in state *s*. It is averaged over joint actions ***a*** (including agent *i*’s) and next states *ś*:

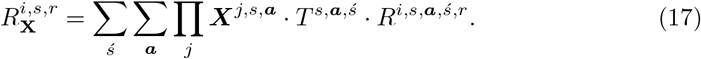

#### 2.2.3 Bellman equation and smoothed Bellman Kronecker 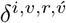

In traditional, expected-value RL, the value of a state *s* for agent *i* is computed via the Bellman equation:

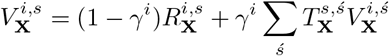

where 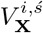 is the value of the next state, *ś*, and 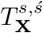 is the effective state transition matrix given joint policy **X**.

In distributional form, we want to compute the probability that (discretized) values *r, v* and 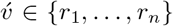 satisfy the Bellman equation 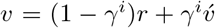. To this end, we introduce a smoothed Bellman Kronecker delta,

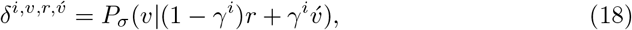

using the same discretization method as above. The idea is that 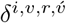 peaks when the Bellman condition 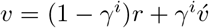 is fulfilled for agent *i*, and its reward bins *v, r*, and 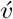, allowing for some uncertainty, *σ*, around the discretization into different bins. We use this smoothed Bellman Kronecker delta to compute the return or value distribution 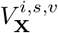 in closed form.

#### 2.2.4 Value distribution 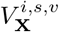

The value distribution 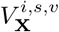 represents the probability that agent *i* in state *s* receives a long-term return *v*. It results from four instances that must occur together. For each short-term reward bin *r* ∈ ℛ, for all possible next states *ś* ∈ 𝒮, and each long-term return bin 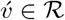, i) *v, r* and 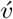 must fulfill the Bellman equation according to probability 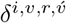, ii) agent *i* must receive short-term reward *r* in state *s*, according to probability 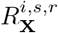, iii) state *s* must transition to state *ś*, according to probability 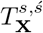, and finally iv) in state *ś*, agent *i* must receive a long-term return 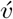. More concisely,

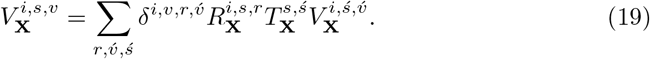

We now summarize the smoothed Bellman-Kronecker delta, reward, and transition probabilities into a tensor 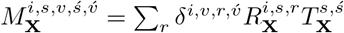. Doing so, we can rewrite Eq. 19 as 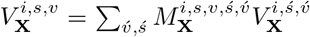. We now combine the indices *s* × *v* = *w* and 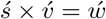, which transforms Eq. 19 into,

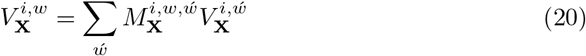

In other terms, for each agent *i* and joint policy *X*, the value distribution 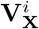 fulfills 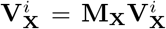. Thus, 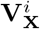 is the eigenvector of the matrix **M**_**X**_ corresponding to eigenvalue 1, normalized in such a way that for each state *s*, the probabilities to receive values *v* ∈ ℛ sum up to 1, 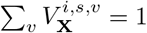.

#### 2.2.5 Q-value distribution 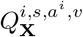

The probability for agent *i* to receive long-term return *v* in state *s* under action *a*^*i*^ is the joint probability that from all possible short-term reward values *r*, all possible next states *ś*, and all possible next long-term returns 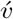, i) *v, r* and 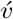 fulfill the Bellman equation according to probability 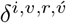, ii) agent *i* receives short-term reward *r* in state *s* under action *a*^*i*^, according to probability 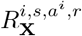, iii) state *s* transitions to state *ś* under action *a*^*i*^, according to probability 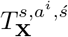, and iv) in state *ś*, agent *i* receives a long-term return 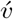, according to probability 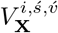. More concisely,

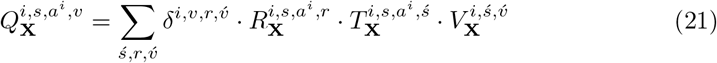

#### 2.2.6 Temporal-difference error

The SARSA and Actor-Critic temporal-difference errors differ slightly (see Section 2.1.4). Noting ⟨.⟩_*d*_ the mean of a distribution on dimension *d*, the deterministic approximation of the temporal-difference error at time *t* for a SARSA agent *i* taking action *a* in state *s*, 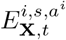, is:

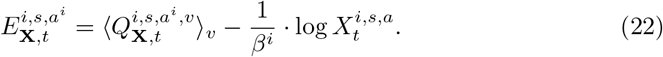

For an Actor-Critic agent, the last term is not needed, as the “critic” (*i*.*e*., 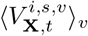) is constant in action (see Section 2.1.4). Therefore, the deterministic approximation of the temporal-difference error at time *t* for an Actor-Critic agent *i* taking action *a* in state *s*, 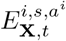, is:

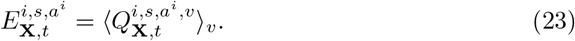

### 2.3 Biasing the Q-value distribution

Optimism (*resp*. pessimism) causes agents to overestimate (*resp*. underestimate) Q-values. There are different ways to obtain an optimistic (*resp*. pessimistic) estimate of a distribution. Here, we consider a weights method. In SI-Section 12.1, we show its advantages over two other methods from the literature, using quantiles and expectiles.

The weights method involves applying a weights tensor, 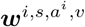, to the discretized distribution 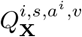 so as to distort it. Although there are different ways to distort a distribution, we use a stepwise function for simplicity, and because it is equivalent with asymmetric RL on a two-armed bandit task (see SI-Section 12.2). For agent *i* in state *s*, the step is located at *Q*^*i,s*^–i.e., the Q-value averaged over actions. When the agent is optimistic (*resp*. pessimistic), a higher weight is put on the part of the istribution which is above (*resp*. below) that step.

Considering a Q-value distribution for agent *i* taking action *a*^*i*^ in state *s*, 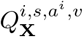, we compute the mean of this distribution, averaged over agent *i*’s strategy 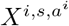:

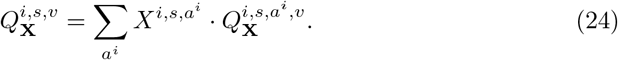

We define the weights tensor 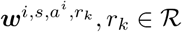, as follows. For all agent *i*, state *s*, action *a*^*i*^, and neuron *r*_*k*_,

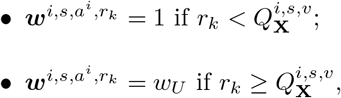

where *w*_*U*_ *<* 1 if the agent is pessimistic, and *w*_*U*_ *>* 1 if it is optimistic.

We obtain the weighted estimate 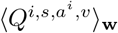 of distribution 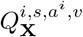 by computing the product of the distribution with the weights vector, normalizing this product by a normalization factor 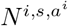, and computing the mean 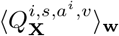 of the resulting distribution:

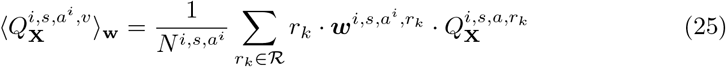

The normalization factor, 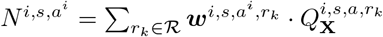, helps us make sure the integral of the weighted distribution over ℝ is 1.

## 3 Results

To validate our framework and highlight its usefulness for uncovering new insights, we test our model across three different choice environments: an individual exploration-exploitation challenge, a social coordination task, and an intertemporal risky choice problem.

### 3.1 Exploration-exploitation challenge

Here, we use our model to confirm previous findings showing that the influence of optimism on performance in a two-armed bandit task is modulated by resource scarcity. The two-armed bandit is a standard task to study the challenge between exploration and exploitation in cognitive psychology. An agent must choose between two options, *a*_1_ and *a*_2_. *a*_1_ gives a reward +1 with probability *p*_1_, and a punishment −1 with probability 1 − *p*_1_; *a*_2_ gives a reward +1 with probability *p*_2_, and a punishment −1 with probability 1 − *p*_2_. By repeatedly choosing between the two options, the agent must learn which is most rewarding on average (Fig. 2A). We choose a SARSA learner here because it explicitly accounts for exploratory behavior.

**Figure 2:**
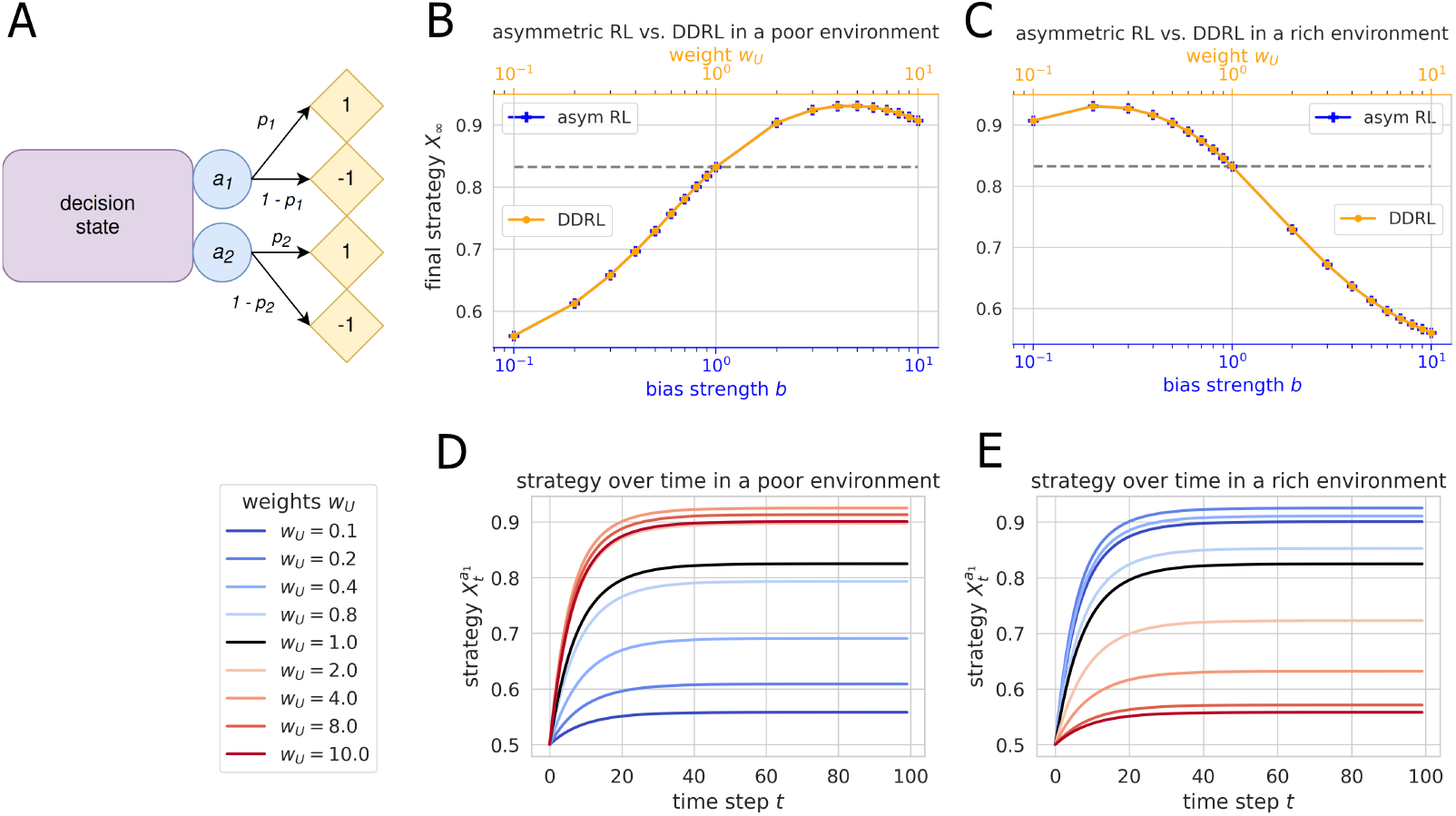
Influence of optimism/pessimism on a two-armed bandit environment with varying resource scarcity. **A** The two-armed bandit environment. **B**–**C** Comparison between final strategies *X*_*∞*_ obtained *via* DDRL (yellow curve) vs. asymmetric RL (blue curve), **B** on a “poor” two-armed bandit task (*p*_1_ = 0.3, *p*_2_ = 0.1); **C** on a “rich” two-armed bandit task (*p*_1_ = 0.9, *p*_2_ = 0.7). These final strategies vary as a function of bias strength 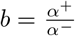. For DDRL agents, the final strategies were obtained from a trajectory of 150 time steps, after checking that this number of time steps was sufficient for convergence (see Fig 2**D**–**E**). For asymmetric agents, the final strategies were obtained analytically, using the formula in [56]. **D**–**E** Strategy plots of a DDRL agent performing a two-armed bandit task with various levels of optimism/pessimism, **D** in a poor environment (*p*_1_ = 0.3, *p*_2_ = 0.1); **E** in a rich environment (*p*_1_ = 0.9, *p*_2_ = 0.7). These plots represent the evolution of an agent’s probability to choose the best arm over time. Blue lines indicate pessimistic agents (*w*_*U*_ *<* 1), while red lines indicate optimistic agents (*w*_*U*_ *>* 1). Neutral agents (*w*_*U*_ = 1) are represented by a black line. In all panels, all DDRL agents had *n* = 29 neurons, and a discretization parameter *σ* = 0.0001 to minimize errors (see SI-Part IV).

In previous research using two-armed bandit tasks, it has been established that optimism (implemented as an asymmetric pair of learning rates) increases or decreases an agent’s performance depending on resource scarcity. Optimism is beneficial for performance in scarce environments, and detrimental in abundant environments [56]. Therefore, we tested our framework on a “poor” two-armed bandit, in which the arms’ reward probabilities are *p*_1_ = 0.3 and *p*_2_ = 0.1, and on a “rich” version of the same task, in which the arms’ reward probabilities are *p*_1_ = 0.9 and *p*_2_ = 0.7.

First, we find that DDRL closely approximates the behavior of an asymmetric-RL agent on a two-armed bandit task (Fig. 2B–C). To do so, we checked that both algorithms converged towards the same final strategy *X*_*∞*_–i.e., an agent’s probability to choose the best arm after convergence. We computed final strategies for different bias strengths. In asymmetric RL, bias strength is defined as 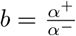 (where *α*^+^ updates positive temporal-difference errors and *α*^−^, negative temporal-difference errors). An analytical calculation of *X*_*∞*_ as a function of *b* is given in [56]. In DDRL, bias strength is modulated by the upper weight, *w*_*U*_ . Here, we simulated the deterministic trajectory of an agent’s strategy *X* over time until convergence for different values of *w*_*U*_ .

Second, we replicate the aforementioned findings on the influence of optimism/ pessimism in poor/rich environments (Fig. 2B–C). In both scarcity conditions, DDRL closely approximates asymmetric RL. More precisely, the final strategy *X*_*∞*_ of a DDRL agent with weight *w*_*U*_ matches the final strategy of an asymmetric-RL agent with bias strength *b* = *w*_*U*_ . We give a formal proof of this equivalence in SI-Section 12.2. Fig. 2B–C shows how the influence of optimism/pessimism on performance is modulated by resource scarcity. In a poor environment (Fig. 2B), optimistic agents (*w*_*U*_ *>* 1) converge to a higher probability of choosing the best arm than neutral (*w*_*U*_ = 1) and pessimistic (*w*_*U*_ *<* 1) agents. In a rich environment (Fig. 2C), by contrast, pessimistic agents converge to a higher probability of choosing the best arm than neutral (*w*_*U*_ = 1) and optimistic (*w*_*U*_ *>* 1) agents. Furthermore, both curves indicate the presence of an optimal bias strength (or upper-weight value), for which the final strategy *X*_*∞*_ is maximal. In our poor environment, this optimal upper-weight value lies close to *w*_*U*_ = 5; whereas, in our rich environment, it is close to *w*_*U*_ = 0.2.

This interaction between optimism/pessimism and resource scarcity, previously established in Ref. [56], is further visualized in Fig. 2D–E. These strategy plots depict the evolution of a DDRL agent’s strategy–i.e., its probability of choosing the best arm–over time for different levels of optimism/pessimism. In a poor environment (Fig 2D), starting from a uniform random strategy (*X* = 0.5), optimistic DDRL agents (*w*_*U*_ *>* 1, red lines) converge towards a higher probability of choosing the best arm than the neutral agent (*w*_*U*_ = 1, black line); whereas pessimistic DDRL agents (*w*_*U*_ *<* 1, blue lines) converge towards lower performance. In line with previous findings [56] and the results mentioned above, this pattern is reversed in a rich environment (Fig. 2E).

Overall, our results confirm that DDRL closely approximates asymmetric RL on a two-armed bandit task with varying resource scarcity. In particular, we replicate previous findings indicating that optimism is beneficial in poor environments, while pessimism enhances performance on the rich version of the task. This means DDRL is a suitable model for optimism/pessimism in this basic choice task. Next, we turn to an environment which is widely used in biology, economics, and sustainability science: the stag-hunt game.

### 3.2 Social coordination task

Here, we use our model to confirm previous findings that show optimism improves coordination in the Stag-Hunt game. The Stag-Hunt game is an iconic coordination problem in the space of social dilemmas [57]. A social dilemma is a situation in which agents individually have an incentive to deviate from outcomes that would be best for the group. Each agent must choose between two actions: cooperate (*c*) or defect (*d*). In the two-agent case, if both cooperate, they get a payoff *R* (called “reward”). If both defect, they get a payoff *P* (called “punishment”). If one cooperates and the other defects, the cooperator receives a payoff of *S* (called “sucker”) and the defector gets a payoff of *T* (called “temptation”) (Fig 3A). In the Stag-Hunt game, the payoff ordering is as follows: *R > T > P > S*. Therefore, both mutual defection and mutual cooperation are Nash equilibria. However, if both agents defect, they receive the sucker’s payoff *S*, which is lower than the maximum reward payoff *R* obtainable by mutual cooperation. This means individual interests align with collective interests, but the best outcome can only be achieved through coordination among agents.

**Figure 3:**
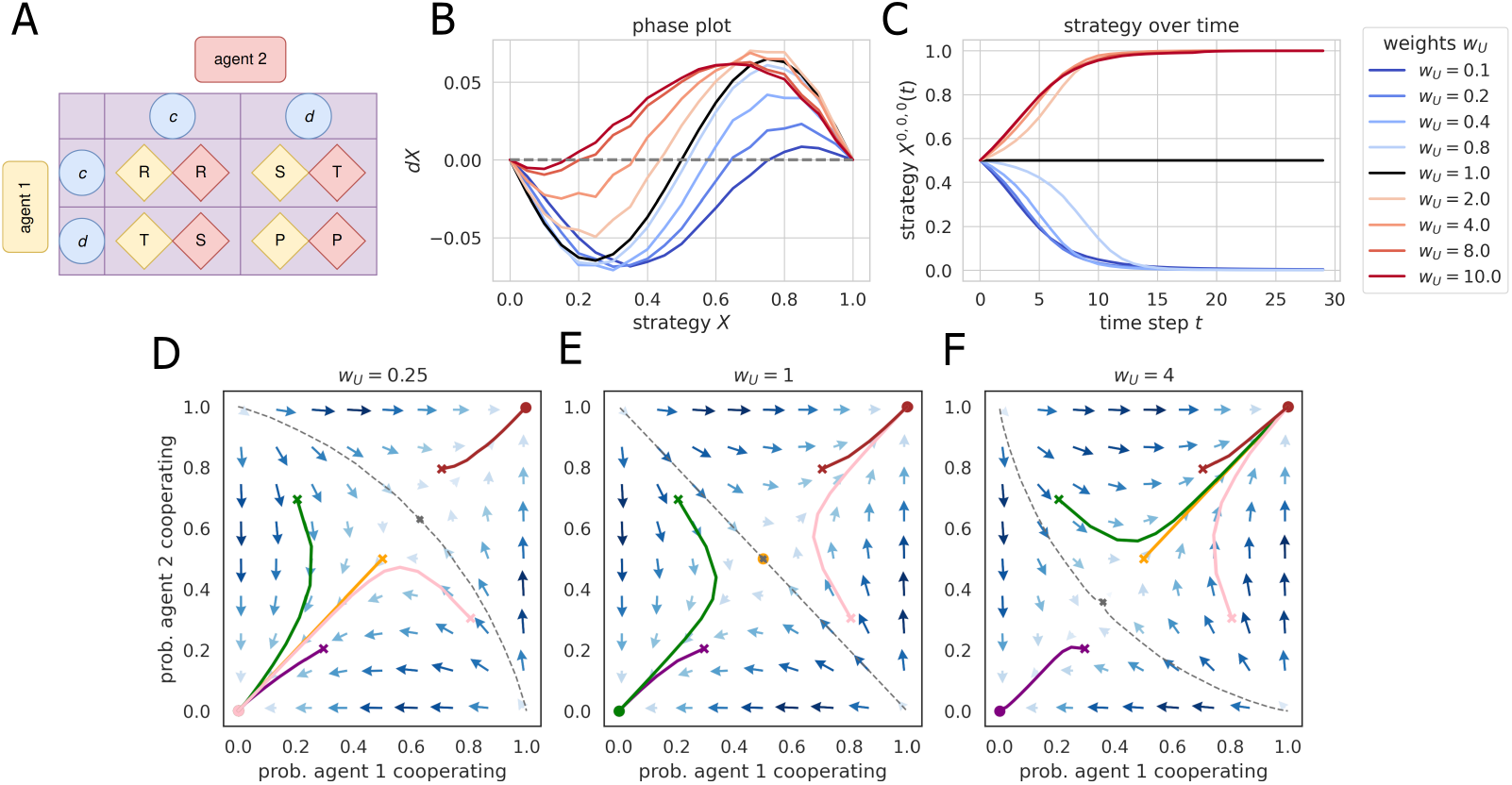
Influence of optimism/pessimism on a Stag-Hunt game. **A** Structure of a social dilemma. In the Stag-Hunt game, *R > T > P > S*. For our analyses, we set *R* = 3, *T* = 1, *P* = 0 and *S* = −2. **B** Phase plot and **C** strategy plot of an agent playing the Stag-Hunt game for various levels of optimism/pessimism. Since the Stag-Hunt game is symmetric among agents, we assumed the second agent had the same strategy as the first. On the phase plot (**B**), *X* = 1 indicates maximal cooperation, and *X* = 0, maximal defection. Intersections of the curves with the *x*-axis indicate stable and unstable fixed points, depending on the sign of the derivative *dX*. The strategy plot (**C**) represents the evolution of an agent’s probability to cooperate over time, given an initial cooperation probability of 0.5. Blue lines represent pessimistic agents (*w*_*U*_ *<* 1), while red lines represent optimistic agents (*w*_*U*_ *>* 1). Neutral agents (*w*_*U*_ = 1) are represented by a black line. **D**–**F** Flow plots of two agents playing the stag-hunt game **D** with a pessimistic heuristic (*w*_*U*_ = 0.25); **E** with no heuristic (“neutral”, *w*_*U*_ = 1); **F** with an optimistic heuristic (*w*_*U*_ = 4). Arrows represent the direction and intensity of the strategies’ derivative *dX*. Colored lines are sample trajectories starting from various initial conditions (+), while round dots indicate fixed points. Dotted gray lines delineate the flow plots’ separatrices–i.e., the threshold of initial cooperation probability above which agents learn towards maximal cooperation, and under which they learn towards maximal defection. In all panels, all DDRL agents had *n* = 29 neurons, and a discretization parameter *σ* = 0.08 to minimize errors (see SI-Part IV).

Previous research in multi-agent reinforcement learning suggests that variants of optimism enhance coordination on various tasks. For instance, Lenient Learning, an algorithm that initially has agents dismiss negative outcomes, has led to performance improvements across multiple coordination games, such as the climb and penalty games [32]. Likewise, Hysteretic Q-learning (a multi-agent equivalent of asymmetric RL) has been shown to improve coordination on several tasks: the aforementioned climb and penalty games, a collaborative ball balancing task, and a pursuit game (where predators need to coordinate to capture prey) [25]. In this section, we investigate whether optimism enhances coordination in the Stag-Hunt game. The climb and penalty games are two three-action versions of the Stag-Hunt problem, which received its name from a story in which hunters need to coordinate to capture a stag, analogous to predators coordinating to capture prey. Thus, the Stag-Hunt game is well-suited to capture the essence of these coordination problems.

In line with previous findings, we find that optimistic DDRL agents converge to mutual cooperation when starting from a uniformly random strategy. In contrast, pessimistic agents converge to the suboptimal equilibrium of mutual defection. A neutral agent remains at its mixed strategy for the payoff values of our Stag-Hunt game (Fig. 3C). Fig. 3B shows that the uniform random strategy (i.e., a cooperation and defection probability of 0.5) is an unstable fixed point of the learning dynamics for neutral agents. Therefore, if a neutral agent’s initial strategy is higher than 0.5 (and provided the second agent’s initial strategy is the same), it converges toward cooperation. If, by contrast, both agents’ initial strategies are below 0.5, they converge toward defection. Optimism (red lines in Fig. 3B) lowers this threshold–i.e., two optimistic DDRL agents may converge towards cooperation even if they started with a strategy *X <* 0.5–while pessimism heightens it. This is why, in Fig. 3C, when all agents start with a uniformly random strategy, optimistic agents end up mutually cooperating, whereas pessimistic agents converge towards mutual defection. Thus, optimism facilitates cooperation, while pessimism hinders it.

The flow plots in Figs. 3D–F give an overview of the learning dynamics in the two-dimensional phase space spanned by each agent’s individual strategy. Thus, the phase plots in Fig 3B are equivalent to the diagonal in the flow plots in Fig 3D–F, for the relevant optimism levels. The strategy changes *dX* are represented by the flow arrows. The separatrix curves separating the two basins of attraction of the two equilibria, mutual cooperation and mutual defection, are shown with dashed gray lines. Optimism distorts these separatrices. When agents are pessimistic, it takes higher initial levels of cooperation to eventually learn to fully cooperate (Fig. 3D). When they are optimistic, learning to cooperate fully is possible even from initially lower levels of cooperation (Fig. 3F).

Additionally, we tested our model on a prisoner’s dilemma with *T > R > P > S*. Here, agents have the incentive to defect, regardless of what the other agent does. Thus, mutual cooperation is no longer an equilibrium despite being the optimal solution for the collective. We find that optimism/pessimism does not affect the learning dynamics. In other terms, whatever the upper-weight value *w*_*U*_, DDRL agents always converge towards mutual defection in this environment (see SI-Section 13.3). In this dilemma, indeed, the reward *R* is lower than the temptation *T*. Therefore, if an agent is optimistic that the other will cooperate, it is still incentivized to defect, since it will then obtain *T* instead of *R*. The mutual-defection equilibrium thus cannot be changed through the optimism/pessimism heuristic. In a stag-hunt game, by contrast, if an agent is optimistic that the other agent will cooperate, it increases its incentive to cooperate as well (see SI-Section 13.4 for an analytical explanation of this difference between the two games).

In short, our results confirm that optimism enhances coordination in DDRL agents, while pessimism hinders it. These results echo previous findings across several multi-agent reinforcement learning algorithms in coordination games. Additionally, our flow plot visualization provides further insights into the collective learning dynamics in coordination games and into how the optimistic/pessimistic heuristic distorts this landscape. Next, we consider a lesser-known intertemporal risky choice problem.

### 3.3 Intertemporal risky choice problem

Here, we use our model to confirm previous findings that show optimism increases risk-taking. We also uncover new findings in this lesser-known choice environment [58, 59], as it incorporates a novel component of decision-making into our analysis: intertemporal choice. In the choice environment, an agent must trade off a low but safe reward with a high reward with potentially negative future consequences. This is implemented via two environmental states: a prosperous and a degraded one. The agent must choose between a risky action and a safe action. In the prosperous state, taking the risky action leads to a reward *r*_*r*_ = 1, whereas the safe action leads to a reward *r*_*s*_ = 0.5. However, choosing the risky action in the prosperous state results in a transition to the degraded state with probability *p*_*c*_ = 0.2. There, the agent gets no reward *r*_*d*_ = 0, and can only transition back to the prosperous state with probability *p*_*r*_ = 0.1 if it chooses the safe action (Fig. 4A). To simplify the analysis, we use an Actor-Critic agent here, since it has no explicit exploration term (see Section 2.1.2). We presume that a SARSA agent would exhibit additional, exploration-related effects and interactions.

**Figure 4:**
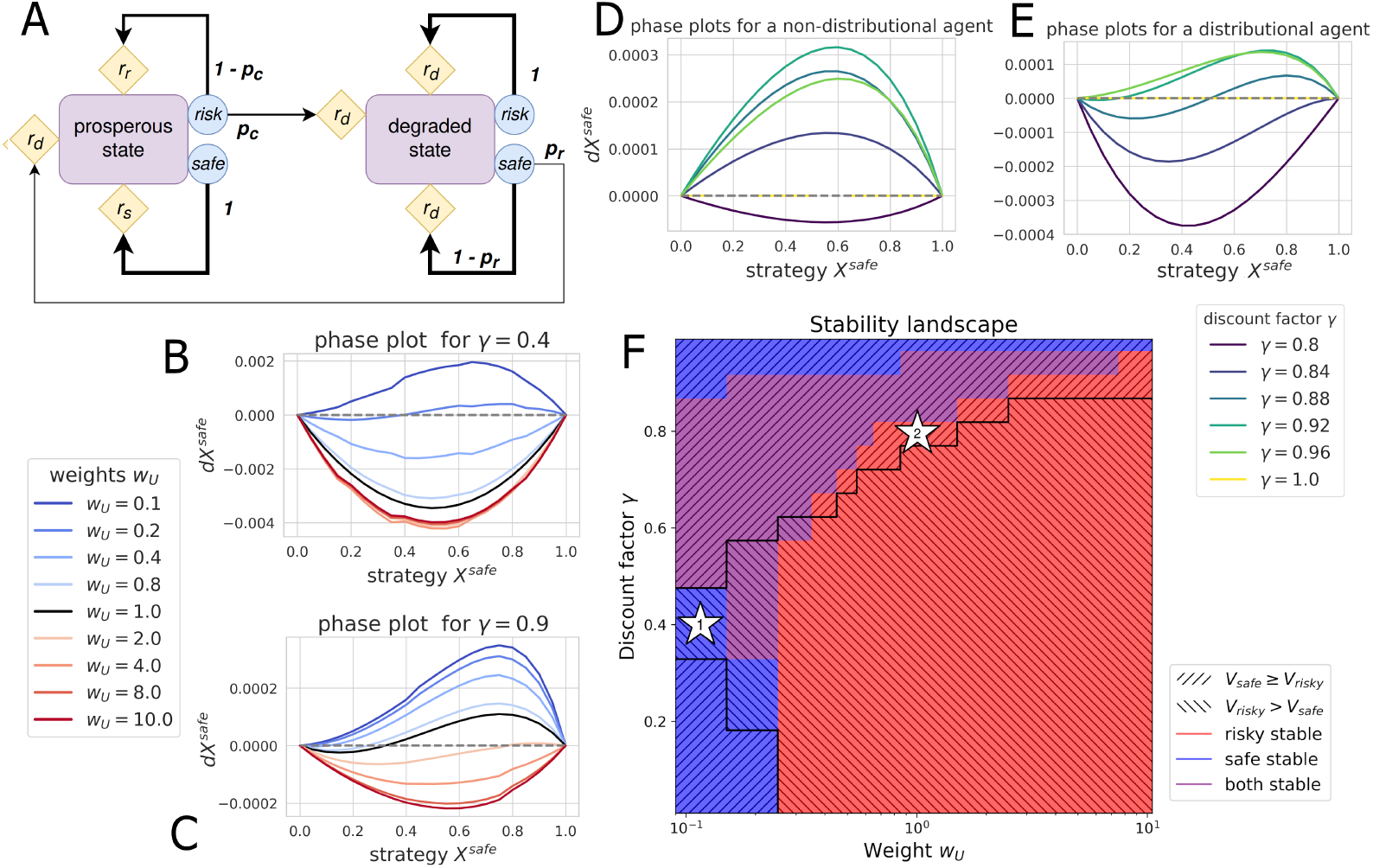
Influence of optimism/pessimism in a risk-reward dilemma. **A** Structure of a risk-reward dilemma. In our analyses, we set the collapse probability *p*_*c*_ = 0.2 and the recovery probability *p*_*r*_ = 0.1; we fixed the risky reward *r*_*r*_ = 1, the safe reward *r*_*s*_ = 0.5, and the degraded reward *r*_*d*_ = 0. **B**–**C** Phase plots of a DDRL agent performing a risk-reward dilemma with various levels of optimism/pessimism, **B** with a low discount factor (*γ* = 0.4); **C** with a high discount factor (*γ* = 0.9). Blue lines indicate pessimistic agents (*w*_*U*_ *<* 1), while red lines indicate optimistic agents (*w*_*U*_ *>* 1). Neutral agents (*w*_*U*_ = 1) are represented by a black line. **D**–**E** Phase plots of an agent performing a risk-reward dilemma with various discount factors, **D** using a regular deterministic Actor-Critic algorithm, **E** using a neutral (*w*_*U*_ = 1) DDRL Actor-Critic algorithm. On all phase plots (**B**–**E**), *X* = 1 corresponds to maximal safety, and *X* = 0, to maximal risk. Intersections with the *x*-axis indicate stable and unstable fixed points (depending on the sign of the derivative *dX*). **F** Stability landscape of a DDRL on a risk-reward dilemma for various combinations of weights *w*_*U*_ and discount factors *γ*. Red patches correspond to regions in the parameter space where the risky policy (*X*^*prosp,risk*^ = 1) is stable, while blue patches indicate that the safe policy (*X*^*prosp,safe*^) is stable. Purple patches correspond to regions in the parameter space where both policies are stable. Stars mark regions of interest, where the safe or risky policy has highest value but is unstable. In all panels, the agent had *n* = 29 neurons, and a discretization parameter *σ* = 0.05 to minimize errors (see SI-Part IV).

Previous research on risk-sensitivity in RL agents uses asymmetric learning rates to implement different attitudes towards risk, where optimism corresponds to risk-seeking and pessimism to risk-aversion [60, 61]. Thus, we hypothesize that in our intertemporal risky choice problem, an optimistic DDRL agent is more likely to choose the risky action than a neutral agent. We also hypothesize that this pattern is reversed for pessimistic DDRL agents.

We confirm both hypotheses. Optimism increases risky behavior while pessimism favors safe behavior (Figs. 4B–C). Figs. 4B–C show the strategy change *dX*^prosp, safe^ versus the strategy *X*^prosp, safe^ to choose the safe action in the prosperous environmental state for different levels of optimism and pessimism and two discount factors. In the degraded state, only the safe action leads to a recovery to the rewarding prosperous state. Thus, we assumed *X*^deg,safe^ = 1 for Figs. 4B–C. Flow plots of the entire strategy space for given discount factors and weight values can be found in SI-Fig. 14.6 and SI-Fig. 14.7.

When the agent cares little about future rewards (low discount factor *γ* = 0.4, Fig. 4B), only intense pessimism can push an agent to act safely. Only the most pessimistic agents (*w*_*U*_ = 0.1 and *w*_*U*_ = 0.2) have a chance to converge towards complete safety in the prosperous state (*X*^prosp,safe^ = 1). All the other agents, including the moderately pessimistic and neutral ones, converge towards maximal risk-taking (*X*^prosp,safe^ = 0). This means that, in such situations, to act safely, intense pessimism can compensate for an agent’s lack of concern for future rewards.

When the agent cares more about future rewards (high discount factor *γ* = 0.9, Fig. 4C), the more optimistic an agent is, the more likely it is to converge toward the risky strategy. In fact, for a moderately-to-strongly optimistic agent (*w*_*U*_ ≥ 4), the risky strategy is the only stable one. A mildly optimistic agent (*w*_*U*_ = 2) could converge to a maximally safe approach, but only if its initial safety levels are very high. A reversed pattern is found for the different levels of pessimism. Because *γ* is high, risk-seeking is overall lower than with a low discount factor.

In both cases, we observe instances of bistability in the strategy space. It depends on the initial strategy to which strategy the agent converges. This is particularly interesting as non-distributional agents (using expected values from a continuous reward spectrum) do not exhibit any bistable regimes (Fig. 4D). This observation aligns with the finding that an optimal strategy in a Markov Decision Process is always deterministic [62]. Fig. 4E compares our distributional agent dynamics without any bias (*w*_*u*_ = 1) for different discount factors. We find certain discount factors *γ* (e.g., *γ* = 0.88, *γ* = 0.92) at which the learning dynamics are bistable. Thus, bistability does not solely emerge from optimism or pessimism, but from an intrinsic feature of our DDRL framework: the discretization of the reward signal. Fig. 4F visualizes how the stability of the safe and risky strategies depends on the interaction of bias weight values *w*_*U*_ and discount factors *γ*. The red zone denotes the stability of the risky strategy (*X*^prosp, risky^ = 1); the blue zone, that of the safe one (*X*^prosp, safe^ = 1); whereas, in the purple zone, both strategies are stable. Thus, bistability emerges for a broad range of weights and discount factors, including for neutral agents (*w*_*U*_ = 1). Overall, we find that these bistable regimes depend on the interplay among the discount factor, the optimism weight, and the reward discretization.

We also find a kind of incoherent choice pattern, which we coin “individual dilemma”. Just as in a social dilemma, an agent would be better off playing the strategy that yields the highest value, but it cannot realize this, as it is pulled towards the other, suboptimal strategy, which is stable (Stars in Fig. 4F). To see this, we highlighted the region in parameter space where the safe policy yields a higher value to the agent than the risky policy (*V*_safe_ *> V*_risky_) and likewise, *V*_risky_ *> V*_safe_. We define 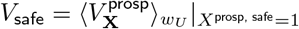 and 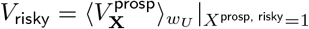. For most parameter values, when a strategy yields a higher overall value, it is also stable. Nevertheless, Fig. 4F exhibits some deviations from this pattern. For example, at *w*_*U*_ = 0.1 and *γ* = 0.402, the safe strategy is stable while the risky strategy yields a higher value (Star 1 in Fig. 4F). At *w*_*U*_ = 1 and *γ* = 0.794, the risky strategy is stable, while the safe strategy yields a higher value (Star 1 in Fig. 4F). Although the state value of the safe strategy is higher than that of the risky one, the state-action value of the risky action is higher than that of the safe action. In SI-Fig. 14.10, we show the corresponding agent’s state-value and state-action-value distributions.

Overall, our results confirm previous findings regarding the relationship between optimism, pessimism, and risky behavior. Optimism enhances risky behavior, whereas pessimism favors risk-averse behavior. Additionally, our framework leads to new, intriguing findings: the emergence of bistable regimes and individual dilemmas.

## 4 Discussion

In this work, we derived the deterministic dynamics of distributional multi-agent reinforcement learning to obtain a unifying, biologically plausible, and tractable framework for modeling individual cognition and behavior across contexts. We applied our framework to model the widely studied optimism heuristic. We established the soundness of our model by reproducing results across three iconic choice domains: an exploration-exploitation challenge, a social coordination task, and a risky choice problem. This demonstrates the potential of our model as a unifying framework to study the optimism-pessimism heuristic in collective learning dynamics.

Our framework rests on a simple, biologically plausible mechanism in which outcome distributions are discretized by a set of neurons. Empirical research has recently provided evidence for an analogous computation of distributions during learning and decision-making in the brain [42, 63]. Thus, our reward discretization should not be viewed simply as a technical trick, but as a meaningful biological feature.

In this regard, our framework could help enhance understanding of how reward information is encoded neurologically. How many neurons are involved in categorizing environmental rewards, and how does this number impact the accuracy of this distribution? In the Supplementary Information, we show how discretization sensitivity, *σ*, affects the error between the expected values of the continuous and discretized reward spectra across various environments (see SI-Part IV). In particular, we found that, for a fixed number of neurons *n*, the sensitivity *σ* that minimizes this error is environment-dependent. Thus, is there a context-dependent mechanism that modulates reward sensitivity? Recent studies suggest such a context-dependent mechanism. It has been found that humans do not encode objective outcomes but rather evaluate options with respect to available alternatives [64]. This has been modeled with RL as range normalization [65], in which the value of an option is rescaled by the difference between its maximum and minimum values in the environment. Our framework could incorporate such a mechanism to explore how varying assumptions about neurological reward representations affect outcomes, leading to empirically testable hypotheses.

Furthermore, our framework could help uncover the empirical basis for optimistic reinforcement learning in the brain. In particular, it is an open question whether optimism in the brain manifests at the level of state-action values or temporal-difference errors. Indeed, asymmetric RL, which is widely used in psychological experiments, locates optimism/pessimism at the level of temporal-difference errors. These are updated differentially through asymmetric learning rates. By contrast, distributional reinforcement learning distorts the state-action return distribution by applying weights. According to prior work, these implementations should produce the same results in stateless environments, but they are expected to yield different outcomes in stateful environments [37]. Thus, comparing our framework with asymmetric RL algorithms in stateful choice environments could generate empirically testable hypotheses that would distinguish between signatures of state-action values versus temporal-difference-error optimism in the brain.

Applying our model to a single-agent intertemporal risky-choice task, we found that reward discretization can lead to a bistable choice regime. In other words, strategies become path-dependent. By contrast, non-distributional agents that compute expected values of a continuous reward spectrum do not exhibit bistable regimes. This finding suggests a novel explanation for the adaptive heuristic of choice hysteresis [66]. Choice hysteresis (also called “perseveration”) is the tendency to repeat past choices in a process akin to habit formation. Thus, it implies path-dependence in an agent’s choice history [26]. Hysteresis in intertemporal choice has been documented from eye-tracking studies [67] to studies about addictive behaviors [68]. It has been shown that hysteresis in sequential decision-making increases in aging rats [69], and hysteresis has even been accounted for in terms of attractor models [70]. In experimental psychology, there is a debate about whether optimism can be reduced to perseveration. Several studies and meta-analyses have striven to disentangle optimism from perseveration in empirical datasets [26, 71]. They concluded that both heuristics were present and irreducible to one another–but it was also shown that the optimism heuristic leads to higher performance in a wide range of two-armed bandit tasks, compared to perseveration [72]. Our framework suggests that, rather than an adaptive heuristic, choice hysteresis is a mere by-product of a more fundamental biological mechanism. We hypothesize that perseveration arises from a lower-level biological feature: reward discretization via a finite set of neurons. Overall, our model provides a unifying framework for investigating the interaction between optimism and choice hysteresis in sequential decision-making.

Our model also gives rise to a novel explanation for an incoherent choice phenomenon, which we termed individual dilemma. In analogy with the well-known social dilemma, an agent would be better off choosing the strategy that provides the highest value, but cannot do so because it is drawn toward the other, suboptimal strategy that is stable. Our results show that this situation arises in a wide range of combinations between discount factors and optimism weights. While individual dilemmas might appear as flaws in our model, there are many cases where human choices are inconsistent. Perhaps one of the most compelling illustrations lies in compulsive behavior. Compulsion occurs when one keeps engaging in actions that feel immediately relieving but are harmful in the long run. It is a key component of, e.g., Tourette’s syndrome and obsessive-compulsive disorder (OCD), and can occur even when patients are completely aware of the counterproductivity of their behavior. On the one hand, because of the ritualistic nature of compulsions, it has been suggested that habit plays a central role in compulsivity; on the other hand, there is a significant overlap between compulsivity and impulsivity–i.e., the reduced ability to delay rewards, which is quantified by the discount factor *γ* [73, 74]. Our findings suggest there could be a complex interplay between reward discounting, optimism/pessimism, and discretization/habit in the emergence of compulsive behavior. Indeed, individual dilemmas appear for certain values of the discount factor and optimism weight. Thus, our framework could help disentangle the contributions of impulsivity, optimism and habit formation in giving rise to incoherent choice patterns. Moreover, our results show that an individual dilemma can be escaped by tuning up the optimism weight or the discount factor, depending on the region of the parameter space one finds oneself in. This opens the door to developing treatments which, by acting on impulsivity and/or optimism levels, could alleviate compulsive symptoms.

Besides empirical questions, our model could be applied to study the role of optimism/pessimism in complex collective-action problems, such as climate change. For tractability reasons, cognitive, social, and environmental factors are often studied in isolation, missing an integrated perspective [47]. Our model can contribute to such an integrated perspective. For example, it could enable insights into the influence of optimism/pessimism on cooperation in the face of climate tipping points [75]. Previous work found that the severity of the tipping impact, the certainty of the tipping threshold, and caring for future rewards are conducive to cooperation [76, 77, 78]. But what if these factors are not enough? Is it better to be optimistic, to induce coordination in the social dilemma, as we have seen in our social coordination task? Or is it better to be pessimistic and risk-averse, to avoid triggering the tipping element, as we have seen in our intertemporal risky-choice problem? Our model could answer these questions, clarify how the effects of optimism and pessimism interact, and identify the conditions under which optimism or pessimism is conducive to cooperation.

Because of its distributional basis, our framework opens the door to representing decision-making heuristics as distortions on state-action return distributions. Among the multitude of referenced heuristics, self-efficacy [79] and confirmation bias [80] could be interesting candidates to model in individual and collective scenarios. This would help address outstanding debates and questions in several disciplines. For instance, it has been suggested that optimism could be a confirmation bias in disguise, at the individual [80, 27] and collective levels [44]. Thus, distinguishing between optimism and confirmation heuristics in individual and collective settings is an essential task for experimental researchers. In this regard, our framework could be used to generate hypotheses and predictions.

We also highlight some limitations of our framework. First, although it builds upon stochastic games, which are the foundation of multi-agent reinforcement learning in machine learning [81], our framework does not scale to large environments (such as large grid worlds or 3D computer games). Such choice environments have large state and reward spaces, leading to a combinatorial explosion and the curse of dimensionality. The analysis of collective dynamics in such high-dimensional environments requires the use of more suitable methods. Second, our framework presupposes categorically discretized neurons. This limits the precision of approximating reward distributions, compared to more sophisticated parameter-estimation methods used in the machine-learning literature [82, 55]. Thus, our framework’s application to machine learning might be limited, as it is not an optimization method. It is a biologically inspired modeling framework designed to improve understanding.

In conclusion, our deterministic distributional multi-agent reinforcement learning framework bridges the gap between the microscopic constraints of neural reward encoding and the macroscopic dynamics of collective behavior. It offers a unified perspective through which seemingly disparate phenomena–from optimism and choice hysteresis to compulsive behavior and social coordination–can be understood as emergent properties of a common underlying mechanism. While we acknowledge the trade-off between tractability and computational scalability, our approach provides a necessary middle ground for cross-disciplinary inquiry across the animal, human, and machine behavioral sciences. Moving forward, this framework offers a robust toolkit for generating testable hypotheses and investigating how individual-level cognitive biases shape collective outcomes in complex real-world challenges, ultimately advancing our understanding of the intricate interplay between neural computation, individual choice, and collective dynamics.

