## Supplementary Information for "Deterministic dynamics of distributional multi-agent reinforcement learning"

### Deterministic Distributional Reinforcement Learning

Clemence Bergerot, Wolfram Barfuss

December 21, 2025

**References** **143**

### 1 Introduction

Deterministic Distributional Reinforcement Learning (DDRL) combines deterministic approaches to RL (Barfuss et al., 2019) with distributional RL (Bellemare et al., 2017). Within this framework, the learning dynamics of multi-agent games can be modeled and visualized as deterministic trajectories. DDRL also enables the implementation of cognitive heuristics (such as optimism/pessimism) as distortions of the value distributions. Therefore, it is particularly suited to studying the impact of cognitive heuristics on collective learning dynamics in a wide range of scenarios.

This Supplementary Information is structured as follows. In **?@sec-agents** , we introduce various classes of DDRL agents, as well as different ways to implement optimism/pessimism. Then, in **?@sec-environments** , we describe the environments we use to obtain results in our manuscript. **?@sec-utilities** contains all required functions to plot the figures. In **?@sec-calibration** , we carry out error tests to calibrate the agents' sensitivity to rewards in different environments. This enables the computation of all results from the manuscript, as well as supplementary results, in **?@sec-results** .

#### 1.1 Reproducibility

The present package and supplementary information were created thanks to nbdev and quarto. This ensures all the simulations are fully reproducible.

The package and notebooks associated with this documentation can be found on this Github repo. There, you can download the package and run the notebooks in order to reproduce our simulations.

To do this, clone the repository:

```
git clone https://github.com/clembergerot/pyDDRL
cd pyDDRL
```

Then set up the virtual environment:

```
python -m venv venv
source venv/bin/activate
```

...and install the package in editable mode:

```
pip install -e .
```

Then you should be able to execute all the notebooks.

#### 1.2 How to use

Plot a simple flowplot:

```python
import numpy as np
import matplotlib.pyplot as plt
import seaborn as sns

# Environment
from pyDDRL.Environments.RiskRewardDilemma import RiskRewardDilemma

# Agent
from pyDDRL.Agents.DDRLBase import DDRLBase
from pyDDRL.Agents.DDRLActorCritic import DDRLActorCritic

# Utilities
from pyCRLD.Utils.Helpers import *
from pyCRLD.Utils import FlowPlot as fp
from pyDDRL.Utils.PlottingTools import *

# Define environment
env = RiskRewardDilemma(pc=0.1, pr=0.2, rc=0.5, rr=1, rd=0)

# Define agent
aei = DDRLActorCritic(env, nr_reward_bins=29, discretization_sigma=0.05,
                      learning_rates=0.1, discount_factors=0.9,
                      Rmin=0, Rmax=1,
                      method="mean", use_prefactor=True)

# Plot
x = ([0], [0], [0])  # prosperous state
y = ([0], [1], [0])  # degraded state

fig, ax = plt.subplots(1, 1, figsize=(3,3))
fp.plot_strategy_flow(aei, x, y, flowarrow_points = np.linspace(0.01 ,0.99, 9),
                      NrRandom=16, axes=ax)
ax.set_title("Flowplot in a risk-reward dilemma")
ax.set_ylabel("prob. safe in deg. state")
ax.set_xlabel("prob. safe in prosp. state")
```

```
WARNING:2025-12-21 19:04:54,681:jax._src.xla_bridge:794: An NVIDIA GPU may be present on this [illegible]
enabled jaxlib is not installed. Falling back to cpu.
```

```
Text(0.5, 0, 'prob. safe in prosp. state')
```

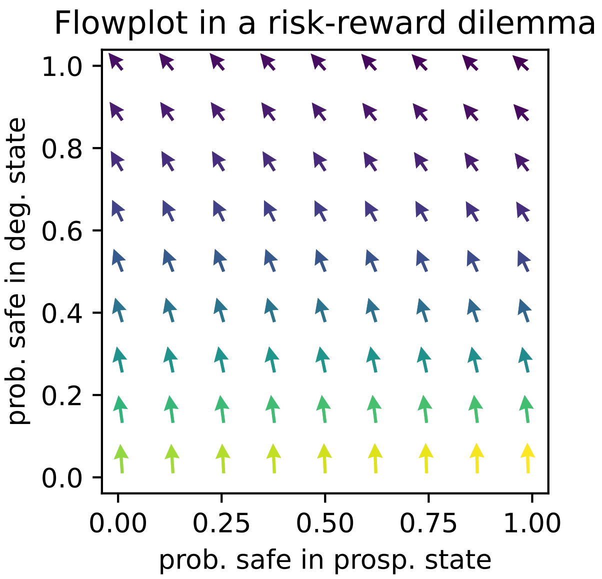

Flowplot in a risk-reward dilemma
prob. safe in deg. state
prob. safe in prosp. state
0.0
0.2
0.4
0.6
0.8
1.0
0.00
0.25
0.50
0.75
1.00

### Part I

### Agents

In this section, we introduce various agent classes as well as biasing methods for the implementation of heuristics. Chapter 2 contains the base functions for computing reward and value distributions. In Chapter 3 , we introduce the functions that are specific to the SARSA algorithm of RL. By contrast, in Chapter 4 , we introduce those that are specific to the Actor-Critic version. Finally, Chapter 5 describes three different ways of implementing the optimism/pessimism heuristic: the quantile method, the expectile method, and the weights method.

### 2 DDRL Base

In this section, we provide the base functions for all DDRL agents.

First, we import everything necessary:

```
#| default_exp Agents.DDRLBase
```

```
#| hide
# Imports for the nbdev development environment
import nbdev
from nbdev.showdoc import *
from fastcore.basics import patch
```

```
#| export
import numpy as np
import scipy.stats as stats

import jax.numpy as jnp
from jax import jit
from functools import partial

import itertools as it

# pyCRLD imports
from pyCRLD.Agents.Base import abase
from pyCRLD.Utils.Helpers import *
from pyCRLD.Utils.Helpers import compute_stationarydistribution
```

```
# imports for this notebook
import matplotlib.pyplot as plt

# actor-critic agent
from pyCRLD.Agents.StrategyActorCritic import stratAC
```

```
## Environment for testing

# two-armed bandit
from pyDDRL.Environments.TwoArmedBandit import TwoArmedBandit
```

#### 2.1 Test environment

We use a two-armed bandit (defined in Section Chapter 6 ) as a test environment. To illustrate different ways of computing reward distributions, we introduce two types of two-armed bandits: one with discrete return, and one with continuous return.

##### 2.1.1 Stochastic two-armed bandit with discrete return

In this version, an agent can choose between two options: * the first arm, $a_1$, yields a reward $+1$ with probability 0.7, and a punishment $-1$ with probability 0.3; * the second arm, $a_2$, yields a reward $+1$ with probability 0.3, and a punishment $-1$ with probability 0.7.

Let's implement this environment:

```
# reward amounts
xs = np.array([[-1, 1],  # first arm
               [-1, 1]])  # second arm

# probabilities
ps = np.array([[0.3, 0.7],  # first arm
               [0.7, 0.3]])  # second arm

# Define environment
tab_1s = TwoArmedBandit(disttype='discrete', xs=xs, ps=ps)
tab_1s.Rdict
```

```
{'x': array([[[[[-1.,  1.]],

          [[-1.,  1.]]]]]),
 'p': array([[[[[0.3, 0.7]],

          [[0.7, 0.3]]]]])}
```

This environment can be visualized as follows:

```
## Plot the return distributions

x = tab_1s.Rdict['x']
p = tab_1s.Rdict['p']

# Plot
fig, axes = plt.subplots(1, 2, figsize=(4.5, 1.5))
for a, ax in enumerate(axes):
    ax.spines[['right', 'top', 'left']].set_visible(False)
    ax.bar(x[0,0,a][0], p[0,0,a][0], width=0.3, color='k')
    ax.set_xlabel("r")
    ax.set_ylim([0,0.8])
    ax.set_title("random return for arm "+str(a+1), fontsize=10)
axes[0].set_ylabel(r'$p(R = r)$')
axes[1].set_yticklabels([])
axes[1].set_yticks([])
plt.show()
```

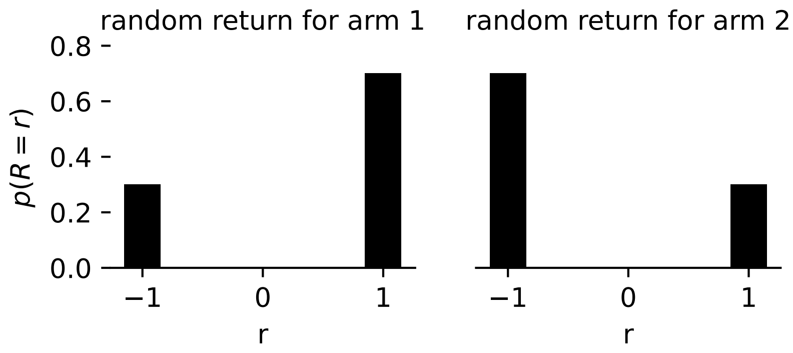

##### 2.1.2 Stochastic two-armed bandit with continuous return

In a stochastic two-armed bandit with continuous return, the random return follows a continuous distribution.

Let's consider the following example, in which an agent can choose between two arms: * the first arm gives a random return that follows a Gaussian distribution, centered around $\mu = 1$, with standard deviation $\sigma = 0.5$; * the second arm gives a random return that follows a Gaussian distribution, centered around $\mu = -1$, with standard deviation $\sigma = 2.0$.

```
# Define environment

distA = ('norm', 1, 0.5)  # first arm
distB = ('norm', -1, 2)  # second arm

tab_cr = TwoArmedBandit(distA=distA, distB=distB,
                        disttype="continuous")
tab_cr
```

```
TwoArmedBandit_norm_1_norm_-1
```

Now let's plot the distribution of the random return for each arm.

```
## Plot the return distributions

x = np.linspace(-10, 10)
norm1 = stats.norm.pdf(x, 1, 0.5)  # first arm
norm2 = stats.norm.pdf(x, -1, 2)  # second arm
norms = [norm1, norm2]

# Plot
fig, axes = plt.subplots(1, 2, figsize=(5, 1.5))
for s_, ax in enumerate(axes):
    ax.spines[['right', 'top', 'left']].set_visible(False)
    ax.plot(x, norms[s_], color='blue')
    ax.set_xlabel("r")
    ax.set_ylim([0,0.8])
    ax.set_title("random return for arm "+str(s_+1), fontsize=10)
axes[0].set_ylabel(r'$p(R = r)$')
axes[1].set_yticklabels([])
```

```
axes[1].set_yticks([])
plt.show()
```

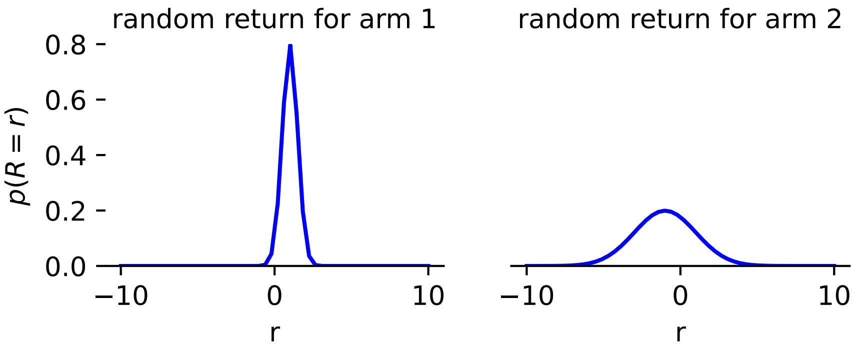

#### 2.2 Discretization and basic reward distribution $R^{i,s,a,\acute{s},r}$

##### 2.2.1 Discretization grid

We assume an agent is equipped with a number $n$ of neurons, such that each neuron $i \in [1, n]$ is sensitive to an amount of reward $r_i$, with $r_1$ the minimum amount of reward perceived and $r_n$ the maximum amount, and for all $i \in [1, n]$ $r_i = r_1 + (i-1)\frac{r_n - r_1}{n-1}$.

The discretization grid is the set $r_i, i \in [1, n]$ of perceived amounts of rewards (also called "grid points"). We call "width" the difference $\Delta r = (i-1)\frac{r_n - r_1}{n-1}$ between two consecutive grid points.

The discretization grid will be the support of all subsequent distributions.

```
#| export
def compute_discretization_grid(Rmin, # Minimum reward perceived
                                Rmax, # Maximum reward perceived
                                Nr # Number of gridpoints
                                ) -> tuple: # Gridpoints, Width
    """
    Compute discretization grid (rewards neurons are sensitive to)
    for base DDRL agent.
    """
    gridpoints = np.linspace(Rmin, Rmax, Nr)
    width  = np.round(gridpoints[1] - gridpoints[0], 8)
    return gridpoints, width
```

##### 2.2.2 Basic reward distribution $R^{i,s,a,\acute{s},r}$

Let us now consider a stochastic game with reward tensor $R$ and transition tensor $T$. For all $r \in (r_j)_{j\in[1,n]}$, we define the reward distribution $R^{i,s,a,\acute{s},r}$ for agent $i$, current state $s$, joint action $a$, and next state $\acute{s}$, as:

$$R^{i,s,a,\acute{s},r} = \int_{r-\frac{\Delta x}{2}}^{r+\frac{\Delta x}{2}} \frac{1}{\sqrt{2\pi\sigma^2}} \exp[-\frac{(y - R^{i,s,a,\acute{s}})^2}{2\sigma^2}]dy$$

Note that each $R^{i,s,a,\acute{s}}$ is an entry of the reward tensor $R$. In other terms, for all $r \in (r_j)_{j\in[1,n]}$, $R^{i,s,a,\acute{s},r}$ is the area under the bell curve centered in $R^{i,s,a,\acute{s}}$, taken between $r - \frac{\Delta r}{2}$ and $r + \frac{\Delta r}{2}$.

For environments with stochastic returns (such as in the stochastic two-armed bandit task), the entries of the reward tensor are themselves distributions.

If these distributions are continuous, we define $F^{i,s,a,\acute{s}}(r)$ the cumulative distribution function of continuous reward distribution ${}^cR^{i,s,a,\acute{s}}(r)$. Then, we discretize as follows:

$$R^{i,s,a,\acute{s},r} = \int_{r-\frac{\Delta x}{2}}^{r+\frac{\Delta x}{2}} F^{i,s,a,\acute{s}}(y)dy$$

We define a dispatcher function, which associates keywords with corresponding distribution shapes, so that we can compute relevant cumulative distribution functions:

```
#| export
def dispatcher():
    dispatcher={'norm':stats.norm, 'alpha':stats.alpha, 'beta':stats.beta,
                'powerlaw':stats.powerlaw}
    return dispatcher
```

Now we can compute the basic reward distribution for our two cases: - stochastic returns following discrete distributions. - stochastic returns following continuous distributions.

```
#| export
def compute_Risjasr(env, # The env object
                    Rmin,  # minimum perceived reward
                    Rmax,  # maximum perceived reward
                    nr,  # The number of reward bins
                    discretization_sigma # Width of the discritization normal
↪   distribution
                   ) -> np.ndarray: # Reward distribution
    """
    Compute basic reward distribution Risjasr.
    """

    # get the discretization grid
    points, width = compute_discretization_grid(Rmin, Rmax, nr)

    # init the reward distribution tensor
    Risjasr = np.zeros(env.R.shape + (nr,))  # last dim is distrib

    # go through the reward tensor of the environment
    for index, _ in np.ndenumerate(env.R):
        if env.disttype == 'continuous':  # return follows continuous distrib
            sigma = env.Rdict['scale'][index]  # sd
            loc = env.Rdict['loc'][index]  # mean
            dist = env.Rdict['dist'][index]  # distribution shape
            # choose the right distribution shape
            disp = dispatcher()
            func = disp[dist]
            # compute discretized distribution
```

```
            for r in range(nr):
                if r==0: # first interval
                    prop = func.cdf(points[r]+width/2, loc, sigma)
                elif r==nr-1: # last interval
                    prop = 1 - func.cdf(points[r]-width/2, loc, sigma)
                else: # all other intervals
                    prop = func.cdf(points[r]+width/2, loc, sigma)\
                        - func.cdf(points[r]-width/2, loc, sigma)
                Risjasr[index +(r,)] = prop

        elif env.disttype == None:  # return is deterministic
            loc = env.R[index]  # mean
            sigma = discretization_sigma  # sigma
            func = stats.norm
            # compute discretized distribution
            for r in range(nr):
                if r==0: # first interval
                    prop = func.cdf(points[r]+width/2, loc, sigma)
                elif r==nr-1: # last interval
                    prop = 1 - func.cdf(points[r]-width/2, loc, sigma)
                else: # all other intervals
                    prop = func.cdf(points[r]+width/2, loc, sigma)\
                        - func.cdf(points[r]-width/2, loc, sigma)
                Risjasr[index +(r,)] = prop

        else:  # return follows discrete distribution
            x = env.Rdict['x'][index]  # values for which prob is non zero
            p = env.Rdict['p'][index]  # prob that return equals those values
            sigma = discretization_sigma  # discretization width
            func = stats.norm
            # compute discretized distribution
            for i in range(len(x)):
                for r in range(nr):
                    if r==0: # first interval
                        prop = p[i] * func.cdf(points[r]+width/2, x[i], sigma)
                    elif r==nr-1: # last interval
                        prop = p[i] * (1 - func.cdf(points[r]-width/2, x[i], sigma))
                    else: # all other intervals
                        prop = p[i] * (func.cdf(points[r]+width/2, x[i], sigma)\
                                       - func.cdf(points[r]-width/2, x[i], sigma))
                    Risjasr[index +(r,)] += prop

    return Risjasr
```

Here are the discretized distributions for the two different two-armed bandits:

1. Stochastic two-armed bandit with discrete return

```
env = tab_1s
Rmin = -1.5
Rmax = 1.5
```

```
nr = 25
sigma = 0.1

gridpoints, width = compute_discretization_grid(Rmin, Rmax, nr)
Rr = compute_Risjasr(env, Rmin, Rmax, nr, sigma)
```

```
# Plot reward distribution for both arms
fig, axes = plt.subplots(1, 2, figsize=(8,3))
for a, ax in enumerate(axes):
    ax.spines[['right', 'top', 'left']].set_visible(False)
    ax.bar(gridpoints, Rr[0,0,a,0,:], width=0.9*width, color='k')
    rm = np.dot(gridpoints, Rr[0,0,a,0,:])
    ax.plot([rm, rm], [0.0, 0.05], c='red', lw=5)
    ax.set_xlabel("r")
    ax.set_ylabel(r"$R^{i,s,a,\acute s, r}$")
    ax.set_title("Reward distribution for arm "+str(a+1))
    ax.set_ylim([0, np.max(Rr[:,:,a,:])+0.01])
plt.show()
```

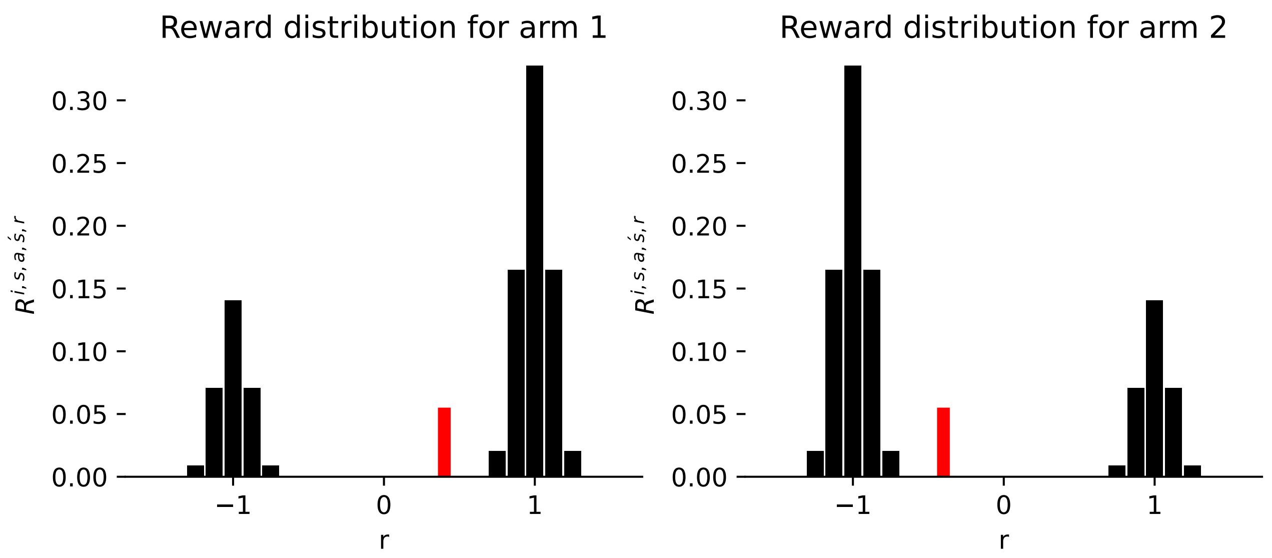

2. Stochastic two-armed bandit with continuous return

```
env = tab_cr
Rmin = -10
Rmax = 10
nr = 29
sigma = 0.1

gridpoints, width = compute_discretization_grid(Rmin, Rmax, nr)
Rr = compute_Risjasr(env, Rmin, Rmax, nr, sigma)
```

```
# Plot reward distribution for both arms
fig, axes = plt.subplots(1, 2, figsize=(8,3))
for a, ax in enumerate(axes):
    ax.spines[['right', 'top', 'left']].set_visible(False)
```

```
    ax.bar(gridpoints, Rr[0,0,a,0,:], width=0.9*width, color='k')
    rm = np.dot(gridpoints, Rr[0,0,a,0,:])
    ax.plot([rm, rm], [0.0, 0.05], c='red', lw=5)
    ax.set_xlabel("r")
    ax.set_ylabel(r"$R^{i,s,a,\acute s, r}$")
    ax.set_title("Reward distribution for arm "+str(a+1))
    ax.set_ylim([0, 0.5])
plt.show()
```

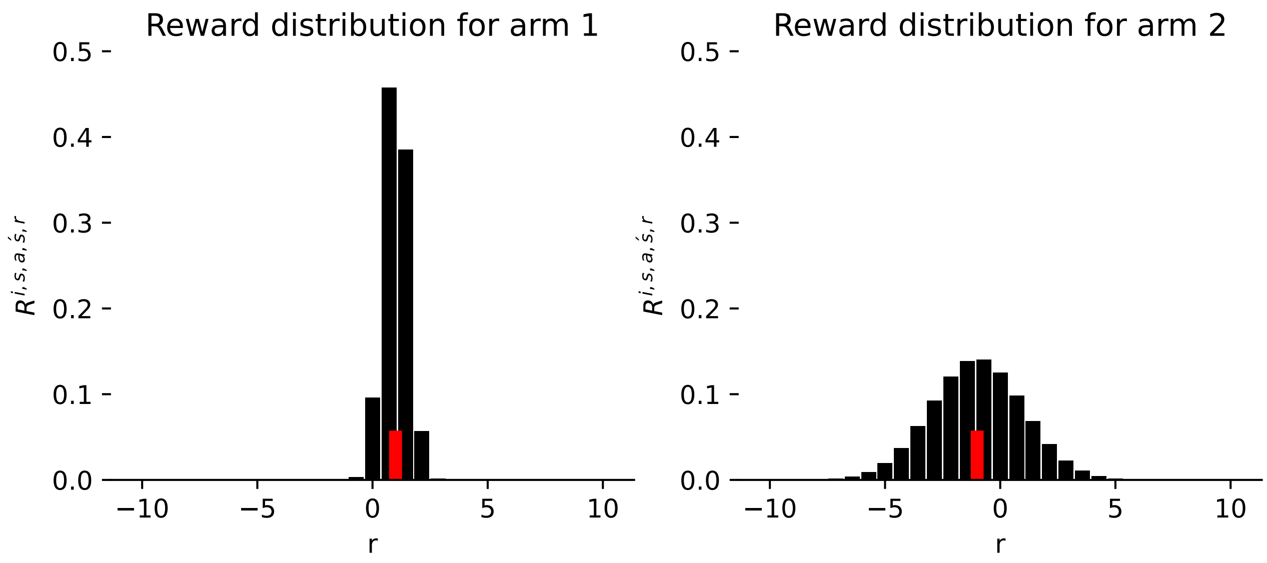

#### 2.3 Smoothed Bellman Kronecker delta $\delta^{i,v,r,\acute{v}}$

In expected-value RL, the value of a state $s$ for agent $i$ is computed via the Bellman equation:

$$V^{i,s} = (1 - \gamma^i)R^{i,s} + \gamma^i V^{i,\acute{s}}$$

where $V^{i,\acute{s}}$ is the value of the next state, $\acute{s}$.

In distributional form, we want to compute the probability that (discretized) values $r$, $v$ and $\acute{v}$ satisfy the Bellman equation $v = (1 - \gamma^i)r + \gamma^i \acute{v}$. To this end, we introduce the smoothed Bellman Kronecker delta, which also allows for some uncertainty about the categorization into different bins. This uncertainty is also modulated by the discretization standard deviation, $\sigma$. Calling $val(r, \acute{v})$ the expression $(1 - \gamma^i)r + \gamma^i \acute{v}$,

$$\delta^{i,v,r,\acute{v}} = \int_{v-\frac{w}{2}}^{v+\frac{w}{2}} \frac{1}{\sqrt{2\pi\sigma^2}} \exp[-\frac{[y - val(r, \acute{v})]^2}{2\sigma^2}]dy$$

The following function computes the Bellman Kronecker delta for given environment, minimum and maximum perceived rewards, number of neurons, and discount factors:

```
 #| export
# Smoothed Bellman Kronecker delta for Bellman equation
def compute_b_ivrv_(env, # env object
                    Rmin,
```

```
                    Rmax,
                    nr, # number of bins
                    discretization_sigma, # width of discretization normal
↪   distribution
                    gamma,  # discount factor
                    pre  # prefactor
                   ) -> np.ndarray:

    """
    Compute smoothed Bellman Kronecker delta b_ivrv_.
    """

    points, width = compute_discretization_grid(Rmin, Rmax, nr)
    sigma = discretization_sigma
    b_ivrv_ = np.ones((env.N, nr, nr, nr)) * -1

    for v in range(nr):
        for r in range(nr):
            for v_ in range(nr):

                valv = pre * points[r] + gamma * points[v_]

                # gaussian distribution around valq with std `sig`
                if v==0:
                    prop = stats.norm.cdf(points[v]+width/2, valv, sigma)
                elif v==nr-1: # last intervall
                    prop = 1 - stats.norm.cdf(points[v]-width/2, valv, sigma)
                else:
                    prop = stats.norm.cdf(points[v]+width/2,
                                          valv, sigma) -
↪   stats.norm.cdf(points[v]-width/2, valv, sigma)

                b_ivrv_[:,v,r,v_] = prop
    return b_ivrv_
```

#### 2.4 DDRL Base class

Now we can define the DDRL Base class:

```
#| export
class DDRLBase(abase):
    """
    Base class for Distributional Deterministic Reinforcement Learning
    in strategy space.
    """

    def __init__(self,
                 env, # An environment object
                 nr_reward_bins=8, # The number of bins to resolve rewards
                 discretization_sigma=0.1, # standard div for reward discretization
                 learning_rates=0.1, # agents' learning rates
```

```
                 discount_factors=0.0, # agents' discount factors
                 Rmin=None,  # minimum perceived reward
                 Rmax=None,  # maximum perceived reward
                 method='mean',  # biasing method
                 tau=0.5,  # tau for expectile and quantile methods
                 wU=1.,  # upper weight for weights method
                 choice_intensities=1.0, # agents' choice intensities
                 opteinsum=True,  # optimize einsum functions
                 use_prefactor=False,  # scale values with rewards
                 **kwargs):

        self.env = env
        Tt = env.T; assert np.allclose(Tt.sum(-1), 1)
        Rt = env.R
        super().__init__(Tt, Rt, discount_factors, True, opteinsum)
        self.F = jnp.array(env.F)

        # learning rates
        self.alpha = make_variable_vector(learning_rates, self.N)

        # intensity of choice
        self.beta = make_variable_vector(choice_intensities, self.N)

        # discount factors
        self.gamma = make_variable_vector(discount_factors, self.N)

        # pre-factor
        self.pre = 1 - self.gamma if use_prefactor else jnp.ones(self.N)

        # number of reward bins
        self.nr = nr_reward_bins

        # reward discretization deviation
        self.discsig = discretization_sigma

        # Rmin and Rmax
        if Rmin==None:
            self.Rmin = env.R.min()
        else:
            self.Rmin = Rmin

        if Rmax==None:
            self.Rmax = env.R.max()
        else:
            self.Rmax = Rmax

        # discretization grid
        disc_grid = compute_discretization_grid(self.Rmin, self.Rmax, self.nr)
        self.gridpoints = disc_grid[0]
        self.gridwidth = disc_grid[1]

        # reward tensor distribution
        self.Rr = compute_Risjasr(env, self.Rmin, self.Rmax, self.nr, self.discsig)
```

```
    # Bellman ivrv_
    self.b_ivrv_ = compute_b_ivrv_(env, self.Rmin, self.Rmax, self.nr,
     ↪  self.discsig, self.gamma, self.pre)

    # method
    self.method = method

    # value for quantile or expectile
    self.tau = tau

    # weights
    self.wU = wU
```

We consider the following agent-environment interface: * 1 agent with learning rate $\alpha = 0.1$, discount factor $\gamma = 0.9$, choice intensity $\beta = 4$; * playing the aforementioned stochastic two-armed bandit task with continuous return.

```
aei = DDRLBase(tab_cr,  # environment
               nr_reward_bins=29,
               discretization_sigma=0.1,
               learning_rates=0.1,
               discount_factors=0.9,
               choice_intensities=4.,
               Rmin=-10,
               Rmax=10)
```

```
WARNING:2025-12-21 19:05:37,158:jax._src.xla_bridge:794: An NVIDIA GPU may be present on this 
enabled jaxlib is not installed. Falling back to cpu.
```

#### 2.5 Computing distributions in DDRL

Now we can compute and plot all the distributions we need.

##### 2.5.1 Reward distributions $R^{i,s,a^i,r}$ and $R^{i,s,r}$

The basic reward distribution can be averaged over joint actions $a$ and next states $\acute{s}$. We call $R^{i,s,a^i,r}$ the reward distribution for agent $i$ taking action $a^i$ in state $s$, averaged over transitions to the next state $\acute{s}$, and over other agents' actions $a^j$:

$$R^{i,s,a^i,r} = \sum_{\acute{s}} \sum_{a^j} \prod_{j} X^{j,s,a^j} T^{s,a^i a^j,\acute{s}} R^{i,s,a^i a^j,\acute{s},r}$$

```
 #| export
@partial(jit, static_argnums=0)
def Risar(self:DDRLBase,
          Xisa
         ) -> jnp.ndarray:
```

```
    i = 0; a = 1; s = 2; s_ = 3; r = 4  # Variables
    j2k = list(range(5, 5+self.N-1))  # other agents
    b2d = list(range(5+self.N-1, 5+self.N-1 + self.N))  # all actions
    e2f = list(range(4+2*self.N, 4+2*self.N + self.N-1))  # all other acts
    Risjasr = self.Rr

    sumsis = [[j2k[l], s, e2f[l]] for l in range(self.N-1)]  # sum inds
    otherX = list(it.chain(*zip((self.N-1)*[Xisa], sumsis)))

    args = [self.Omega, [i]+j2k+[a]+b2d+e2f] + otherX\
        + [self.T, [s]+b2d+[s_], Risjasr, [i, s]+b2d+[s_, r],
           [i, s, a, r]]
    Risar = jnp.einsum(*args, optimize=self.opti)
    return Risar

DDRLBase.Risar = Risar
```

We assume the agent chooses arm 1 with probability 0.3, and arm 2 with probability 0.7:

```
# agent's strategy
X = np.ones((1,1,2)) * 0.3
X[:,:,1] = 1 - X[:,:,0]
```

Now we can plot the reward distribution $R^{i,s,a^i,r}$ for each arm.

```
Rr = aei.Risar(X)

# Plot reward distribution Risar for both arms
fig, axes = plt.subplots(1, 2, figsize=(8,3))
for a, ax in enumerate(axes):
    ax.spines[['right', 'top', 'left']].set_visible(False)
    ax.bar(gridpoints, Rr[0,0,a,:], width=0.9*width, color='k')
    rm = np.dot(gridpoints, Rr[0,0,a,:])
    ax.plot([rm, rm], [0.0, 0.05], c='red', lw=5)
    ax.set_xlabel("r")
    ax.set_ylabel(r"$R^{i,s,a,r}$")
    ax.set_title("Reward distribution for arm "+str(a+1))
    ax.set_ylim([0, np.max(Rr[0,0,:,:]+0.01)])
plt.show()
```

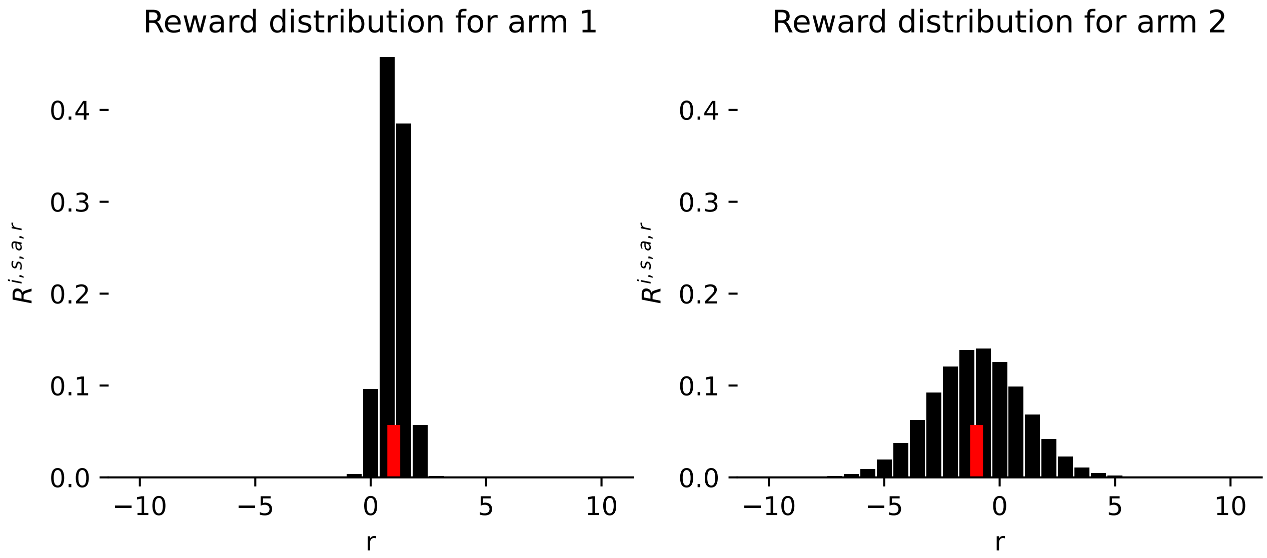

The reward distribution $R^{i,s,r}$ denotes the probability that agent $i$ receives reward $r$ in state $s$. It is averaged over joint actions $a$ (including agent $i$'s) and next states $\acute{s}$:

$$R^{i,s,r} = \sum_{\acute{s}} \sum_{a} \prod_{j} X^{j,s,a} \cdot T^{s,a,\acute{s}} \cdot R^{i,s,a,\acute{s},r}$$

```
 #| export
@partial(jit, static_argnums=0)
def Risr(self:DDRLBase, Xisa):
    """
    Compute reward distribution Risr joint strategy `Xisa`.
    """
    i = 0; s = 1; sprim = 2; r=3; b2d = list(range(4, 4+self.N))
    Risjasr = jnp.array(self.Rr)

    X4einsum = list(it.chain(*zip(Xisa,
                                  [[s, b2d[a]] for a in range(self.N)])))

    args = X4einsum + [self.T, [s]+b2d+[sprim],
                       Risjasr, [i, s]+b2d+[sprim, r], [i, s, r]]

    return jnp.einsum(*args, optimize=self.opti)

DDRLBase.Risr= Risr  # Monkey-patching - possibly problematic?
```

Now plotting this reward distribution for our agent:

```
Rr = aei.Risr(X)

# Plot reward distribution Risr for both arms
fig, ax = plt.subplots(1, 1, figsize=(4,3))
ax.spines[['right', 'top', 'left']].set_visible(False)
ax.bar(gridpoints, Rr[0,0,:], width=0.9*width, color='k')
rm = np.dot(gridpoints, Rr[0,0,:])
ax.plot([rm, rm], [0.0, 0.05], c='red', lw=5)
ax.set_xlabel("r")
```

```
ax.set_ylabel(r"$R^{i,s,r}$")
ax.set_title("Reward distribution averaged over actions")
plt.show()
```

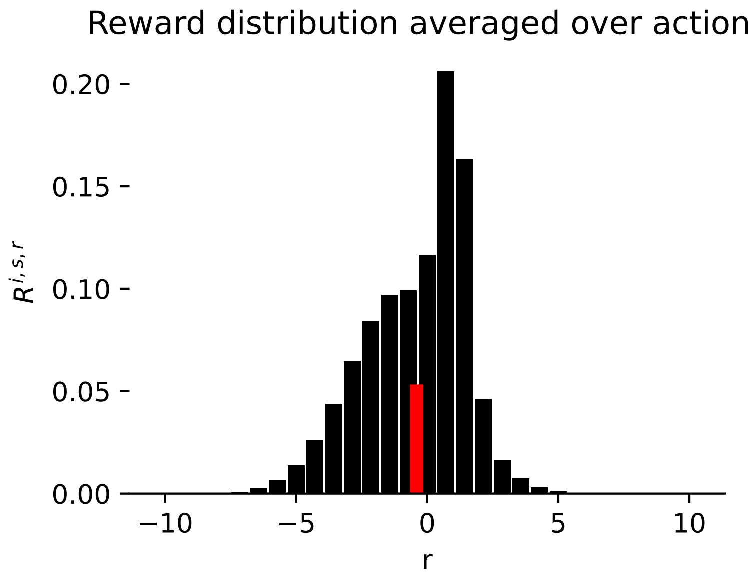

#### 2.6 Value distribution $V^{i,s,v}$ and biased value

We now calculate the value distribution $V^{i,s,v}$. The probability that the random value $V$ equals $v$ for agent $i$ in state $s$ is a conjunction of the following probabilities: * that random reward $R$ for agent $i$ in state $s$ equals $r$; * that there is a transition from state $s$ to state $\acute{s}$; * that $v$, $r$ and $\acute{v}$ conform to the above Bellman equation.

The probability that there is a transition from any state $s$ to any state $\acute{s}$ is given by the environment's stationary distribution, $\bar{T}^{s,\acute{s}}$.

Therefore,

$$V^{i,s,v} = \sum_{r,\acute{v},\acute{s}} \delta^{i,v,r,\acute{v}} R^{i,s,r} \bar{T}^{s,\acute{s}} V^{i,\acute{s},\acute{v}}$$

We now define

$$M^{i,s,v,\acute{s},\acute{v}} = \sum_{r} \delta^{i,v,r,\acute{v}} R^{i,s,r} \bar{T}^{s\acute{s}}$$

Thus,

$$V^{i,s,v} = \sum_{\acute{v},\acute{s}} M^{i,s,v,\acute{s},\acute{v}} V^{i,\acute{s},\acute{v}}$$

We can combine the indices $s \times v = w$ and $\acute{s} \times \acute{v} = \acute{w}$. Now, $V^{i,w} = \sum_{\acute{w}} M^{i,w,\acute{w}} V^{i,\acute{w}}$. This makes it apparent that the return distribution should be found in the eigenvalues of the matrix $M$.

The following function computes the matrix $M$:

```python
#| export
@partial(jit, static_argnums=0)
def M_iww_(self:DDRLBase,
           Xisa  # Behavior profiles
          ) -> jnp.ndarray:
    """
    Compute matrix M_iww_ given joint strategy `Xisa`.
    """
    Tss = self.Tss(Xisa)
    Risr = self.Risr(Xisa)
    b_ivrv_ = self.b_ivrv_

    i=0; v=1; r=2; v_=3; s=4; s_=5

    M = jnp.einsum(b_ivrv_, [i,v,r,v_], Risr, [i, s, r], Tss,
                   [s, s_], [i, s, v_, s_, v], optimize=self.opti)
    jnp.allclose(M.sum((-2,-1)), 1.0)
    return M

DDRLBase.M_iww_ = M_iww_
```

Now, we can compute the value distribution $V^{i,s,v}$:

```python
#| export
@partial(jit, static_argnums=0)
def Visv(self:DDRLBase,
        Xisa  # Behavior profiles
        ) -> jnp.ndarray:
    """
    Compute value distribution Visv given joint strategy `Xisa`.
    """

    M_iww_ = self.M_iww_(Xisa)
    Visv = jnp.ones((self.N, self.Z, self.nr)) * -1

    for i in range(self.N):
        mat = M_iww_[i].reshape(self.Z*self.nr, self.Z*self.nr)
        _pS = compute_stationarydistribution(mat)
        ix = jnp.max(jnp.where(_pS.mean(0)!=-10, jnp.arange(_pS.shape[0]), -1))
        pS = _pS[:, ix]

        Ps = pS.reshape(self.Z, self.nr).sum(-1)
        # assert jnp.allclose(self.Ps(Xisa), Ps, atol=1e-4)

        VIsv = pS.reshape(self.Z, self.nr) / pS.reshape(self.Z, self.nr).sum(-1,
↪   keepdims=True)

        Visv = Visv.at[i, ...].set(VIsv)

        # Visv[i, ...] = VIsv

    return Visv
```

```
DDRLBase.Visv = Visv
```

Here is an example with the above agent-environment interface:

```
Vv = aei.Visv(X)

# Plot reward distribution Risar for both arms
fig, ax = plt.subplots(1, 1, figsize=(4,3))
ax.spines[['right', 'top', 'left']].set_visible(False)
ax.bar(gridpoints, Vv[0,0,:], width=0.9*width, color='k')
rm = np.dot(gridpoints, Vv[0,0,:])
ax.plot([rm, rm], [0.0, 0.01], c='red', lw=5)
ax.set_xlabel("v")
ax.set_ylabel(r"$V^{i,s,v}$")
ax.set_title("Value distribution in state 0")
```

```
Text(0.5, 1.0, 'Value distribution in state 0')
```

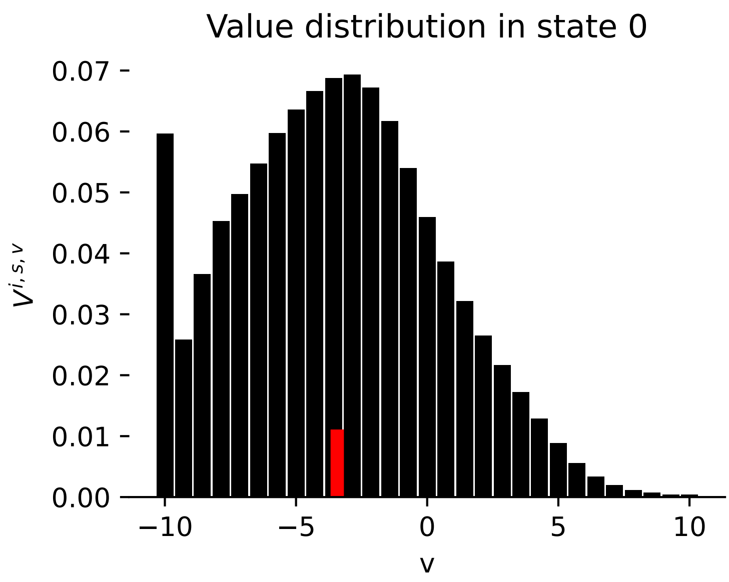

#### 2.7 Additional functions

The functions below help compute the agents' learning trajectories in strategy space. The first one performs a learning step, while the second one performs a reverse step.

```
 #|export
@partial(jit, static_argnums=0)
def step(self:DDRLBase,
         Xisa,  # Behavior profiles
        ) -> jnp.ndarray:
    """
    Performs a learning step along the reward-prediction/temporal-difference error
```

```
    in strategy space, given joint strategy `Xisa`.
    """
    TDe = self.TDerror(Xisa)
    n = jnp.newaxis
    XexpaTDe = Xisa * jnp.exp(self.alpha[:,n,n] * TDe)
    return XexpaTDe / XexpaTDe.sum(-1, keepdims=True), TDe

DDRLBase.step = step
```

```
 #|export
@partial(jit, static_argnums=0)
def reverse_step(self:DDRLBase,
                 Xisa,  # Behavior profiles
                 ) -> jnp.ndarray:
    """
    Performs a reverse learning step along the reward-prediction/temporal-difference
 ↪  error
    in strategy space, given joint strategy `Xisa`.
    """
    TDe = self.TDerror(Xisa)
    n = jnp.newaxis
    XexpaTDe = Xisa * jnp.exp(self.alpha[:,n,n] * -TDe)
    return XexpaTDe / XexpaTDe.sum(-1, keepdims=True), TDe

DDRLBase.reverse_step = reverse_step
```

```
 #| hide
import nbdev; nbdev.nbdev_export()
```

### 3 DDRL SARSA

In this section, we provide functions that are specific to the SARSA version of Distributional Deterministic Reinforcement Learning.

First, we import everything necessary:

```
#| default_exp Agents.DDRLSarsa
```

```
#| hide
# Imports for the nbdev development environment
import nbdev
from nbdev.showdoc import *
from fastcore.basics import patch
```

```
#| export
import numpy as np
from scipy.stats import norm

import jax.numpy as jnp
from jax import jit
from functools import partial

import itertools as it

from pyCRLD.Agents.Base import abase
from pyCRLD.Utils.Helpers import *
from pyCRLD.Utils.Helpers import compute_stationarydistribution

from pyDDRL.Agents.DDRLBase import DDRLBase

from pyDDRL.Agents.Methods import quantile, expectile, weights
```

```
# imports for this notebook
import matplotlib.pyplot as plt
from pyCRLD.Utils import FlowPlot as fp

# test environment
from pyDDRL.Environments.RiskRewardDilemma import RiskRewardDilemma
```

#### 3.1 Test environment

We use a risk-reward dilemma (defined in Chapter 8 ) as a test environment. In this task, the agent can choose between a high-risk, high-gain option, and a low-risk, low-gain option. More precisely, we give our environment the following structure:

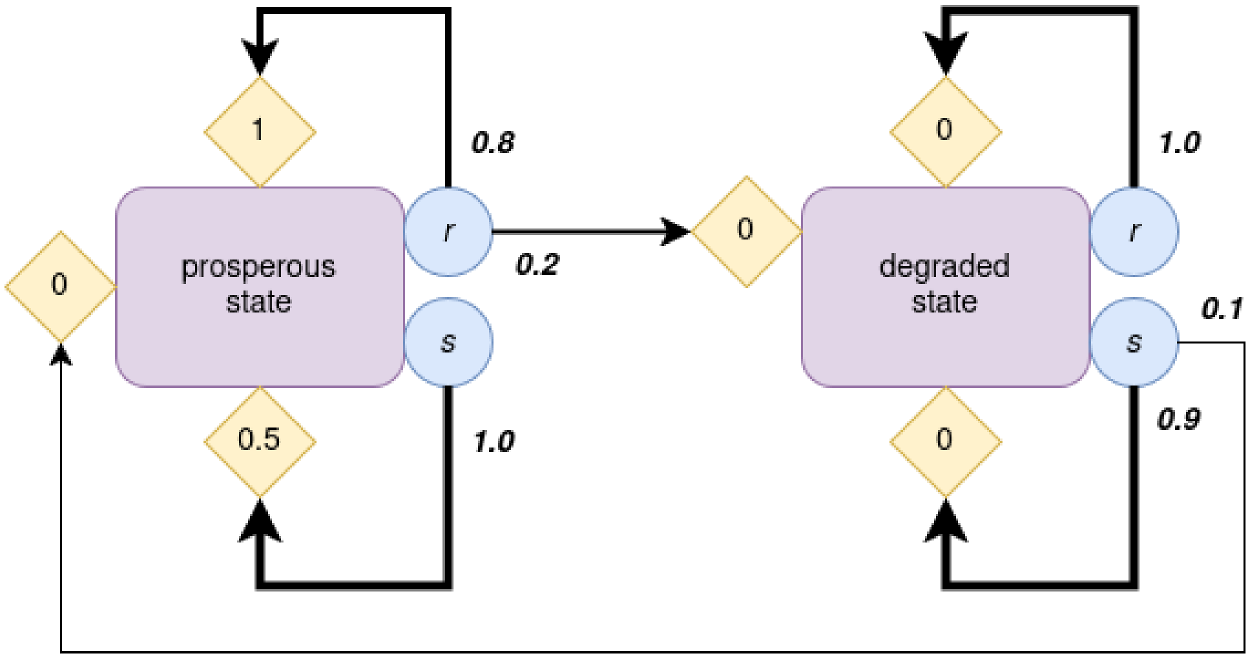

Figure 3.1: A risk-reward dilemma.

If an agent chooses the risky action $r$ in the prosperous state, it gets a higher reward. But there is a probability that it transitions to a degraded state, where it gets no reward. The only way to escape this degraded state is to choose the safe action, $s$.

```
# Define environment
env = RiskRewardDilemma(pc=0.2, pr=0.1, rc=0.5, rr=1, rd=0)
env
```

```
RiskRewardDilemma_0.2_0.1_0.5_1_0
```

#### 3.2 DDRL SARSA class

Now, we can define the DDRL SARSA class:

```
 #| export
class DDRLSarsa(DDRLBase):
    """
    Base class for SARSA version of DDRL in strategy space.
    """

    def __init__(self,
                 env,
                 **kwargs):

        super().__init__(env, **kwargs)
```

We consider the following agent-environment interface: * 1 agent with learning rate $\alpha = 0.1$, discount factor $\gamma = 0.4$, choice intensity $\beta = 50$; * playing the aforementioned risk-reward dilemma.

```python
aei = DDRLSarsa(env,
                nr_reward_bins=29,
                discretization_sigma=0.1,
                learning_rates=0.1,
                choice_intensities=50.,
                discount_factors=0.4,
                Rmin=-0.5,
                Rmax=1.5,
                use_prefactor=True)
```

```
WARNING:2025-12-21 19:06:14,112:jax._src.xla_bridge:794: An NVIDIA GPU may be present on this 
enabled jaxlib is not installed. Falling back to cpu.
```

#### 3.3 Computing distributions in DDRL-SARSA

Now we can compute and plot all the distributions we need.

##### 3.3.1 Q-value distributions $Q^{i,s,a,v}$ and ${}^{next}Q^{i,s,a,\acute{v}}$

In detRL, the next-state Q-value for agent $i$ taking action $a$ in state $s$ is averaged over other agents' actions $a^j$ and next states $\acute{s}$:

$$^{next}Q^{i,s,a} = \sum_{\acute{s}} \sum_{a^j} X^{j,s,a^j} T^{s,a,a^j,\acute{s}} V^{i,\acute{s}}$$

In a distributional framework, therefore,

$$^{next}Q^{i,s,a,\acute{v}} = \sum_{\acute{s}} \sum_{a^j} X^{j,s,a^j} T^{s,a,a^j,\acute{s}} V^{i,\acute{s},\acute{v}}$$

Just like state values, Q-values conform to the Bellman equation. Therefore, we can obtain the Q-value distribution for agent $i$, action $a$ and state $s$, from the next-state Q-value distribution for $(i, a, s)$, the reward distribution for $(i, a, s)$, and the smoothed Bellman Kronecker delta:

$$Q^{i,s,a,v} = \sum_{r,\acute{v}} \delta^{i,v,r,\acute{v}} R^{i,s,a,r} \cdot {}^{next}Q^{i,s,a,\acute{v}}$$

```python
 #| export
@partial(jit, static_argnums=0)
def Qisaq(self:DDRLSarsa,
          Xisa:jnp.ndarray, # Joint strategy
          Tisas:jnp.ndarray=None, # Optional transition for speed-up
          ) -> jnp.ndarray:  # Average state-action values
    """
    Compute Q-value distribution Qisaq, given joint strategy `Xisa`.
    """
    Risar = self.Risar(Xisa)
    b_ivrv_ = self.b_ivrv_
    Visv = self.Visv(Xisa)
    Tisas = self.Tisas(Xisa) if Tisas is None else Tisas
```

```
    i = 0; s = 1; a = 2; s_ = 3; v = 4; v_ = 5; r = 6
    nextQisaq = jnp.einsum(Tisas, [i,s,a,s_], Visv, [i,s_,v], [i,s,a,v],
                           optimize=self.opti)

    Qisaq = jnp.einsum(b_ivrv_, [i,v,r,v_], Risar, [i,s,a,r], nextQisaq, [i,s,a,v_],
 ↪  [i,s,a,v])

    return Qisaq

DDRLSarsa.Qisaq = Qisaq
```

We assume that, in both states, the agent takes the safe action with probability 0.2, and the risky action with probability 0.8.

```
# agent's strategy
X = np.ones((1, 2, 2)) * 0.2
X[:,:,1] = 1 - X[:,:,0]
X
```

```
array([[[0.2, 0.8],
        [0.2, 0.8]]])
```

Now we can plot the Q-value distribution for each action in the prosperous state.

```
Qq = aei.Qisaq(X)

# Plot Q-value distribution Qisaq for both actions
fig, axes = plt.subplots(1, 2, figsize=(8,3))
actions = ["safe", "risky"]
for a, ax in enumerate(axes):
    ax.spines[['right', 'top', 'left']].set_visible(False)
    ax.bar(aei.gridpoints, Qq[0,0,a,:], width=0.9*aei.gridwidth, color='k')
    rm = np.dot(aei.gridpoints, Qq[0,0,a,:])
    ax.plot([rm, rm], [0.0, 0.02], c='red', lw=5)
    ax.set_xlabel("v")
    ax.set_ylabel(r"$Q^{i,s,a,v}$")
    ax.set_title("Q-value distribution for "+str(actions[a])+" action")
    ax.set_ylim([0, np.max(Qq[0,0,:,:]+0.01)])
plt.show()
```

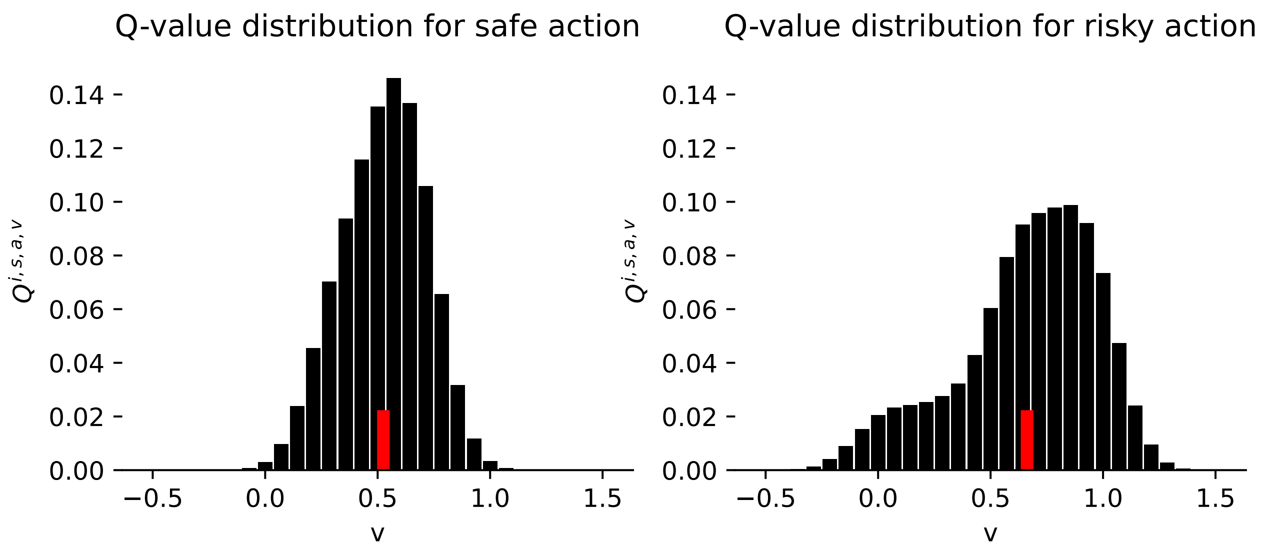

Now we can compute and plot the next-state Q-value distribution:

```
#| export
@partial(jit, static_argnums=0)
def NextQisaq(self:DDRLSarsa,
              Xisa          # Joint strategy
              ) -> jnp.ndarray: # Next values
    """
    Compute next-state Q-value distribution NextQisaq,
    given joint strategy `Xisa`.
    """
    Qisaq = self.Qisaq(Xisa)

    i = 0; a = 1; s = 2; s_ = 3; q = 4
    j2k = list(range(6, 6+self.N-1))  # other agents
    b2d = list(range(6+self.N-1, 6+self.N-1 + self.N))  # all actions
    e2f = list(range(5+2*self.N, 5+2*self.N + self.N-1))  # all other acts

    sumsis = [[j2k[l], s, e2f[l]] for l in range(self.N-1)]  # sum inds
    otherX = list(it.chain(*zip((self.N-1)*[Xisa], sumsis)))

    NextQisq = jnp.einsum(Qisaq, [i, s_, a, q], Xisa, [i, s_, a], [i, s_, q])

    args = [self.Omega, [i]+j2k+[a]+b2d+e2f] + otherX +\
    [self.T, [s]+b2d+[s_], NextQisq, [i, s_, q], [i, s, a, q]]
↪
    return jnp.einsum(*args, optimize=self.opti)

DDRLSarsa.NextQisaq = NextQisaq
```

```
Qq = aei.NextQisaq(X)

# Plot next-state Q-value distribution nextQisaq for both actions
fig, axes = plt.subplots(1, 2, figsize=(8,3))
actions = ["safe", "risky"]
for a, ax in enumerate(axes):
```

```
    ax.spines[['right', 'top', 'left']].set_visible(False)
    ax.bar(aei.gridpoints, Qq[0,0,a,:], width=0.9*aei.gridwidth, color='k')
    rm = np.dot(aei.gridpoints, Qq[0,0,a,:])
    ax.plot([rm, rm], [0.0, 0.02], c='red', lw=5)
    ax.set_xlabel("v")
    ax.set_ylabel(r"${}^{next} Q^{i,s,a,v}$")
    ax.set_title("next-state Q-value distribution for "+str(actions[a])+" action",
                 fontsize=10)
    ax.set_ylim([0, np.max(Qq[0,0,:,:]+0.01)])
plt.show()
```

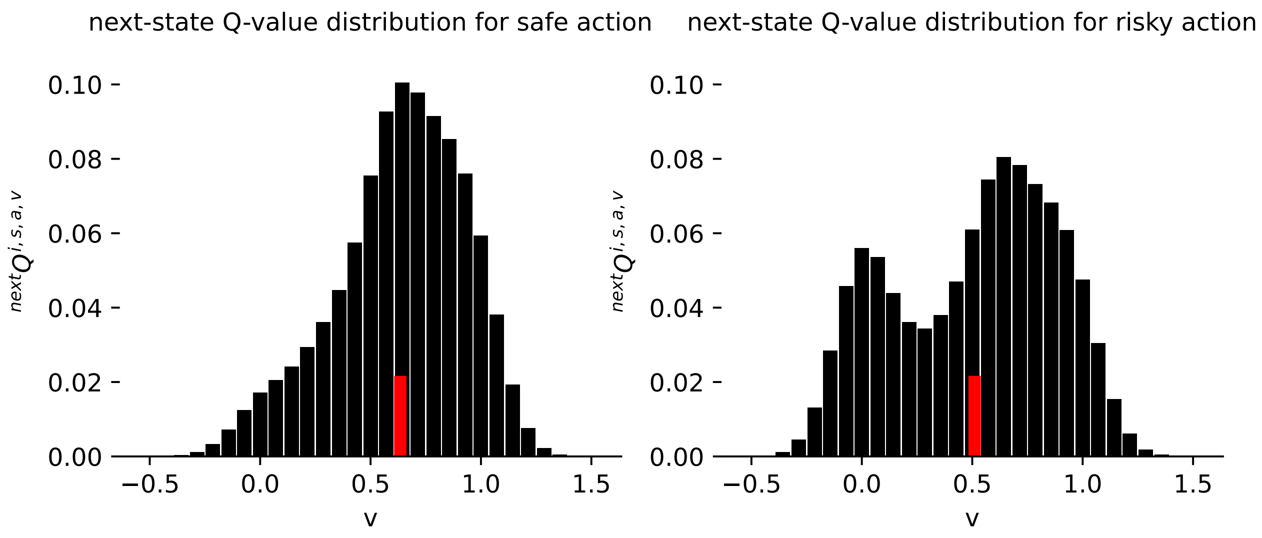

#### 3.4 (Biased) Q-value $Q^{i,s,a}$

A classic estimate of the Q-value is the mean of the Q-value distribution. But a biased agent will get an optimistic or a pessimistic estimate of this distribution. To obtain this biased estimate, we can apply the quantile, expectile, or weights method to the value distribution. To know more about these methods, please refer to Chapter 5 .

Here, we compute the (biased) Q-value from the previous Q-value distribution. When the weights method is used, we take the Q-value $Q^{i,s}$ as a baseline:

```
 #| export
@partial(jit, static_argnums=0)
def Qisa(self,
         Xisa:jnp.ndarray, # Joint strategy
         ) -> jnp.ndarray:  # Average state-action values
    """
    Compute Q-value for state `s` and action `a`.
    """
    Qisaq = self.Qisaq(Xisa)  # Q-value distrib
    gpoints = jnp.ones((self.nr)) * -1
    gpoints = gpoints.at[:].set(self.gridpoints)
    Qisa = jnp.einsum('ijkl,l->ijk', Qisaq, gpoints)
    i = 0; a = 1; s = 2
```

```
    Qis = jnp.einsum(Qisa, [i, s, a], Xisa, [i, s, a], [i, s])  # baseline
    # Baseline: mean of Qisaq
    Qis_expanded = jnp.expand_dims(Qis, axis=2)  # shape (N, Z, 1)
    baseline = jnp.repeat(Qis_expanded, self.M, axis=2)  # shape (N, Z, M)
    if self.method=='mean':  # take mean
        return Qisa
    elif self.method=='quantile':  # use quantile
        return quantile(self.tau, Qisaq, gpoints)
    elif self.method=='expectile':  # use expectile
        return expectile(self.tau, Qisaq, gpoints)
    elif self.method=='weights':  # use weights
        w = jnp.ones((self.N, self.Z, self.M, len(self.gridpoints)))
        n = jnp.newaxis
        upper = self.gridpoints[n,n,n,:] >= baseline[:,:,:,n]
        upper = upper * (self.wU - 1)
        w = w + upper
        wQisaq = weights(w, Qisaq, gpoints)
        Qisa = jnp.einsum('ijkl,l->ijk', wQisaq, gpoints)  # take the new distrib's
↪   mean
        return Qisa

DDRLSarsa.Qisa = Qisa
```

Here, we define an optimistic agent, and we compare neutral and optimistic estimates of the Q-value.

```
## Define optimistic agent

aei_opt = DDRLSarsa(env,
                    nr_reward_bins=29,
                    discretization_sigma=0.1,
                    learning_rates=0.1,
                    choice_intensities=50.,
                    discount_factors=0.4,
                    Rmin=-0.5,
                    Rmax=1.5,
                    use_prefactor=True,
                    method="weights",
                    wU=3)
```

```
Qq = aei.Qisaq(X)
qm = aei.Qisa(X)[0,0]
qm_opt = aei_opt.Qisa(X)[0,0]

# Plot Q-value distribution Qisaq for both arms
fig, axes = plt.subplots(1, 2, figsize=(8,3))
actions = ["safe", "risky"]
for a, ax in enumerate(axes):
    ax.spines[['right', 'top', 'left']].set_visible(False)
    ax.bar(aei.gridpoints, Qq[0,0,a,:], width=0.9*aei.gridwidth, color='k')
    ax.plot([qm[a], qm[a]], [0.0, 0.02], c='red', lw=5,
            label="neutral Q-value")
```

```
    ax.plot([qm_opt[a], qm_opt[a]], [0.0, 0.02], c='limegreen',
            lw=5, label="optimistic Q-value")
    ax.set_xlabel("v")
    ax.set_ylabel(r"$Q^{i,s,a,v}$")
    ax.set_title("Q-value distribution for "+str(actions[a])+" action")
    ax.set_ylim([0, np.max(Qq[0,0,:,:]+0.01)])
axes[1].legend(bbox_to_anchor=(0.9, 0.6))

plt.show()
```

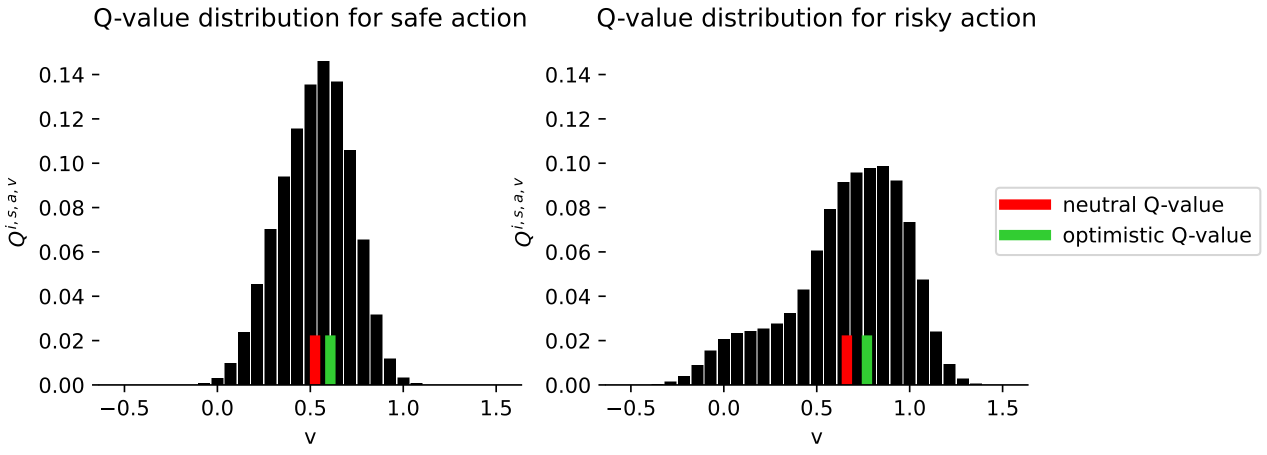

#### 3.5 Temporal-difference error

From the Q-value estimate, we can compute the temporal-difference error:

```
#|export
@partial(jit, static_argnums=(0,2))
def TDerror(self:DDRLSarsa,
            Xisa,  # Behavior profiles
            norm=False) -> jnp.ndarray:
    """
    Compute reward-prediction/temporal-difference error for
    strategy SARSA dynamics, given joint strategy `Xisa`.
    """
    Q = self.Qisa(Xisa)
    n = jnp.newaxis
    E = Q - (1 / self.beta[:,n,n]) * jnp.log(Xisa)
    E *= self.beta[:,n,n]

    E = E - E.mean(axis=2, keepdims=True) if norm else E
    return E

DDRLSarsa.TDerror = TDerror
```

It is now possible to plot our SARSA agent's deterministic trajectory in phase space.

```python
# Flow Plot

fig, ax = plt.subplots(1,1, figsize=(3,3))
x = ([0], [0], [0])  # prosperous state
y = ([0], [1], [0])  # degraded state

# Flow plot
fp.plot_strategy_flow(aei, x, y, flowarrow_points = np.linspace(0.01 ,0.99, 9),
                      NrRandom=16, axes=ax)

# Example trajectory
Xinit = np.ones((1,2,2)) * 0.8
Xinit[0,1,0] = 0.2
Xinit[:,:,1] = 1 - Xinit[:,:,0]
trj, fpr = aei.trajectory(Xinit, Tmax=1000, tolerance=1e-6)
fp.plot_trajectories([trj], x, y, fprs=[fpr], axes=ax, cols=['red'])

ax.set_xlabel("prob. to act safe in prosp. state", fontsize=9)
ax.set_ylabel("prob. to act safe in deg. state", fontsize=9)

plt.show()
```

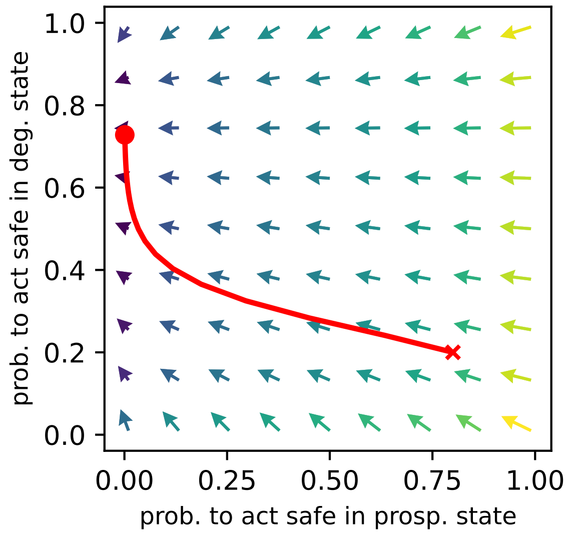

```python
#| hide
import nbdev; nbdev.nbdev_export()
```

### 4 DDRL Actor-Critic

In this section, we provide functions that are specific to the Actor-Critic version of Distributional Deterministic Reinforcement Learning.

First, we import everything necessary:

```
#| default_exp Agents.DDRLActorCritic
```

```
#| hide
# Imports for the nbdev development environment
import nbdev
from nbdev.showdoc import *
from fastcore.basics import patch
```

```
#| export
import numpy as np
from scipy.stats import norm

import jax.numpy as jnp
from jax import jit
from functools import partial

import itertools as it

from pyCRLD.Agents.Base import abase
from pyCRLD.Utils.Helpers import *
from pyCRLD.Utils.Helpers import compute_stationarydistribution

from pyDDRL.Agents.DDRLBase import DDRLBase

from pyDDRL.Agents.Methods import quantile, expectile, weights
```

```
# imports for this notebook
import matplotlib as mpl
import matplotlib.pyplot as plt
import seaborn as sns
from pyCRLD.Utils import FlowPlot as fp

# test environment
from pyDDRL.Environments.RiskRewardDilemma import RiskRewardDilemma

# regular Actor-Critic agent
from pyCRLD.Agents.StrategyActorCritic import stratAC
```

#### 4.1 Test environment

We use a risk-reward dilemma as a test environment. In this task, the agent can choose between a high-risk, high-gain option, and a low-risk, low-gain option. More precisely, we give our environment the following structure:

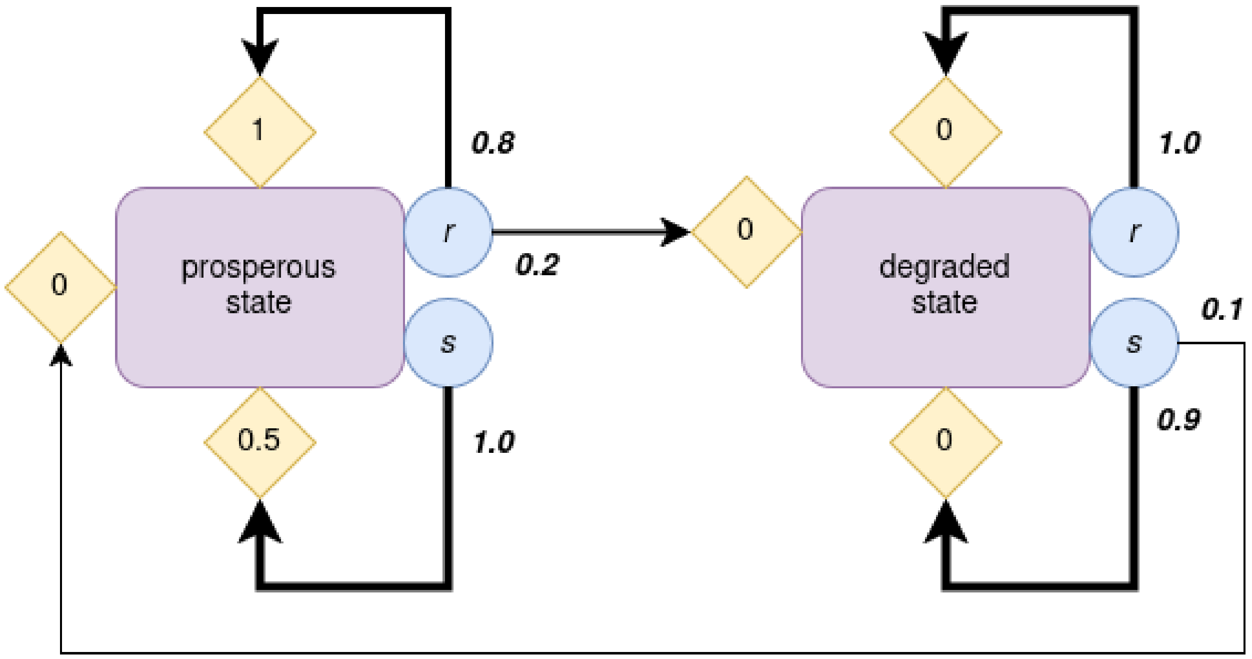

Figure 4.1: A risk-reward dilemma.

If an agent chooses the risky action $r$ in the prosperous state, it gets a higher reward. But there is a probability that it transitions to a degraded state, where it gets no reward. The only way to escape this degraded state is to choose the safe action, $s$.

```
# Define environment
env = RiskRewardDilemma(pc=0.2, pr=0.1, rc=0.5, rr=1, rd=0)
env
```

```
RiskRewardDilemma_0.2_0.1_0.5_1_0
```

#### 4.2 DDRL Actor-Critic class

Now, we can define the DDRL Actor-Critic class:

```
 #| export
class DDRLActorCritic(DDRLBase):
    """
    Base class for Actor-Critic version of DDRL in strategy space.
    """

    def __init__(self,
```

```
                 env,
                 **kwargs):

        super().__init__(env, **kwargs)
```

We consider the following agent-environment interface: * 1 agent with learning rate $\alpha = 0.1$ and discount factor $\gamma = 0.4$; * playing the aforementioned risk-reward dilemma.

```
aei = DDRLActorCritic(env,
                      nr_reward_bins=29,
                      discretization_sigma=0.1,
                      learning_rates=0.1,
                      discount_factors=0.4,
                      Rmin=-0.5,
                      Rmax=1.5,
                      use_prefactor=True)
```

```
WARNING:2025-12-21 19:07:00,244:jax._src.xla_bridge:794: An NVIDIA GPU may be present on this 
enabled jaxlib is not installed. Falling back to cpu.
```

#### 4.3 Computing distributions in DDRL-SARSA

Now we can compute and plot all the distributions we need.

##### 4.3.1 state-action value distribution $V^{i,s,a^i,\acute{v}}$

Just like state values, state-action values conform to the Bellman equation. Therefore, we can obtain the state-action value distribution for agent $i$, action $a^i$ and state $s$, from the next-state Q-value distribution for $(i, a, s)$, the reward distribution for $(i, a, s)$, and the smoothed Bellman Kronecker delta:

$$V^{i,s,a,v} = \sum_{r,\acute{v}} \delta^{i,v,r,\acute{v}} R^{i,s,a,r} \cdot {}^{next}V^{i,s,a,\acute{v}}$$

```
 #| export
@partial(jit, static_argnums=0)
def Visav(self:DDRLActorCritic,
          Xisa:jnp.ndarray, # Joint strategy
          Tisas:jnp.ndarray=None, # Optional transition for speed-up
          ) -> jnp.ndarray:  # Average state-action values
    """
    Compute Q-value distribution Visaq, given joint strategy `Xisa`.
    """
    Risar = self.Risar(Xisa)
    b_ivrv_ = self.b_ivrv_
    Visv = self.Visv(Xisa)
    Tisas = self.Tisas(Xisa) if Tisas is None else Tisas

    i = 0; s = 1; a = 2; s_ = 3; v = 4; v_ = 5; r = 6
    nextVisav = jnp.einsum(Tisas, [i,s,a,s_], Visv, [i,s_,v], [i,s,a,v],
```

```
                            optimize=self.opti)

    Visav = jnp.einsum(b_ivrv_, [i,v,r,v_], Risar, [i,s,a,r], nextVisav, [i,s,a,v_],
 ↪  [i,s,a,v])

    return Visav

DDRLActorCritic.Visav = Visav
```

We assume that, in both states, the agent takes the safe action with probability 0.2, and the risky action with probability 0.8.

```
# agent's strategy
X = np.ones((1, 2, 2)) * 0.2
X[:,:,1] = 1 - X[:,:,0]
X
```

```
array([[[0.2, 0.8],
        [0.2, 0.8]]])
```

Now we can plot the Q-value distribution for each action in the prosperous state.

```
Vv = aei.Visav(X)

# Plot Q-value distribution Visav for both actions
fig, axes = plt.subplots(1, 2, figsize=(8,3))
actions = ["safe", "risky"]
for a, ax in enumerate(axes):
    ax.spines[['right', 'top', 'left']].set_visible(False)
    ax.bar(aei.gridpoints, Vv[0,0,a,:], width=0.9*aei.gridwidth, color='k')
    rm = np.dot(aei.gridpoints, Vv[0,0,a,:])
    ax.plot([rm, rm], [0.0, 0.02], c='red', lw=5)
    ax.set_xlabel("v")
    ax.set_ylabel(r"$V^{i,s,a,v}$")
    ax.set_title("value distribution for "+str(actions[a])+" action")
    ax.set_ylim([0, np.max(Vv[0,0,:,:]+0.01)])
plt.show()
```

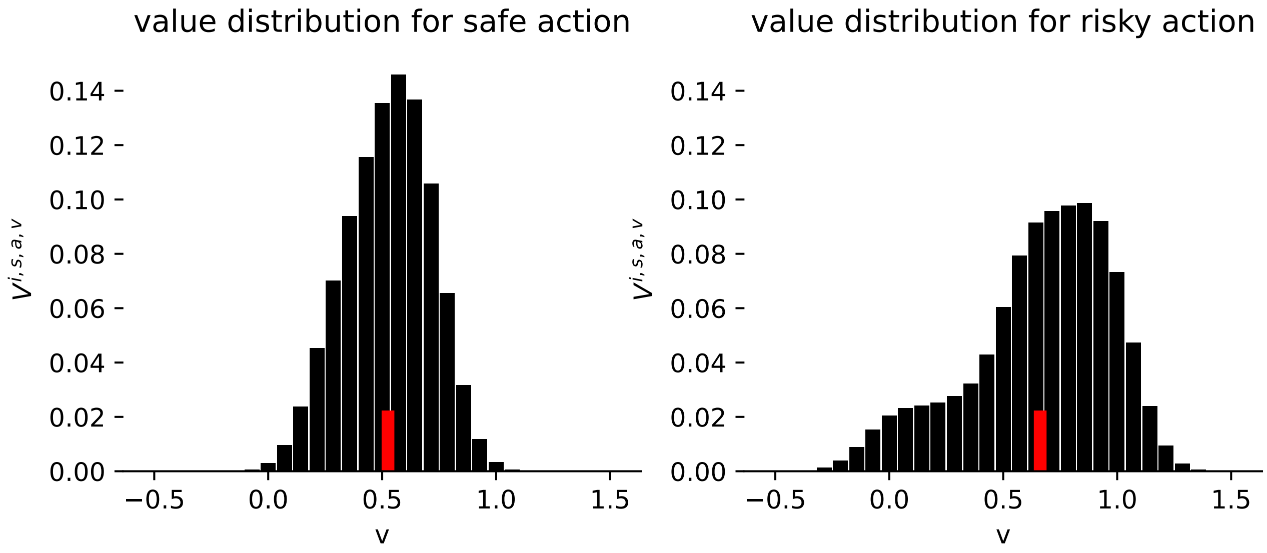

#### 4.4 (Biased) state-action value $V^{i,s,a^i}$

A classic estimate of the state-action value is the mean of the state-action value distribution. But a biased agent will get an optimistic or a pessimistic estimate of this distribution. To obtain this biased estimate, we can apply the quantile, expectile, or weights method to the value distribution. To know more about these methods, please refer to Chapter 5 .

Here, we compute the (biased) state-action value from the previous state-action value distribution. When the weights method is used, we take the state value $V^{i,s}$ as a baseline:

```
#| export
@partial(jit, static_argnums=0)
def Visa(self,
          Xisa:jnp.ndarray, # Joint strategy
          ) -> jnp.ndarray:  # Average state-action values
    """
    Compute Q-value for state `s` and action `a`.
    """
    Visav = self.Visav(Xisa)  # Q-value distrib
    gpoints = jnp.ones((self.nr)) * -1
    gpoints = gpoints.at[:].set(self.gridpoints)
    Visa = jnp.einsum('ijkl,l->ijk', Visav, gpoints)
    i = 0; a = 1; s = 2
    Vis = jnp.einsum(Visa, [i, s, a], Xisa, [i, s, a], [i, s])
    # Baseline: mean of Visaq
    Vis_expanded = jnp.expand_dims(Vis, axis=2)  # shape (N, Z, 1)
    baseline = jnp.repeat(Vis_expanded, self.M, axis=2)  # shape (N, Z, M)
    if self.method=='mean':  # take mean
        return Visa
    elif self.method=='quantile':  # use quantile
        return quantile(self.tau, Visav, gpoints)
    elif self.method=='expectile':  # use expectile
        return expectile(self.tau, Visav, gpoints)
    elif self.method=='weights':  # use weights
        w = jnp.ones((self.N, self.Z, self.M, len(self.gridpoints)))
```

```
        n = jnp.newaxis
        upper = self.gridpoints[n,n,n,:] >= baseline[:,:,:,n]
        upper = upper * (self.wU - 1)
        w = w + upper
        wVisav = weights(w, Visav, gpoints)
        Visa = jnp.einsum('ijkl,l->ijk', wVisav, gpoints)  # take the new distrib's
 ↪  mean
        return Visa

DDRLActorCritic.Visa = Visa
```

Here, we define an optimistic agent, and we compare neutral and optimistic estimates of the state-action value.

```
## Define optimistic agent

aei_opt = DDRLActorCritic(env,
                          nr_reward_bins=29,
                          discretization_sigma=0.1,
                          learning_rates=0.1,
                          discount_factors=0.4,
                          Rmin=-0.5,
                          Rmax=1.5,
                          use_prefactor=True,
                          method="weights",
                          wU=3)
```

```
Vv = aei.Visav(X)
vm = aei.Visa(X)[0,0]
vm_opt = aei_opt.Visa(X)[0,0]

# Plot state-action value distribution Visaq for both arms
fig, axes = plt.subplots(1, 2, figsize=(8,3))
actions = ["safe", "risky"]
for a, ax in enumerate(axes):
    ax.spines[['right', 'top', 'left']].set_visible(False)
    ax.bar(aei.gridpoints, Vv[0,0,a,:], width=0.9*aei.gridwidth, color='k')
    ax.plot([vm[a], vm[a]], [0.0, 0.02], c='red', lw=5,
            label="neutral value")
    ax.plot([vm_opt[a], vm_opt[a]], [0.0, 0.02], c='limegreen',
            lw=5, label="optimistic value")
    ax.set_xlabel("v")
    ax.set_ylabel(r"$Q^{i,s,a,v}$")
    ax.set_title("value distribution for "+str(actions[a])+" action")
    ax.set_ylim([0, np.max(Vv[0,0,:,:]+0.01)])
axes[1].legend(bbox_to_anchor=(0.9, 0.6))
plt.show()
```

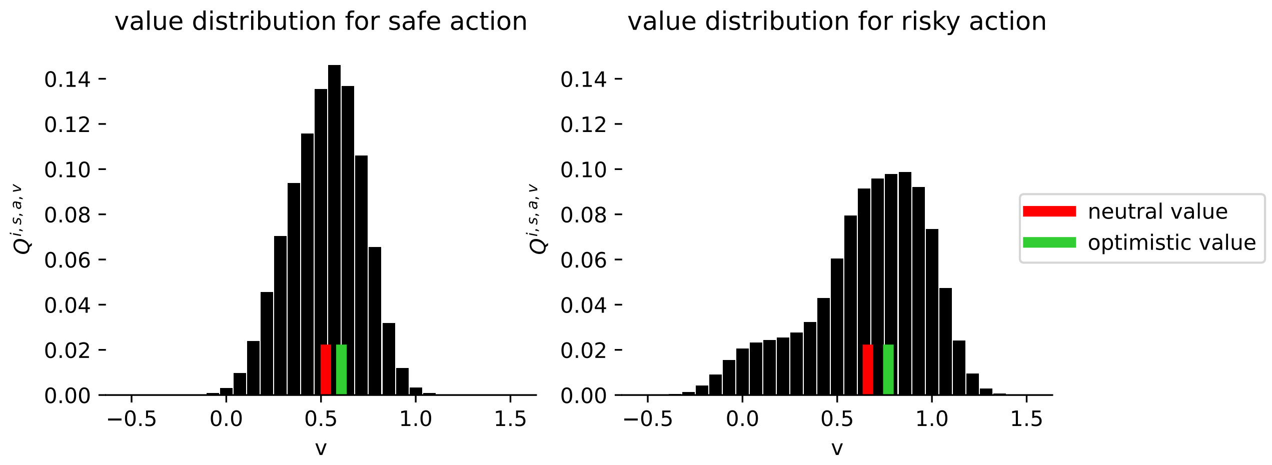

#### 4.5 Temporal-difference error

From the Q-value estimate, we can compute the temporal-difference error. For an Actor-Critic agent in the deterministic limit, this temporal-difference error is simply the state-action value.

```
 #|export
@partial(jit, static_argnums=(0,2))
def TDerror(self:DDRLActorCritic,
            Xisa,  # Behavior profiles
            norm=False) -> jnp.ndarray:
    """
    Compute reward-prediction/temporal-difference error for
    strategy actor-critic dynamics, given joint strategy `Xisa`.
    """
    V = self.Visa(Xisa)
    n = jnp.newaxis
    E = V
    E *= self.beta[:,n,n]

    E = E - E.mean(axis=2, keepdims=True) if norm else E
    return E

DDRLActorCritic.TDerror = TDerror
```

It is now possible to plot our actor-critic agent's deterministic trajectory in phase space.

```
# Flow Plot

fig, ax = plt.subplots(1,1, figsize=(3,3))
x = ([0], [0], [0])  # prosperous state
y = ([0], [1], [0])  # degraded state

# Flow plot
fp.plot_strategy_flow(aei, x, y, flowarrow_points = np.linspace(0.01 ,0.99, 9),
                      NrRandom=16, axes=ax)
```

```python
# Example trajectory
Xinit = np.ones((1,2,2)) * 0.8
Xinit[0,1,0] = 0.2
Xinit[:,:,1] = 1 - Xinit[:,:,0]
trj, fpr = aei.trajectory(Xinit, Tmax=1000, tolerance=1e-6)
fp.plot_trajectories([trj], x, y, fprs=[fpr], axes=ax, cols=['red'])

ax.set_xlabel("prob. to act safe in prosp. state", fontsize=9)
ax.set_ylabel("prob. to act safe in deg. state", fontsize=9)

plt.show()
```

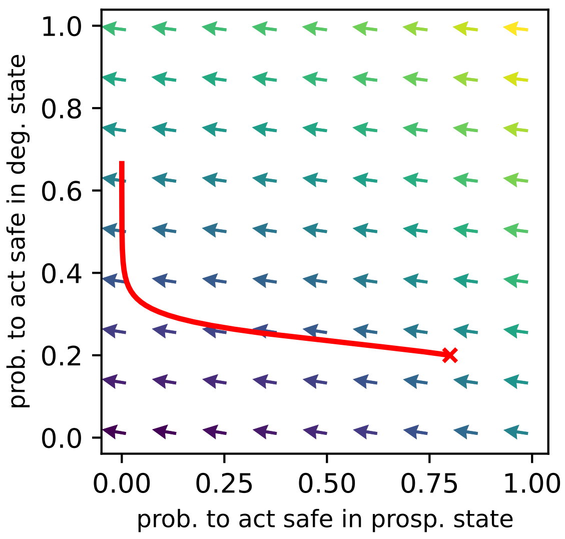

```python
#| hide
import nbdev; nbdev.nbdev_export()
```

### 5 Biasing methods

In this section, we introduce three different methods for implementing optimism/pessimism: * the quantile method * the expectile method * the weights method

First, we import what we need:

```
#| default_exp Agents.Methods
```

```
#| hide
# Imports for the nbdev development environment
import nbdev
from nbdev.showdoc import *
from fastcore.basics import patch
```

```
#| export
import numpy as np
import scipy.stats as stats

import jax.numpy as jnp
from jax import jit
from functools import partial

import itertools as it
```

```
# imports for testing
import matplotlib.pyplot as plt

# test environment
from pyDDRL.Environments.TwoArmedBandit import TwoArmedBandit

# test agent
from pyDDRL.Agents.DDRLSarsa import DDRLSarsa
```

#### 5.1 Test environment

As a test environment, we use the same one-state two-armed bandit as in Chapter 2:

```
# Define environment
xs = np.array([[-1, 1],
               [-1, 1]])
ps = np.array([[0.3, 0.7],
               [0.7, 0.3]])
tab_1s = TwoArmedBandit(disttype='discrete', xs=xs, ps=ps)
tab_1s.Rdict
```

```
{'x': array([[[[[-1.,  1.]],

          [[-1.,  1.]]]]]),
 'p': array([[[[[0.3, 0.7]],

          [[0.7, 0.3]]]]])}
```

#### 5.2 Quantile method

An optimistic estimate of the Q-value distribution can be obtained by selecting one of this distribution's quantiles instead of the mean. If $P$ is a distribution, then its $\tau$-quantile ($\tau \in [0,1]$) is a value $t$ such that:

$$\tau \int_{-\infty}^{t} P(x)dx = (1-\tau)\int_{t}^{\infty} P(x)dx$$

Thus, a distribution's 0.5-quantile is its median.

In our discretized version, we find an approximation of the $\tau$-quantile by summing all bin heights and taking the first grid point $x_k$ such that:

$$\sum_{i=0}^{k} b(x_i) \geq \tau$$

where $b(x_i)$ denotes the Q-distribution's bin height at value $x_i$.

Then, the deterministic approximation of the temporal-difference error is calculated using the Q-value distribution's chosen $\tau$-quantile instead of its mean.

```
#| export
def quantile(tau, dist, gpoints):
    """
    Computes the `tau`-quantile of any distribution `dist`,
    given a set of gridpoints `gpoints`.
    """
    dim = dist[..., -1].shape  # shape of the distrib except last axis
    qdist0 = np.ones(dim) * -1
    qdist = jnp.ones(dim) * -1  # will store the quantiles for each axis
    sums = jnp.cumsum(dist, axis=-1)  # add up bins
    mask = sums < tau  # find tau such that sum >= tau
    qidx = jnp.sum(mask, axis=-1)  # indices of the tau-quantile for each axis
    for index, _ in np.ndenumerate(qdist0):
        idx = qidx[index] - 1
        qdist = qdist.at[index].set((gpoints[idx]+gpoints[idx+1])/2)  # quantile
        # is between these two points
    return qdist
```

#### 5.3 Expectile method

An expectile is to the mean what a quantile is to the median. According to a more formal definition, $t$ is the $\tau$-expectile of a probability distribution with cumulative distribution function $F$ if:

$$(1-\tau)\int_{-\infty}^{t}(t-x)dF(x) = \tau\int_{t}^{\infty}(x-t)dF(x)$$

In our discretized framework, let us call $x_i$, $i \in [0, n]$ the gridpoints. We call $F_i$ the value of the cumulative distribution function at $x_i$. Then the continuous formula is equivalent to:

$$(1-\tau)\sum_{i=0}^{k}(x_k - x_i)\frac{F_i - F_{i-1}}{x_i - x_{i-1}} = \tau\sum_{i=k+1}^{n}(x_i - x_k)\frac{F_i - F_{i-1}}{x_i - x_{i-1}}$$

with $x_k$ the gridpoint that fulfills this condition. For all $i \in [1, n]$, $x_i - x_{i-1} = w$, with $w$ the gap between two gridpoints. Moreover, for all $i \in [1, n]$, $F_i - F_{i-1} = b_i$, where $b_i$ is the value of the probability distribution at $x_i$. Therefore, the equation becomes:

$$(1-\tau)\sum_{i=0}^{k}(x_k - x_i)b_i = \tau\sum_{i=k+1}^{n}(x_i - x_k)b_i$$

or:

$$\frac{\sum_{i=0}^{k}(x_k - x_i)b_i}{\sum_{i=k+1}^{n}(x_i - x_k)b_i} = \frac{\tau}{1-\tau}$$

In our discretized version, we find an approximation of the $\tau$-expectile by summing all bin heights and taking the first value $x_k$ such that:

$$\frac{\sum_{i=0}^{k}(x_k - x_i)b_i}{\sum_{i=k+1}^{n}(x_i - x_k)b_i} \geq \frac{\tau}{1-\tau}$$

```
 #| export
def expectile(tau, dist, gpoints):
    """
    Computes the `tau`-expectile of any distribution `dist`,
    given a set of gridpoints `gpoints`.
    """
    dim = dist.shape  # shape of dist
    dim1 = dist[..., -1].shape  # shape of dist - last axis

    # Constructing the b_i matrix
    bdiffs = np.zeros(dim[:-1]+(len(gpoints), len(gpoints)))
    bdiffs = np.repeat(dist[..., np.newaxis], len(gpoints), axis=-1)
    bdiffs = jnp.rot90(bdiffs, 1, axes=(-2, -1))

    # Constructing the x_i and x_k matrices
    xi = jnp.zeros((len(gpoints), len(gpoints)))
    xk = jnp.zeros((len(gpoints), len(gpoints)))
    xi = xi.at[:, :].set(gpoints)
```

```
    xk = xk.at[:, :].set(gpoints)
    xk = xk.T

    # Computing x_i - x_k and x_k - x_i (on same matrix)
    xdiffs = jnp.zeros(dim[:-1]+(len(gpoints), len(gpoints)))
    adiffs = jnp.abs(xi - xk)
    xdiffs = xdiffs.at[..., :, :].set(adiffs)

    # Computing (x_i - x_k) * b_i and (x_k - x_i) * bi
    xb = xdiffs * bdiffs
    xb_num = jnp.tril(xb, 0)  # (x_k - x_i) * bi
    xb_den = jnp.triu(xb, 0)  # (x_i - x_k) * b_i

    # Computing sum{(x_k - x_i)*b_i}/sum{(x_i - x_k)*b_i} for all k
    candidates = jnp.sum(xb_num[..., :-1, :], axis=-1) / jnp.sum(xb_den[..., :-1,
↪   :], axis=-1)

    # Compute expectiles
    edist0 = np.ones(dim1) * -1
    edist = jnp.ones(dim1) * -1  # will store expectiles
    mask = candidates < tau / (1 - tau)  # find the expectile
    eidx = jnp.sum(mask, axis=-1)  # expectile indices
    for index, _ in np.ndenumerate(edist0):
        idx = eidx[index] - 1
        edist = edist.at[index].set((gpoints[idx] + gpoints[idx+1]) / 2)
    return edist
```

#### 5.4 Weights method

The weights method multiplies the distribution's bins by a weight vector $w$. To implement optimism or pessimism to a distribution $P$, we usually apply a stepwise function – i.e., for all grid values $x_j$ that are below a certain baseline (for instance, the mean of the Q-value distribution), $w$ is 1, and for all grid values that are above this baseline, $w$ is $w_U$, where $w_U < 1$ if the agent is pessimistic, and $w_U > 1$ if the agent is optimistic. The resulting distribution, $\tilde{P}$, is then normalized:

$$\tilde{P} \leftarrow \frac{\tilde{P}}{\int_{x_1-\frac{\Delta x}{2}}^{x_N+\frac{\Delta x}{2}} \tilde{P} dv}$$

```
#| export
def weights(w, dist, gpoints):
    """
    Computes a weighted estimate of any distribution `dist`,
    given a set of gridpoints `gpoints`.
    """
    new_dist = w * dist  # apply the weights
    # Normalize the new distributions
    norm = jnp.sum(new_dist, axis=-1)
    norm = jnp.repeat(norm[..., jnp.newaxis], len(gpoints), axis=-1)
    new_dist = new_dist / norm
    return new_dist
```

#### 5.5 Comparison of the 3 methods on a two-armed bandit task

Here, we compare the optimistic estimates of Q-values given by the three methods. First, we obtain a Q-value distribution from a neutral DDRL-SARSA agent, assuming a random strategy.

```
# Agent-env interface
aei = DDRLSarsa(tab_1s, nr_reward_bins=29, discretization_sigma=0.1,
                learning_rates=0.1, choice_intensities=4.,
                discount_factors=0., Rmin=-2, Rmax=2)
```

```
WARNING:2025-12-21 19:07:37,935:jax._src.xla_bridge:794: An NVIDIA GPU may be present on this 
enabled jaxlib is not installed. Falling back to cpu.
```

```
# Random strategy
X = np.ones((1,1,2)) * 0.5
```

```
# Q-value distrib
Qq = aei.Qisaq(X)

# Plot Q-value distrib for arm 1
fig, ax = plt.subplots(1, 1, figsize=(4,3))
ax.spines[['right', 'top', 'left']].set_visible(False)
ax.bar(aei.gridpoints, Qq[0,0,0,:], width=0.9*aei.gridwidth, color='k')
rm = np.dot(aei.gridpoints, Qq[0,0,0,:])
ax.plot([rm, rm], [0.0, 0.01], c='red', lw=5)
ax.set_xlabel("v")
ax.set_ylabel(r"$Q^{i,s,a,v}$")
ax.set_title("Q-value distribution for arm 1")
```

```
Text(0.5, 1.0, 'Q-value distribution for arm 1')
```

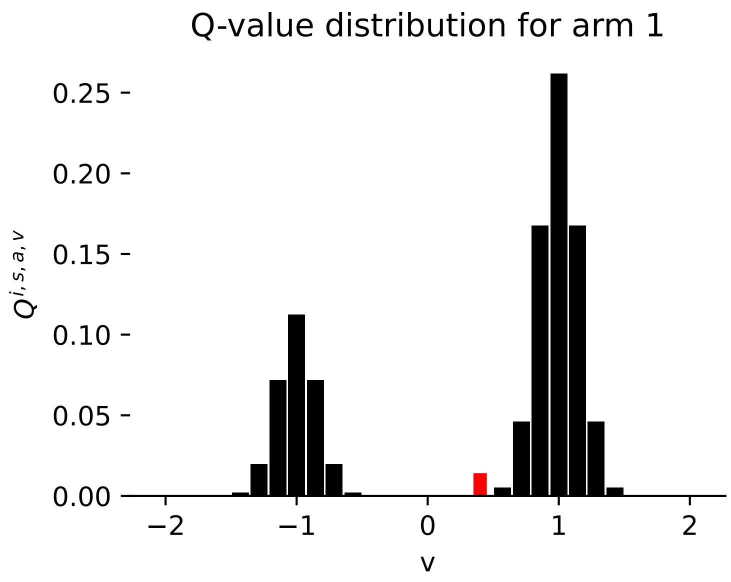

Now, we can compute and plot the different optimistic estimates of this distribution.

```
# Quantile
tau = 0.8
q_estimate = quantile(tau, Qq[0,0,0,:], aei.gridpoints)

# Expectile
e_estimate = expectile(tau, Qq[0,0,0,:], aei.gridpoints)

# Weights
# First, we need to create the weights vector
wU = 2  # upper weights value
w = np.ones((len(aei.gridpoints)))  # future weights vector
gpoints = jnp.ones((aei.nr)) * -1
gpoints = gpoints.at[:].set(aei.gridpoints)
Qisa = jnp.einsum('ijkl,l->ijk', Qq, gpoints)
i = 0; a = 1; s = 2
Qis = jnp.einsum(Qisa, [i, s, a], X, [i, s, a], [i, s])  # baseline
mask = aei.gridpoints >= Qis  # which part of the distrib
# is above baseline
mask = mask * (wU - 1)
w = w + mask  # weights vector

# Then, we get the weighted distribution
new_distrib = weights(w, Qq[0,0,0,:], aei.gridpoints)

# We can now compute the estimate
w_estimate = np.dot(new_distrib, aei.gridpoints)
```

```
# Q-value distrib
Qq = aei.Qisaq(X)

# Plot Q-value distrib for arm 1
fig, ax = plt.subplots(1, 1, figsize=(4,3))
ax.spines[['right', 'top', 'left']].set_visible(False)
qm = np.dot(aei.gridpoints, Qq[0,0,0,:])  # compute mean
ax.bar(aei.gridpoints, Qq[0,0,0,:], width=0.9*aei.gridwidth, color='k')
ax.plot([q_estimate, q_estimate], [0.0, 0.03], c='orange', lw=3,
        label='0.8-quantile')
ax.plot([e_estimate, e_estimate], [0.0, 0.03], c='limegreen', lw=3,
        label='0.8-expectile')
ax.plot([w_estimate, w_estimate], [0.0, 0.03], c='mediumpurple', lw=3,
        label='weight '+r'$w=2$')
ax.plot([qm, qm], [0.0, 0.03], c='red', lw=3,
        label='mean')
ax.set_xlabel("v")
ax.set_ylabel(r"$Q^{i,s,a,v}$")
ax.legend(bbox_to_anchor=(0.9, 0.6))
ax.set_title("Q-value distribution for arm 1")

plt.show()
```

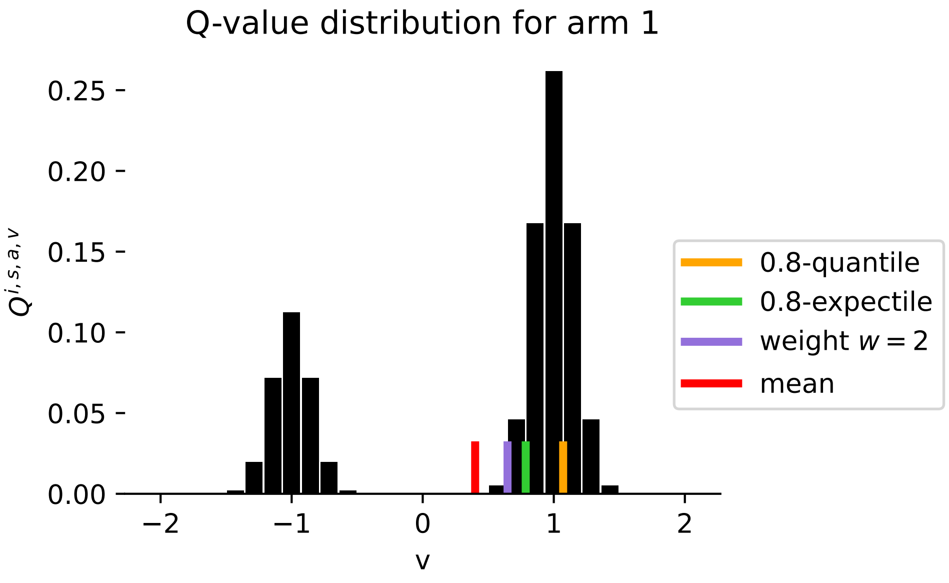

```
#| hide
import nbdev; nbdev.nbdev_export()
```

### Part II

### Environments

In this section, we introduce three environment classes that are widely used in various disciplines, such as cognitive psychology or sustainability studies: a two-armed bandit task ( Chapter 6 ), a social dilemma ( Chapter 7 ), and a risk-reward dilemma ( Chapter 8 ).

We will use these environments to compute our results in **?@sec-results** .

### 6 Two-armed bandit

In this section, we introduce a simple choice task: the two-armed bandit. This task is a well-known paradigm for studying decision-making in experimental psychology. In this environment, an agent finds itself in a decision state, $s_0$. It can choose between two options: * a first arm, $a_1$, which gives rewards according to a random variable $X_1$ following probability distribution $P_1$; * a second arm, $a_2$, which gives rewards according to a random variable $X_2$ following probability distribution $P_2$. Often, in experimental psychology, these probability distributions are discrete and bimodal. The agent must then learn which arm is more rewarding on average, by sampling the bandit repeatedly.

Here is an example, where $a_1$ yields a reward $+1$ with probability $p_1$ (and a punishment $-1$ with probability $1 - p_1$), and $a_2$ yields a a reward $+1$ with probability $p_2$ (and a punishment $-1$ with probability $1 - p_2$):

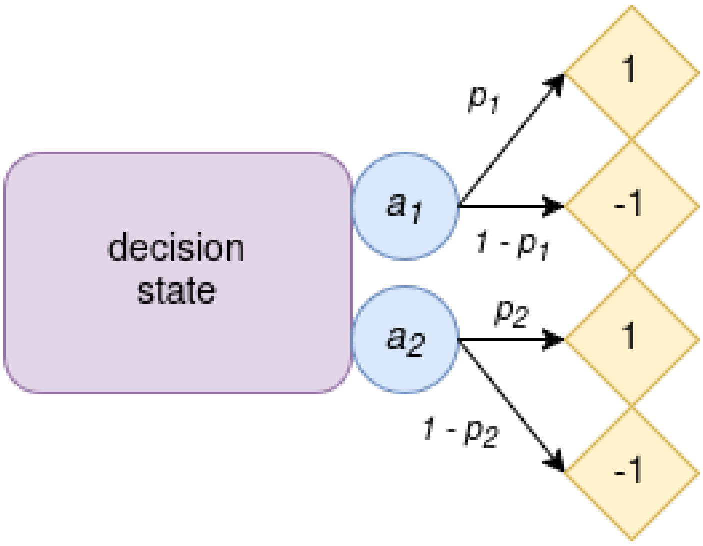

Figure 6.1: A two-armed bandit.

Let's implement a two-armed bandit task.

First, we import what we need:

```
#| default_exp Environments.TwoArmedBandit
```

```
#| hide
# Imports for the nbdev development environment
import nbdev
from nbdev.showdoc import *
from fastcore.basics import patch
```

```
# imports for testing
import matplotlib.pyplot as plt
import numpy as np

# test agents
from pyCRLD.Agents.Base import abase
from pyCRLD.Agents.StrategySARSA import stratSARSA

from pyDDRL.Agents.DDRLBase import DDRLBase
from pyDDRL.Agents.DDRLSarsa import DDRLSarsa

# utils
from pyCRLD.Utils import FlowPlot as fp
from pyDDRL.Utils.Trajectories1D import trajectories_1D
from pyDDRL.Utils.PhasePlot import phaseplot
```

```
#| export
from pyCRLD.Environments.Base import ebase
from pyCRLD.Utils.Helpers import make_variable_vector

from fastcore.utils import *
from fastcore.test import *

from typing import Iterable
import numpy as np
import scipy.stats as stats
```

```
#| export
class TwoArmedBandit(ebase):
    """
    Distributional Two-Armed Bandit Environment with a single state.
    """

    def __init__(self,
                 disttype="continuous", # continuous or discrete
                 distA=None, # reward distribution for option A
                 distB=None,  # reward distribution for option B
                 xs=None,  # reward values
                 ps=None):  # reward probabilities

        self.N = 1  # number of agents
        self.M = 2  # number of actions
        self.Z = 1  # number of states

        self.disttype = disttype  # distribution type: "continuous" or "discrete" or
         ↪ None
        self.dist = [distA, distB]  # dist shape for arm 1 and for arm 2
        self.xs = xs  # reward values
        self.ps = ps  # reward probabilities

        self.Aset = self.actions()  # to have them available for the creation
        self.Sset = self.states()   # of the Transition and Reward Tensors
```

```
        self.state = 1 # initial state

        self.T = self.TransitionTensor()
        self.R = self.RewardTensor()
        self.Rdict = self.RewardDict()
        self.F = np.array(self.FinalStates())
        # super().__init__()
```

```
#| export
@patch
def actions(self:TwoArmedBandit):
    """The action set"""
    return [['A', 'B']]  # two actions
```

```
#| export
@patch
def states(self:TwoArmedBandit):
    """The states set"""
    return ['O']  # 1 state
```

```
#| export
@patch
def TransitionTensor(self:TwoArmedBandit):
    """Get the Transition Tensor."""
    dim = np.concatenate(([self.Z],
                          [self.M],
                          [self.Z]))
    Tsas = np.ones(dim) * (-1)

    for index, _ in np.ndenumerate(Tsas):
        Tsas[index] = 1.  # always come back to decision state
    return Tsas
```

```
#| export
@patch
def RewardTensor(self:TwoArmedBandit):
    """Get the Reward Tensor R[i,s,a1,...,aN,s']."""
    dim = np.concatenate(([self.N],
                          [self.Z],
                          [self.M],
                          [self.Z]))
    Risas = np.zeros(dim)

    for index, _ in np.ndenumerate(Risas):
        Risas[index] = self._reward(index[0], index[1], index[2],
                                    index[3])
    return Risas

@patch
```

```python
def _reward(self:TwoArmedBandit,
            i:int, # the agent index
            s:int, # the state index
            a:int, # indices for joint actions
            s_:int  # the next-state index
            ) -> float:  # reward value
    """
    Returns the reward value for agent `i` in current state `s`, under joint action
↪   `jA`, when transitioning to next state `s_`.
    """
    if self.Aset[0][a] == 'A':  # if action was A
        if self.disttype == 'continuous':
            return self.dist[0][1]  # the agents receive mean reward
        else:  # discrete
            return np.dot(self.xs[0, :], self.ps[0, :])

    else:  # if action was B
        if self.disttype == 'continuous':
            return self.dist[1][1]  # the agents receive mean reward
        else:  # discrete
            return np.dot(self.xs[1, :], self.ps[1, :])
```

```python
#| export
@patch
def RewardDict(self:TwoArmedBandit):
    """Get the Reward Dict."""

    if self.disttype == 'continuous':
        dim = np.concatenate(([self.N],
                              [self.Z],
                              [self.M],
                              [self.Z]))
        loc = np.zeros((dim))  # mean reward
        scale = np.zeros((dim))  # sd
        distr = np.empty((dim), dtype=object)  # distribution type

        loc[:, :, 0, :] = self.dist[0][1]  # mean R after choosing option A
        loc[:, :, 1, :] = self.dist[1][1]  # mean R after choosing option B
        scale[:, :, 0, :] = self.dist[0][2]
        scale[:, :, 1, :] = self.dist[1][2]
        distr[:, :, 0, :] = self.dist[0][0]
        distr[:, :, 1, :] = self.dist[1][0]

        # Create dict with previous arrays
        d = dict.fromkeys(["dist", "loc", "scale"])
        d['dist'] = distr
        d['loc'] = loc
        d['scale'] = scale

    else:  # discrete
        dim = np.concatenate(([self.N],
                              [self.Z],
```

```
                                [self.M],
                                [self.Z],
                                [len(self.xs[0, :])]))
        x = np.zeros((dim))
        p = np.zeros((dim))
        x[:, :, 0, :, :] = self.xs[0, :]
        x[:, :, 1, :, :] = self.xs[1, :]
        p[:, :, 0, :, :] = self.ps[0, :]
        p[:, :, 1, :, :] = self.ps[1, :]

        # Create dict with previous arrays
        d = dict.fromkeys(["x", "p"])
        d['x'] = x
        d['p'] = p

    return d
```

```
 #| export
@patch
def id(self:TwoArmedBandit):
    """
    Returns id string of environment
    """
    # Default
    if self.disttype == 'continuous':
        dA = self.dist[0][0]
        dB = self.dist[1][0]
        muA = self.dist[0][1]
        muB = self.dist[1][1]
        id = f"{self.__class__.__name__}_"+\
        f"{dA}_{str(muA)}_{dB}_{str(muB)}"

    else:  # discrete
        rew = self.xs[0, 1]
        pun = self.xs[0, 0]
        pA = self.ps[0, 1]
        pB = self.ps[1, 1]
        id = f"{self.__class__.__name__}_"+\
        f"{str(rew)}_{str(pun)}_{str(pA)}_{str(pB)}"

    return id
```

```
 #| export
@patch
def FinalStates(self:TwoArmedBandit):
    """Default final states: no final states"""
    return np.zeros(self.Z, dtype=int)
```

#### 6.1 Stochastic two-armed bandit task with discrete return

Here, we implement the above example of a standard two-armed bandit task in experimental psychology. In this task, an agent must choose between two options, $a_1$ and $a_2$. $a_1$ gives a reward $+1$ with probability $p_1$, and a punishment $-1$ with probability $1 - p_1$. $a_2$ gives a reward $+1$ with probability $p_2$, and a punishment $-1$ with probability $1 - p_2$. Choosing repeatedly between the two options, the agent must learn which one is most rewarding on average. We model the two-armed bandit task as a simple, 1-state Markov environment:

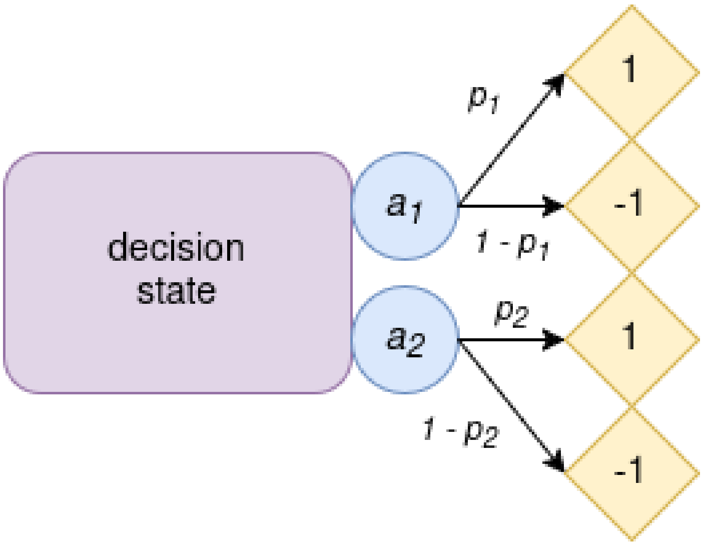

Figure 6.2: Two-armed bandit.

Below, we define a two-armed bandit task with $p_1 = 0.7$ and $p_2 = 0.3$.

```
# Define environment
# reward values
xs = np.array([[-1, 1],  # first arm
               [-1, 1]])  # second arm
# reward probs
ps = np.array([[0.3, 0.7],  # first arm
               [0.7, 0.3]])  # second arm
tab = TwoArmedBandit(disttype='discrete', xs=xs, ps=ps)
```

Now, we define an agent-environment interface comprised of a SARSA agent playing the aforementioned task. This agent has a learning rate $\alpha = 0.1$, a discount factor $\gamma = 0$, and a choice intensity $\beta = 4$.

```
# Agent-environment interface
aei = stratSARSA(tab,
                 learning_rates=0.1,
                 discount_factors=0,
                 choice_intensities=4.,
                 use_prefactor=False)
```

```
WARNING:2025-12-21 19:08:16,542:jax._src.xla_bridge:794: An NVIDIA GPU may be present on this 
enabled jaxlib is not installed. Falling back to cpu.
```

We can plot the agent's deterministic trajectories and phase plot. Here, the "strategy" $X$ denotes the probability to choose the most rewarding arm. Both plots show what strategy the agent converges to. In the phase plot, the fixed points are the intersections of the $dX$ curve with 0. In the phase plot, we also showed the value of the RPE given a strategy $X$ (in blue).

```
# Figure
x = ([0], [0], [0])  # (i,s,a) to plot along the x-axis
res = np.linspace(0, 1., 11)  # initial strategies
res[0] = 0.0001
res[-1] = 0.9999
res_phase = np.linspace(0, 1., 51)  # resolution for phase plot
res_phase[0] = 0.001
res_phase[-1] = 0.999

# Plot deterministic trajectories and phase plot
fig, (ax1, ax2) = plt.subplots(1, 2, figsize=(8, 3))
trajectories_1D(aei, x, res, NrRandom=1, traj_len=70, axes=[ax1])
phaseplot(aei, x, res_phase, NrRandom=1, axes=[ax2])
ax1.set_title("deterministic trajectories")
ax1.set_xlabel("time step "+r'$t$')
ax1.set_ylabel("probability to choose arm 1")
ax2.set_xlabel("strategy "+r'$X$')
ax2.set_title("phase plot")
plt.show()
```

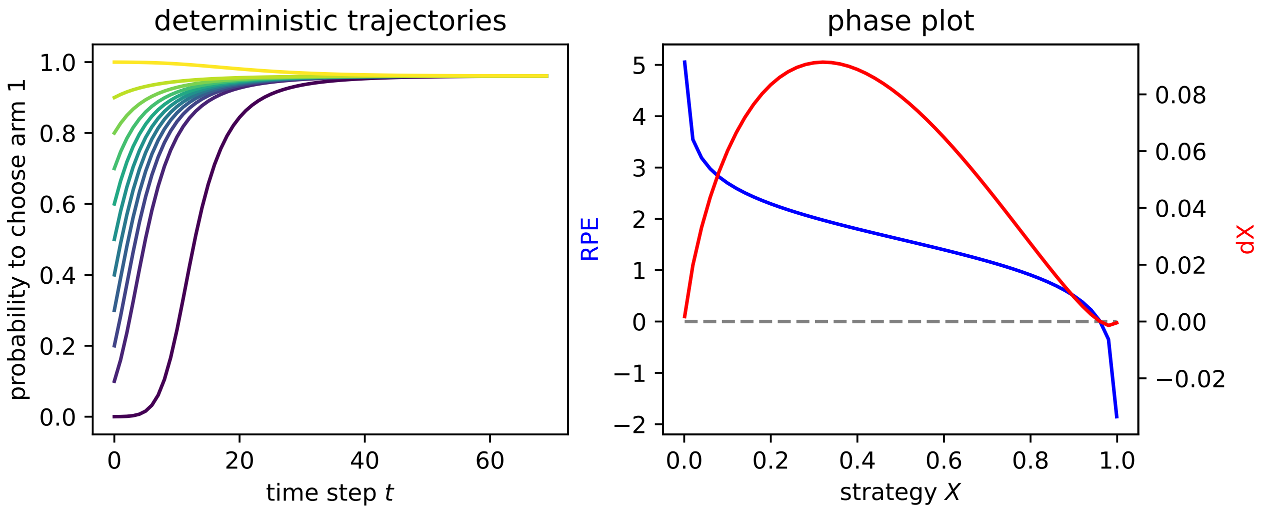

#### 6.2 Stochastic two-armed bandit with continuous return

Although we do not use this feature in our article, it is possible to implement a stochastic two-armed bandit with continuous return, where the random return follows a continuous distribution.

Let's consider the following example, in which an agent can choose between two arms: * the first arm gives a random return that follows a Gaussian distribution, centered around $\mu = 1$, with standard deviation $\sigma = 0.5$ * the second arm gives a random return that follows a Gaussian distribution, centered around $\mu = -1$, with standard deviation $\sigma = 2.0$

```
# Define environment

distA = ('norm', 1, 0.5)  # first arm
distB = ('norm', -1, 2)  # second arm

tab_cr = TwoArmedBandit(distA=distA, distB=distB,
                        disttype="continuous")
tab_cr
```

```
TwoArmedBandit_norm_1_norm_-1
```

Now let's plot the distribution of the random return for each arm.

```
## Plot the return distributions

x = np.linspace(-10, 10)
norm1 = stats.norm.pdf(x, 1, 0.5)  # first arm
norm2 = stats.norm.pdf(x, -1, 2)  # second arm
norms = [norm1, norm2]

# Plot
fig, axes = plt.subplots(1, 2, figsize=(5, 1.5))
for s_, ax in enumerate(axes):
    ax.spines[['right', 'top', 'left']].set_visible(False)
    ax.plot(x, norms[s_], color='blue')
    ax.set_xlabel("r")
    ax.set_ylim([0,0.8])
    ax.set_title("random return for arm "+str(s_+1), fontsize=10)
axes[0].set_ylabel(r'$p(R = r)$')
axes[1].set_yticklabels([])
axes[1].set_yticks([])
plt.show()
```

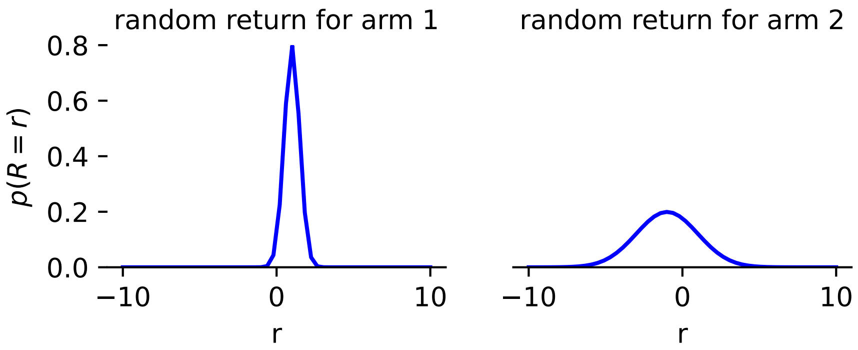

It is possible to use this stochastic two-armed bandit as part of an agent-environment interface. The above distribution will then be discretized by the agent (see Chapter 2 ).

We define an agent-environment interface comprised of a distributional SARSA agent playing the aforementioned task. This agent has a learning rate $\alpha = 0.1$, a discount factor $\gamma = 0$, and a choice intensity $\beta = 4$.

```
# Agent-environment interface
aei = DDRLSarsa(tab_cr,
                nr_reward_bins=29,
                discretization_sigma=0.001,
                learning_rates=0.1,
                choice_intensities=4.,
                discount_factors=0.,
                Rmin=-5,
                Rmax=5,
                use_prefactor=False)
```

Now plotting the agent's deterministic trajectories and phase plot:

```
# Figure
x = ([0], [0], [0])  # (i,s,a) to plot along the x-axis
res = np.linspace(0, 1., 11)  # initial strategies
res[0] = 0.0001
res[-1] = 0.9999
res_phase = np.linspace(0, 1., 51)  # resolution for phase plot
res_phase[0] = 0.001
res_phase[-1] = 0.999

# Plot deterministic trajectories and phase plot
fig, (ax1, ax2) = plt.subplots(1, 2, figsize=(8, 3))
trajectories_1D(aei, x, res, NrRandom=1, traj_len=70, axes=[ax1])
phaseplot(aei, x, res_phase, NrRandom=1, axes=[ax2])
ax1.set_title("deterministic trajectories")
ax1.set_xlabel("time step "+r'$t$')
ax1.set_ylabel("probability to choose arm 1")
ax2.set_xlabel("strategy "+r'$X$')
ax2.set_title("phase plot")
plt.show()
```

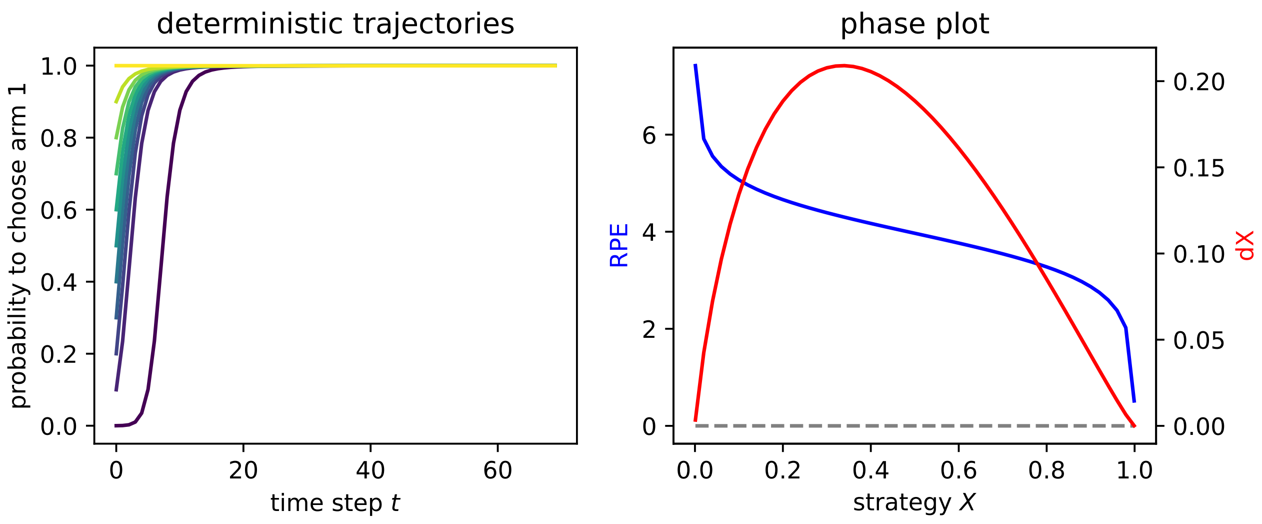

```
#| hide
import nbdev; nbdev.nbdev_export()
```

### 7 Social dilemma

Here, we consider two social dilemmas: the stag-hunt game and the prisoner's dilemma.

The stag-hunt game is a type of social dilemma where two or more agents must coordinate on the best action. Hence, it is often called a "coordination problem". In our version, we consider two agents who must take one of two actions: cooperate ($c$) or defect ($d$).

If both agents cooperate, they get a payoff $R$ (called "reward"). If they both defect, they get a payoff $P$ (called "punishment"). If one cooperates and the other defects, the cooperator gets a payoff $S$ (called "sucker") and the defector, a payoff $T$ (called "temptation"). In a stag-hunt game, $R > T > P > S$. This means the agents' main challenge is coordination: cooperating at the same time in order to get the maximum reward.

Note that, if $T > R > P > S$, the game becomes a prisoner's dilemma: both agents are incentivized to defect, because the temptation is now the maximum payoff.

The structure of a social dilemma is shown below.

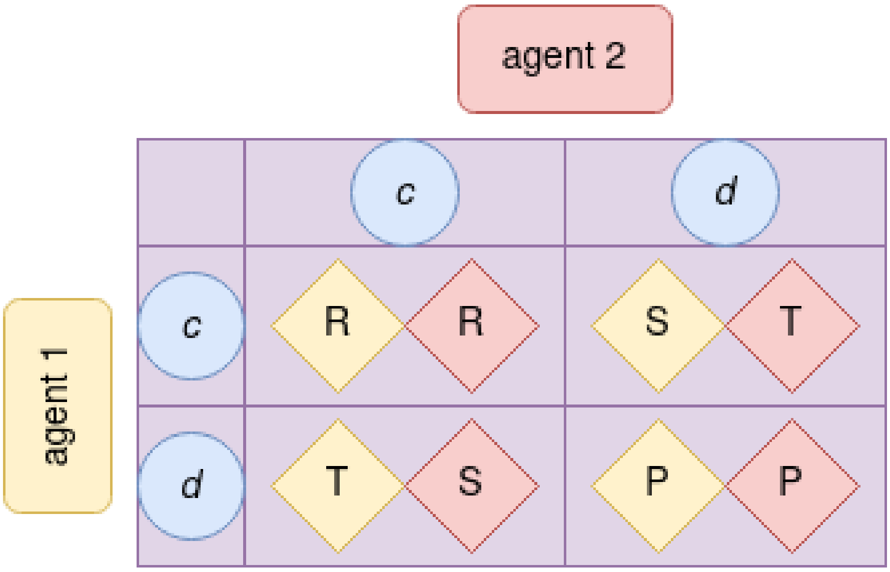

Figure 7.1: A social dilemma.

Let's implement a social dilemma.

First, we import what we need:

```
#| default_exp Environments.SocialDilemma
```

```
#| hide
# Imports for the nbdev development environment
import nbdev
from nbdev.showdoc import *
from fastcore.basics import patch
```

```
# imports for testing
import matplotlib.pyplot as plt
import numpy as np

# test agent
from pyCRLD.Agents.Base import abase
from pyCRLD.Agents.StrategySARSA import stratSARSA

# utils
from pyCRLD.Utils import FlowPlot as fp
from pyDDRL.Utils.Trajectories1D import trajectories_1D
from pyDDRL.Utils.PhasePlot import phaseplot
```

```
#| export
from pyCRLD.Environments.Base import ebase

from fastcore.utils import *
from fastcore.test import *

import numpy as np
```

```
#| export
class SocialDilemma(ebase):
    """
    Symmetric 2-agent 2-action Social Dilemma Matrix Game.
    """

    def __init__(self,
                 R:float,  # reward of mutual cooperation
                 T:float,  # temptation of unilateral defection
                 S:float,  # sucker's payoff of unilateral cooperation
                 P:float): # punsihment of mutual defection
        self.N = 2  # number of agents
        self.M = 2  # number of actions
        self.Z = 1  # number of states

        self.disttype = None  # deterministic rewards

        self.Re = R  # reward
        self.Te = T  # temptation
        self.Su = S  # sucker
        self.Pu = P  # punishment

        self.state = 0 # inital state
        super().__init__()
```

```
#| export
@patch
def TransitionTensor(self:SocialDilemma):
    """Get the Transition Tensor."""
    Tsas = np.ones((self.Z, self.M, self.M, self.Z))
```

```
    return Tsas
```

```
#| export
@patch
def RewardTensor(self:SocialDilemma):
    """Get the Reward Tensor R[i,s,a1,...,aN,s']."""

    R = np.zeros((2, self.Z, 2, 2, self.Z))

    R[0, 0, :, :, 0] = [[self.Re , self.Su],
                        [self.Te , self.Pu]]
    R[1, 0, :, :, 0] = [[self.Re , self.Te],
                        [self.Su , self.Pu]]

    return R
```

```
#| export
@patch
def actions(self:SocialDilemma):
    """The action sets"""
    return [['c', 'd'] for _ in range(self.N)]
```

```
#| export
@patch
def states(self:SocialDilemma):
    """The states set"""
    return ['.']
```

```
#| export
@patch
def id(self:SocialDilemma):
    """
    Returns id string of environment
    """
    # Default
    id = f"{self.__class__.__name__}_"+\
        f"{self.Te}_{self.Re}_{self.Pu}_{self.Su}"
    return id
```

#### 7.1 Example

We implement both a stag-hunt game and a prisoner's dilemma.

```
# Define environments
envSH = SocialDilemma(R=3, T=1, P=0, S=-2)  # stag hunt
envPD = SocialDilemma(R=1, T=3, P=0, S=-2)  # prisoner's dilemma
```

Now, we define an agent-environment interface comprised of a SARSA agent playing the aforementioned tasks. This agent has a learning rate $\alpha = 0.1$, a discount factor $\gamma = 0.9$, and a choice intensity $\beta = 50$.

```python
# Agent-environment interface
aeiSH = stratSARSA(envSH,
                   learning_rates=0.1,
                   discount_factors=0.9,
                   choice_intensities=50.,
                   use_prefactor=True)

aeiPD = stratSARSA(envPD,
                   learning_rates=0.1,
                   discount_factors=0.9,
                   choice_intensities=50.,
                   use_prefactor=True)
```

```
WARNING:2025-12-21 19:08:38,826:jax._src.xla_bridge:794: An NVIDIA GPU may be present on this 
enabled jaxlib is not installed. Falling back to cpu.
```

We plot the agent's deterministic trajectories in phase space on the flow plots below:

```python
# Figure
x = ([0], [0], [0])  # (i,s,a) to plot along the x-axis
y = ([1], [0], [0])  # (i,s,a) to plot along the y-axis

# Plot flow plot
fig, (ax1, ax2) = plt.subplots(1, 2, figsize=(7, 3))
fp.plot_strategy_flow(aeiSH, x, y, flowarrow_points = np.linspace(0.01 ,0.99, 9),
                      NrRandom=16, axes=ax1)
fp.plot_strategy_flow(aeiPD, x, y, flowarrow_points = np.linspace(0.01 ,0.99, 9),
                      NrRandom=16, axes=ax2)
ax1.set_title("Stag hunt")
ax2.set_title("Prisoner's dilemma")
ax1.set_xlabel("prob. agent 1 cooperates")
ax2.set_xlabel("prob. agent 1 cooperates")
ax1.set_ylabel("prob. agent 2 cooperates")
plt.show()
```

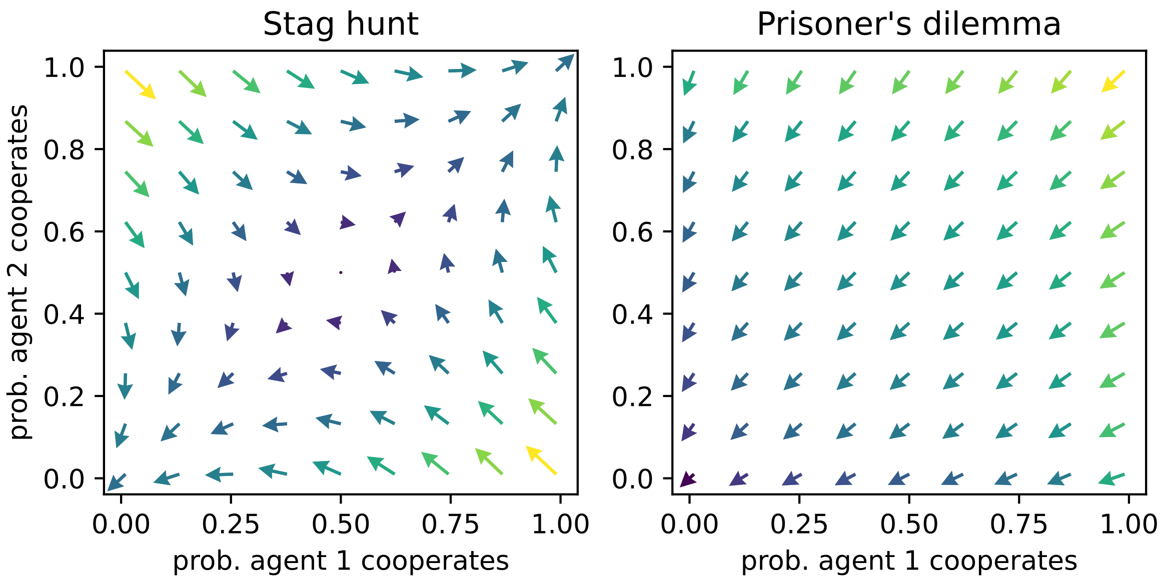

```
#| hide
import nbdev; nbdev.nbdev_export()
```

### 8 Risk-reward dilemma

In a risk-reward dilemma (Barfuss, 2022), an agent can choose between a risky action, $r$, and a safe action, $s$. The environment contains a prosperous state and a degraded state. In the prosperous state, the risky action yields a higher reward than the safe action. However, choosing $r$ also makes it more likely to transition to the degraded state, in which rewards are always null or overall undesirable. In the degraded state, only the safe action gives the agent a chance to transition back to the prosperous state.

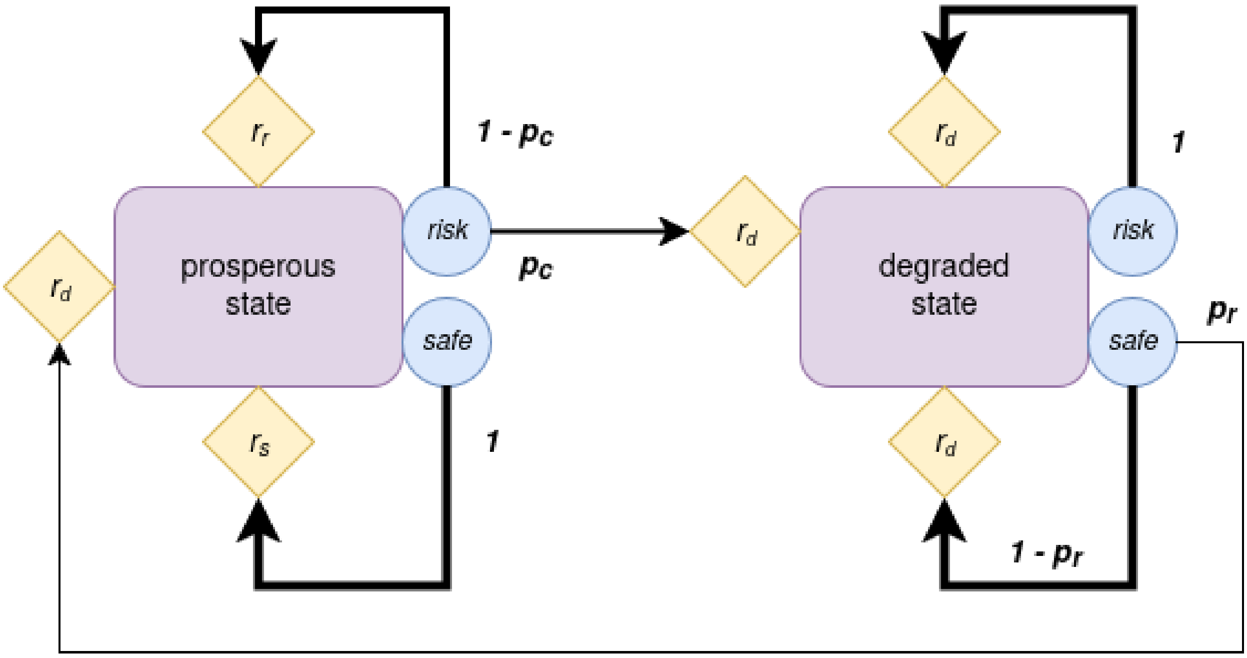

Figure 8.1: A risk-reward dilemma.

Let's implement a risk-reward dilemma.

First, we import what we need:

```
#| default_exp Environments.RiskRewardDilemma
```

```
#| hide
# Imports for the nbdev development environment
import nbdev
from nbdev.showdoc import *
from fastcore.basics import patch
```

```
# imports for testing
import matplotlib.pyplot as plt
import numpy as np
```

```
# test agent
from pyCRLD.Agents.Base import abase
from pyCRLD.Agents.StrategyActorCritic import stratAC

# utils
from pyCRLD.Utils import FlowPlot as fp
from pyDDRL.Utils.Trajectories1D import trajectories_1D
from pyDDRL.Utils.PhasePlot import phaseplot
```

```
#| export
from pyCRLD.Environments.Base import ebase
from pyCRLD.Utils.Helpers import make_variable_vector

from fastcore.utils import *
from fastcore.test import *

from typing import Iterable
import numpy as np
```

```
#| export
class RiskRewardDilemma(ebase):
    """
    Risk-Reward Dilemma.
    """

    def __init__(self,
                 pc:float,  # collapse probability
                 pr:float,  # recovery probability
                 rc:float,  # cautious reward
                 rr:float,  # risky reward
                 rd:float,  # reward in degraded state
                 asym=False,  # whether deg state is asymmetric
                 rdr=0.01):  # risky reward in deg state if deg state is asymmmetric

        self.N = 1
        self.M = 2
        self.Z = 2

        self.pc = pc
        self.pr = pr
        self.rc = rc
        self.rr = rr
        self.rd = rd
        if asym==True:
            self.rdr = rdr
        else:
            self.rdr = rd

        self.distype = None

        self.Aset = self.actions()  # to have them available for the creation
```

```python
        self.Sset = self.states()   # of the Transition and Reward Tensors

        self.state = 1 # initial state

        self.T = self.TransitionTensor()
        self.R = self.RewardTensor()
        self.F = np.array(self.FinalStates())
        # super().__init__()
```

```python
#| export
@patch
def actions(self:RiskRewardDilemma):
    """The action set"""
    return ['c', 'r']
```

```python
#| export
@patch
def states(self:RiskRewardDilemma):
    """The states set"""
    return ['p', 'd']
```

```python
#| export
@patch
def TransitionTensor(self:RiskRewardDilemma):
    """Get the Transition Tensor."""
    dim = np.concatenate(([self.Z],
                          [self.M for _ in range(self.N)],
                          [self.Z]))
    Tsas = np.ones(dim) * (-1)

    for index, _ in np.ndenumerate(Tsas):
        Tsas[index] = self._transition_probability(index[0],
                                                   index[1],
                                                   index[2])
    return Tsas

@patch
def _transition_probability(self:RiskRewardDilemma,
                            s:int, # the state index
                            a:int, # index for action
                            s_:int  # the next-state index
                           ) -> float:  # transition probability
    """
    Returns the transition probability for current state `s`, joint action `jA`, and
↪   next state `s_`.
    """
    transitionprob = 0

    if self.Sset[s] == 'p': # if we are in the prosperous state
        # determine the agent's choice
        choice = self.Aset[a]
```

```
        if choice == 'c':  # cautious
            transitionprob = 0
        else:  # risky
            transitionprob = self.pc

        if self.Sset[s_] == 'd': # if we transitioned to the degraded state
            return transitionprob  # that is our transition probability
        else:  # back to the prosperous state
            return 1 - transitionprob

    else:  # if we are in the degraded state
        # determine the agent's choice
        choice = self.Aset[a]
        if choice == 'c':  # cautious
            transitionprob = self.pr
        else:  # risky
            transitionprob = 0

        if self.Sset[s_] == 'p': # if we transitioned to the prosperous state
            return transitionprob  # that is our transition probability
        else:  # back to the prosperous state
            return 1 - transitionprob
```

```
 #| export
@patch
def RewardTensor(self:RiskRewardDilemma):
    """Get the Reward Tensor R[i,s,a1,...,aN,s']."""
    dim = np.concatenate(([self.N],
                          [self.Z],
                          [self.M],
                          [self.Z]))
    Risas = np.zeros(dim)

    for index, _ in np.ndenumerate(Risas):
        Risas[index] = self._reward(index[0], index[1], index[2],
                                    index[3])
    return Risas

@patch
def _reward(self:RiskRewardDilemma,
            i:int, # the agent index
            s:int, # the state index
            a:int, # the action index
            s_:int  # the next-state index
            ) -> float:  # reward value
    """
    Returns the reward value for agent `i` in current state `s`, under joint action
 ↪  `jA`, when transitioning to next state `s_`.
    """
    if self.Sset[s] == 'd' or self.Sset[s_] == 'd':  # if either current or next
     ↪  state is degraded
        # determine the agent's choice
```

```
        choice = self.Aset[a]
        if choice == 'c':  # cautious
            reward = self.rd
        else:  # risky
            reward = self.rdr
        return reward

    else:  # if current and next state are prosperous
        # determine the agent's choice
        choice = self.Aset[a]
        if choice == 'c':  # cautious
            reward = self.rc
        else:  # risky
            reward = self.rr
        return reward
```

```
#| export
@patch
def id(self:RiskRewardDilemma):
    """
    Returns id string of environment
    """
    # Default
    pc = self.pc
    pr = self.pr
    rc = self.rc
    rr = self.rr
    rd = self.rd

    id = f"{self.__class__.__name__}_"+\
        f"{str(pc)}_{str(pr)}_{str(rc)}_{str(rr)}_{str(rd)}"
    return id
```

```
#| export
@patch
def FinalStates(self:RiskRewardDilemma):
    """Default final states: no final states"""
    return np.zeros(self.Z, dtype=int)
```

#### 8.1 Example

We use the following risk-reward dilemma, which we also implement below.

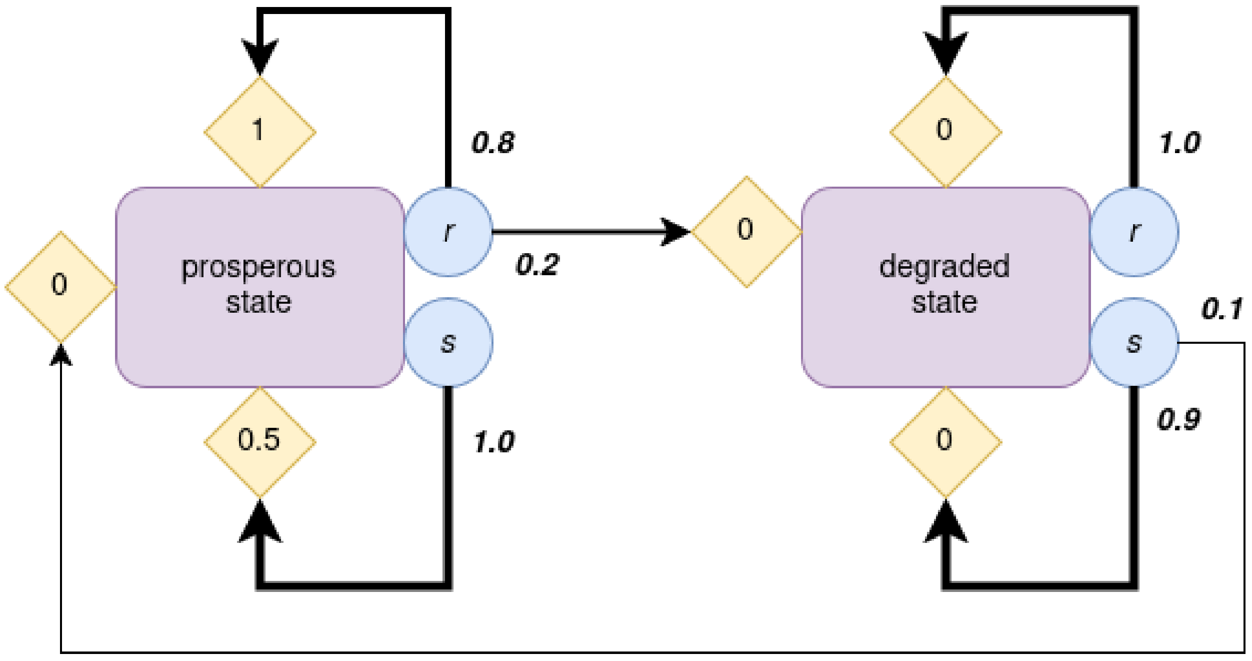

Figure 8.2: Risk-reward dilemma with specified values.

```
# Define environment
rrd = RiskRewardDilemma(pc=0.2, pr=0.1, rc=0.5, rr=1, rd=0)
```

Now, we define an agent-environment interface comprised of an actor-critic agent playing the aforementioned task. This agent has a learning rate $\alpha = 0.1$, and a discount factor $\gamma = 0.9$.

```
# Agent-environment interface
aei = stratAC(rrd,
              learning_rates=0.1,
              discount_factors=0.9,
              use_prefactor=True)
```

```
WARNING:2025-12-21 19:08:54,422:jax._src.xla_bridge:794: An NVIDIA GPU may be present on this 
enabled jaxlib is not installed. Falling back to cpu.
```

We plot the agent's deterministic trajectories in phase space on the flow plot below:

```
# Figure
x = ([0], [0], [0])  # (i,s,a) to plot along the x-axis
y = ([0], [1], [0])  # (i,s,a) to plot along the y-axis

# Plot flow plot
fig, ax1 = plt.subplots(1, 1, figsize=(4, 4))
fp.plot_strategy_flow(aei, x, y, flowarrow_points = np.linspace(0.01 ,0.99, 9),
                      NrRandom=16, axes=ax1)
# Sample trajectory
Xinit = np.ones((1,2,2)) * 0.4
Xinit[:,1,0] = 0.2
```

```
Xinit[:,:,1] = 1 - Xinit[:,:,0]  # initial strategy
trj, fpr = aei.trajectory(Xinit, Tmax=5000, tolerance=1e-6)  # compute traj
fp.plot_trajectories([trj], x, y, fprs=[fpr], axes=ax1, cols=['orange'])
ax1.set_title("Risk-reward dilemma")
ax1.set_xlabel("prob. safe action in prosp. state")
ax1.set_ylabel("prob. safe action in deg. state")
plt.show()
```

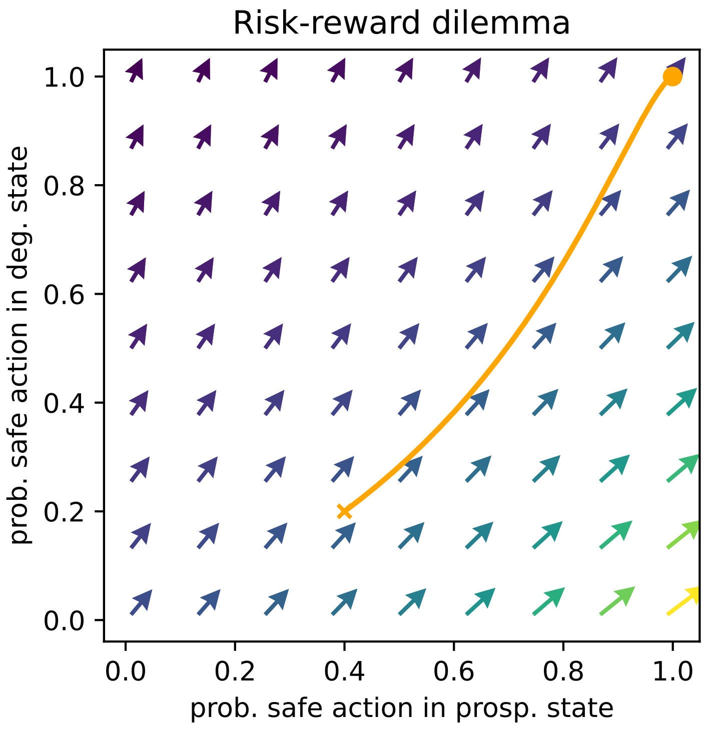

```
#| hide
import nbdev; nbdev.nbdev_export()
```

### Part III

### Utilities

In this section, we introduce tools for plotting and computing results. Chapter 9 contains functions to plot deterministic trajectories of the agents' strategies from given initial conditions. Chapter 10 contains functions to plot the derivative of agents' strategies against the strategies themselves. Finally, Chapter 11 contains specific plotting functions we use to generate the plots in our article.

### 9 Trajectories

This tool is particularly useful to visualize convergence and deterministic trajectories along one dimension. In particular, it is suited to environments with only one agent and/or one state, such as a two-armed bandit task, that we use below as an example.

First, we import what we need:

```
#| default_exp Utils.Trajectories1D
```

```
#| hide
# Imports for the nbdev development environment
import nbdev
from nbdev.showdoc import *
from fastcore.test import *
```

```
#| export
import numpy as np
import matplotlib as mpl
import matplotlib.pyplot as plt
import itertools as it

from collections.abc import Callable

from pyDOE import lhs

from fastcore.utils import *
```

```
# imports for testing
from pyDDRL.Environments.TwoArmedBandit import TwoArmedBandit
from pyCRLD.Agents.Base import abase
from pyCRLD.Agents.StrategySARSA import stratSARSA
```

#### 9.1 Implementation

```
#| export
def trajectories_1D(mae,  # CRLD multi-agent environment object
                    x:tuple,  # which phase space axes to plot along x axes
                    res,  # specify range & resolution
                    NrRandom:int=3,  # how many random (in the other dimensions)
↪  stratgies for averaging
                    traj_len:int=500,  # trajectory length
                    # col:str='LEN',  # color indicates either strength of flow via
                     ↪  colormap, otherwise a fixed color name
```

```
                    cmap='viridis',  # Colormap
                    # kind='quiver+samples', # Kind of plot: "streamplot",
                     ↪  "quiver+samples", "quiver", ...
                    # sf=0.5,  # Scale factor for quiver arrows
                    # lw=1.0,  # Line width for streamplot
                    # dens=0.75,  # Density for streamplot
                    acts=None,  # Action descriptions
                    conds=None,  # Conditions descriptions
                    axes:Iterable=None,  # Axes to plot into
                    verbose=False,  # shall I talk to you while working?
                    ):
"""
Create a flow plot in strategy space in one dimension.
"""
# Checks and balances
# xlens, amx, lens = _checks_and_balances(x, y)

# Fig and Axes
if axes==None:
    states = mae.Z
    if mae.Z == 1:
        fig, ax = plt.subplots(1, states, figsize=(states * 3.2, 2.8))
        axes = [ax]
    else:
        fig, axes = plt.subplots(1, states, figsize=(states * 3.2, 2.8))
else:
    axes = axes

cmap = mpl.cm.get_cmap(cmap)
ext = np.linspace(0, 1, len(res))
colors = cmap(tuple(ext))

# x axis
xs = np.arange(0, traj_len)

# The Plots
for i in range(len(x[1])): # go through each plot sequentially
    xinds = (x[0][0], x[1][i], x[2][0])  # fix this later
    # obtain the data to plot
    trajs = _data_to_plot(mae, res, xinds, traj_len, NrRandom,
                          phasespace_items=_strategies, verbose=verbose)
    for j in range(len(trajs[i, :, 0])):
        # do the plot
        color = colors[j]
        axes[i].plot(xs, trajs[i, j, :], color=color)

# Decorations
# lens = max(xlens)
if acts is None:
    acts = [f'act.{i}' for i in range(mae.M)]
    acts = mae.env.Aset
if conds is None:
    conds = [f'state {i}' for i in range(mae.Z)]
```

```
    for i in range(len(x[1])):
        axes[i].set_title(conds[i])
        axes[i].set_xlabel("step")
    axes[0].set_ylabel("prob. act. "+str(xinds[2]))

    return axes
```

```
#| export
def _data_to_plot(mae,  # CRLD multi-agent environment object
                 res:Iterable,  # range & resolution of flow arrows
                 xinds:tuple,  # of indices of the phase space object to plot along
↪  the x axis
                 traj_len:int,  # trajectory len
                 NrRandom:int,  # how many random (in the other dimensions)
↪  stratgies for averaging
                 # difffunc:Callable,  # to compute which kind of arrows to plot
                  ↪  (RPE or dX)
                 phasespace_items:Callable,  # to obtain phase space items for one
↪  ax plot point
                 verbose=False  # shall I talk to you while working?
                 ):

    l = len(res)
    trajs = np.zeros((mae.Z, l, traj_len))

    for i in range(mae.Z):
        for j, xval in enumerate(res):
            Xs = _strategies(mae, xinds, xval, NrRandom)  # generate random
↪  strategies to average over
            xtrajs = np.zeros((NrRandom, traj_len))
            for k, xk in enumerate(Xs):
                traj, fpreached = mae.trajectory(xk, traj_len) # compute trajectory
                xtrajs[k, :] = traj[:, xinds[0], i, xinds[2]]  # compute trajectory
            trajs[i, j, :] = np.mean(xtrajs, axis=0)  # add trajectory

    return trajs
```

```
#| export
def _strategies(mae,  # CRLD multi-agent environment object
               xinds:tuple,  # of indices of the phase space item to plot along the
↪  x axis
               xval:float,  # the value of the phase space item to plot along the x
↪  axis
               NrRandom,  # how many random (in the other dimensions) stratgies for
↪  averaging
              ) -> np.ndarray: # Array of joint strategies
    """
    Creates strategies (as a particular type of phase space item) for one ax plot
↪  point.
    All strategies have value `xval` at the `xinds` index and value `yval` at the
↪  `yinds`.
    """
```

```
    N, C, M = mae.N, mae.Q, mae.M  # Number of agents, conditions, actions
    # Xs = np.random.rand(NrRandom, N, C, M)  # random policies
    Xs = lhs(N*C*M, NrRandom).reshape(NrRandom, N, C, M)  # using latin hypercube
↪   sampling
    Xs = Xs / Xs.sum(axis=-1, keepdims=True)  # properly normalised

    xi, xc, xa = xinds; xa_ = tuple(set(range(M)) - set([xa]))

    Xs[:, xi, xc, xa] = xval  # setting x and y values

    # normalisation
    Xs[:,xi,xc,xa_] = (1-Xs[0, xi, xc, xa]) * Xs[:,xi,xc,xa_]\
        / np.sum(Xs[:,xi,xc,xa_], axis=-1, keepdims=True)

    return Xs
```

#### 9.2 Example with a two-armed bandit task

Let's consider a two-armed bandit task with $p_1 = 0.7$ and $p_2 = 0.3$.

```
# Define environment
xs = np.array([[-1, 1],
               [-1, 1]])
ps = np.array([[0.3, 0.7],
               [0.7, 0.3]])
tab = TwoArmedBandit(disttype='discrete', xs=xs, ps=ps)
```

Now, we define an agent-environment interface comprised of a SARSA agent playing the aforementioned task. This agent has a learning rate $\alpha = 0.1$, a discount factor $\gamma = 0$, and a choice intensity $\beta = 4$.

```
# Agent-environment interface
aei = stratSARSA(tab,
                 learning_rates=0.1,
                 discount_factors=0,
                 choice_intensities=4.,
                 use_prefactor=False)
```

```
WARNING:2025-12-21 19:09:19,752:jax._src.xla_bridge:794: An NVIDIA GPU may be present on this
enabled jaxlib is not installed. Falling back to cpu.
```

```
x = ([0], [0], [0])  # (i,s,a) to plot along the x-axis
res = np.linspace(0, 1., 11)  # initial strategies
res[0] = 0.0001
res[-1] = 0.9999
```

Now plotting the trajectories:

```
fig, ax = plt.subplots(1, 1, figsize=(4, 3))
trajectories_1D(aei, x, res, NrRandom=1, traj_len=70, axes=[ax])
ax.set_title("deterministic trajectories")
ax.set_xlabel("time step "+r'$t$')
ax.set_ylabel("probability to choose arm 1")
plt.show()
```

```
/tmp/ipykernel_10416/1852574533.py:35: MatplotlibDeprecationWarning: The get_cmap function was
  cmap = mpl.cm.get_cmap(cmap)
```

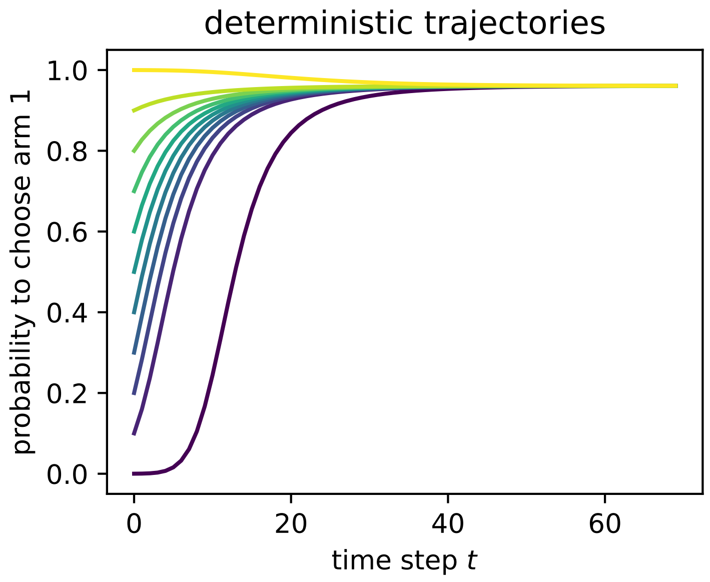

```
#| hide
import nbdev; nbdev.nbdev_export()
```

### 10 Phase plot

A phase plot represents the derivative $dX$ of a strategy $X$ against this same strategy $X$. The intersection of the resulting curve with the $x$-axis is a fixed point of the dynamics.

This tool is particularly useful to visualize convergence and equilibria along one dimension. In particular, it is suited to environments with only one agent and/or one state, such as a two-armed bandit task, that we use below as an example.

First, we import what we need:

```
#| default_exp Utils.PhasePlot
```

```
#| hide
# Imports for the nbdev development environment
import nbdev
from nbdev.showdoc import *
from fastcore.test import *
```

```
#| export
import numpy as np
import matplotlib as mpl
import matplotlib.pyplot as plt
import itertools as it

from collections.abc import Callable

from pyDOE import lhs

from fastcore.utils import *
```

```
# imports for testing
from pyDDRL.Environments.TwoArmedBandit import TwoArmedBandit
from pyCRLD.Agents.Base import abase
from pyCRLD.Agents.StrategySARSA import stratSARSA
```

#### 10.1 Implementation

```
#| export
def phaseplot(mae,  # CRLD multi-agent environment object
              x:tuple,  # which phase space axes to plot along x axes
              res,  # specify range & resolution
              NrRandom:int=3,  # how many random (in the other dimensions) stratgies
↪ for averaging
```

```
            traj_len:int=500,  # trajectory length
            # col:str='LEN',  # color indicates either strength of flow via
             ↪  colormap, otherwise a fixed color name
            cmap='viridis',  # Colormap
            # kind='quiver+samples', # Kind of plot: "streamplot",
             ↪  "quiver+samples", "quiver", ...
            # sf=0.5,  # Scale factor for quiver arrows
            # lw=1.0,  # Line width for streamplot
            # dens=0.75,  # Density for streamplot
            acts=None,  # Action descriptions
            conds=None,  # Conditions descriptions
            axes:Iterable=None,  # Axes to plot into
            verbose=False,  # shall I talk to you while working?
            ):
    """
    Create a phase plot in strategy space.
    """

    # Fig and Axes
    if axes == None:
        states = mae.Z
        if mae.Z == 1:
            fig, ax = plt.subplots(1, states, figsize=(states * 3.8, 2.8))
            axes = [ax]
        else:
            fig, axes = plt.subplots(1, states, figsize=(states * 3.8, 2.8))
    else:
        axes = axes

    # cmap = mpl.cm.get_cmap(cmap)
    ext = np.linspace(0, 1, len(res))

    # x axis
    xs = res
    zero = np.zeros((len(xs)))

    # The Plots
    for i in range(len(x[1])): # go through each plot sequentially
        xinds = (x[0][0], x[1][i], x[2][0])  # fix this later
        # obtain the data to plot
        RPEs, dXs = _data_to_plot(mae, res, xinds, traj_len, NrRandom,
                                  phasespace_items=_strategies, verbose=verbose)
        axes[i].plot(xs, RPEs[i, :], color="blue")
        axes[i].plot(xs, zero, color="grey", ls='dashed')
        twinax = axes[i].twinx()
        twinax.plot(xs, dXs[i, :], color="red")

        # Set default limits before aligning
        axes[i].set_ylim(min(RPEs[i, :]), max(RPEs[i, :]))  # Primary y-axis
        twinax.set_ylim(min(dXs[i, :]), max(dXs[i, :]))     # Secondary y-axis
        if i==len(x[1])-1:
            twinax.set_ylabel("dX", color="red")
```

```
        # Align the zero levels of both axes
        align_zero_limits(axes[i], twinax)

    # Decorations
    # lens = max(xlens)
    if acts is None:
        acts = [f'act.{i}' for i in range(mae.M)]
        acts = mae.env.Aset
    if conds is None:
        conds = [f'state {i}' for i in range(mae.Z)]
    for i in range(len(x[1])):
        axes[i].set_title(conds[i])
        axes[i].set_xlabel("prob. act. "+str(xinds[2]))
    axes[0].set_ylabel("RPE", color="blue")

    # fig.tight_layout()

    return axes
```

```
 #| export
# Function to adjust y-axis limits to ensure 0 is aligned
def align_zero_limits(ax1, ax2, padding_factor=0.05):
    # Get the limits of both y-axes
    ax1_min, ax1_max = ax1.get_ylim()
    ax2_min, ax2_max = ax2.get_ylim()

    # Compute ranges and negative-to-positive ratios
    ax1_range = max(ax1_max, 0) - min(ax1_min, 0)
    ax2_range = max(ax2_max, 0) - min(ax2_min, 0)

    ax1_neg_to_pos_ratio = -ax1_min / ax1_max if ax1_max != 0 else 1
    ax2_neg_to_pos_ratio = -ax2_min / ax2_max if ax2_max != 0 else 1

    # Adjust limits proportionally to keep 0 aligned
    if ax1_neg_to_pos_ratio > ax2_neg_to_pos_ratio:
        # Adjust ax2 to match ax1's ratio
        new_ax2_min = -ax2_max * ax1_neg_to_pos_ratio
        ax2.set_ylim(new_ax2_min, ax2_max)
        ax1.set_ylim(ax1_min )
    else:
        # Adjust ax1 to match ax2's ratio
        new_ax1_min = -ax1_max * ax2_neg_to_pos_ratio
        ax1.set_ylim(new_ax1_min, ax1_max)

    # Add padding (white space) to both axes
    ax1_min, ax1_max = ax1.get_ylim()
    ax2_min, ax2_max = ax2.get_ylim()

    # Extend the limits by a percentage (padding_factor)
    ax1_padding = (ax1_max - ax1_min) * padding_factor
    ax2_padding = (ax2_max - ax2_min) * padding_factor
```

```
    # Set the new padded limits
    ax1.set_ylim(ax1_min - ax1_padding, ax1_max + ax1_padding)
    ax2.set_ylim(ax2_min - ax2_padding, ax2_max + ax2_padding)
```

```
#| export
def _data_to_plot(mae,  # CRLD multi-agent environment object
                  res:Iterable,  # range & resolution of flow arrows
                  xinds:tuple,  # of indices of the phase space object to plot along
↪ the x axis
                  traj_len:int,  # trajectory len
                  NrRandom:int,  # how many random (in the other dimensions)
↪ stratgies for averaging
                  # difffunc:Callable,  # to compute which kind of arrows to plot
                   ↪ (RPE or dX)
                  phasespace_items:Callable,  # to obtain phase space items for one
↪ ax plot point
                  verbose=False  # shall I talk to you while working?
                  ):

    l = len(res)
    RPEs = np.zeros((mae.Z, l))
    dXs = np.zeros((mae.Z, l))

    for i in range(mae.Z):
        for j, xval in enumerate(res):
            Xs = _strategies(mae, xinds, xval, NrRandom)  # generate random
↪ strategies to average over
            RPE_ = np.zeros((NrRandom))
            for k, xk in enumerate(Xs):
                prederror = mae.TDerror(xk, norm=True) # compute pred error
                RPE_[k] = prederror[xinds[0], i, xinds[2]]  # pred error for indices
↪ of interest
            RPEs[i, j] = np.mean(RPE_)  # add trajectory
            dX_ = np.array(_dXisa_s(Xs, mae))  # compute change in Xisa
            dX = np.mean(dX_, axis=0)  # average over random strategies
            dXs[i, j] = dX[xinds[0], i, xinds[2]]  # dX for indices of interest

    return RPEs, dXs
```

```
#| export
def _dXisa_s(Xisa_s:Iterable, # of joint strategies `Xisa`
             mae  # CRLD multi-agent environment object
             ) -> np.ndarray:  # joint strategy differences
    """Compute `Xisa`(t+1)-`Xisa`(t) for all `Xisa_s`."""
    return np.array([mae.step(Xisa)[0] - Xisa for Xisa in Xisa_s])
```

```
#| export
def _strategies(mae,  # CRLD multi-agent environment object
                xinds:tuple,  # of indices of the phase space item to plot along the
↪ x axis
                xval:float,  # the value of the phase space item to plot along the x
↪ axis
```

```
                NrRandom,  # how many random (in the other dimensions) stratgies for
↪   averaging
               ) -> np.ndarray: # Array of joint strategies
    """
    Creates strategies (as a particular type of phase space item) for one ax plot
↪   point.
    All strategies have value `xval` at the `xinds` index and value `yval` at the
↪   `yinds`.
    """
    N, C, M = mae.N, mae.Q, mae.M  # Number of agents, conditions, actions
    # Xs = np.random.rand(NrRandom, N, C, M)  # random policies
    Xs = lhs(N*C*M, NrRandom).reshape(NrRandom, N, C, M)  # using latin hypercube
↪   sampling
    Xs = Xs / Xs.sum(axis=-1, keepdims=True)  # properly normalised

    xi, xc, xa = xinds; xa_ = tuple(set(range(M)) - set([xa]))

    Xs[:, xi, xc, xa] = xval  # setting x and y values

    # normalisation
    Xs[:,xi,xc,xa_] = (1-Xs[0, xi, xc, xa]) * Xs[:,xi,xc,xa_]\
        / np.sum(Xs[:,xi,xc,xa_], axis=-1, keepdims=True)

    return Xs
```

#### 10.2 Example with a two-armed bandit task

Let's consider a two-armed bandit task with $p_1 = 0.7$ and $p_2 = 0.3$.

```
# Define environment
xs = np.array([[-1, 1],
               [-1, 1]])
ps = np.array([[0.3, 0.7],
               [0.7, 0.3]])
tab = TwoArmedBandit(disttype='discrete', xs=xs, ps=ps)
```

Now, we define an agent-environment interface comprised of a SARSA agent playing the aforementioned task. This agent has a learning rate $\alpha = 0.1$, a discount factor $\gamma = 0$, and a choice intensity $\beta = 4$.

```
# Agent-environment interface
aei = stratSARSA(tab,
                 learning_rates=0.1,
                 discount_factors=0,
                 choice_intensities=4.,
                 use_prefactor=False)
```

```
WARNING:2025-12-21 19:09:34,562:jax._src.xla_bridge:794: An NVIDIA GPU may be present on this 
enabled jaxlib is not installed. Falling back to cpu.
```

```
x = ([0], [0], [0])  # (i,s,a) to plot along the x-axis
res_phase = np.linspace(0, 1., 51)  # resolution for phase plot
res_phase[0] = 0.001
res_phase[-1] = 0.999
```

Now making the phase plot:

```
fig, ax = plt.subplots(1, 1, figsize=(4, 3))
phaseplot(aei, x, res_phase, NrRandom=1, axes=[ax])
ax.set_xlabel("strategy "+r'$X$')
ax.set_title("phase plot")
plt.show()
```

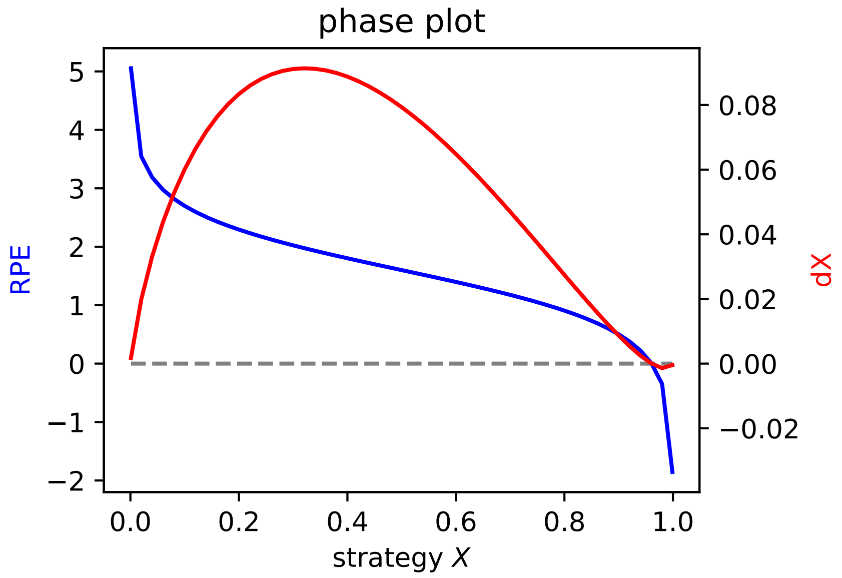

```
 #| hide
import nbdev; nbdev.nbdev_export()
```

### 11 Plotting tools

Here, we introduce the more elaborate plotting tools we need to produce figures for Section **?@sec-results** . First, we import what we need:

```
#| default_exp Utils.PlottingTools
```

```
#| hide
# Imports for the nbdev development environment
import nbdev
from nbdev.showdoc import *
from fastcore.test import *
```

```
#| export
import numpy as np
import matplotlib as mpl
import matplotlib.pyplot as plt
import itertools as it
import seaborn as sns

from collections.abc import Callable

from pyDOE import lhs
import jax.numpy as jnp

from fastcore.utils import *
```

```
#| export
# Agents
from pyCRLD.Agents.Base import abase
from pyCRLD.Agents.StrategySARSA import stratSARSA
from pyCRLD.Agents.StrategyActorCritic import stratAC

from pyDDRL.Agents.DDRLBase import DDRLBase
from pyDDRL.Agents.DDRLSarsa import DDRLSarsa
from pyDDRL.Agents.DDRLActorCritic import DDRLActorCritic

# Environments
from pyDDRL.Environments.TwoArmedBandit import TwoArmedBandit
```

```
#| export
# Utilities
from pyCRLD.Utils.Helpers import *
from pyDDRL.Utils.Trajectories1D import trajectories_1D
from pyDDRL.Utils.PhasePlot import phaseplot, _data_to_plot, _strategies, _dXisa_s
from pyCRLD.Utils import FlowPlot as fp
```

#### 11.1 Implementation

##### 11.1.1 Compare phase plots, trajectories and mean rewards for different optimism levels

```
#| export
## Functions to generate phase plots + trajs + mean rewards

# Obtain dX as a function of X for phase plots
# Also works when N=2 or Z=2
def obtain_dXs(aei,  # agent-environment interface
               res,  # range & resolution of strategies
               xinds:tuple,  # of indices of the phase space object to plot along
↪  the x axis
               traj_len:int,  # trajectory len
               otherXs=None,  # when N=2 or Z=2: values of X in 2nd condition
               ):

    l = len(res)
    dXs = np.zeros((aei.Z, l))  # will store the dXs

    for i in range(aei.Z):

        for j, xval in enumerate(res):  # loop over Xs

            if (aei.N > 1) or (aei.Z > 1): # 2 states or 2 actions
                xval_ = otherXs[j]
            else:
                xval_ = None

            X = obtain_Xs(aei, xinds, xval, xval_)  # create X with values along all
↪  dims
            prederror = aei.TDerror(X, norm=True)  # compute pred error
            dX = aei.step(X)[0] - X  # compute dX
            dXs[i, j] = dX[xinds[0], i, xinds[2]][0]  # dX for i,s,a of interest

    return dXs

def obtain_Xs(aei,  # CRLD multi-agent environment object
              xinds:tuple,  # of indices of the phase space item to plot along the x
↪  axis
              xval:float,  # the value of the phase space item to plot along the x
↪  axis
              xval_:float  # the value of X in the 2nd condition
              ):

    N, C, M = aei.N, aei.Z, aei.M  # Number of agents, states, actions

    Xs = np.zeros((N, C, M))  # will store the Xs

    xi, xc, xa = xinds  # plotted agent, state, action
    xa_ = tuple(set(range(M)) - set(xa))  # other action
```

```
    Xs[xi, xc, xa] = xval  # setting x value

    # probability of taking other action
    Xs[xi,xc,xa_] = 1-Xs[xi, xc, xa]

    if C > 1:  # more than one state
        xc_ = tuple(set(range(C)) - set(xc))  # other state
        Xs[xi, xc_, xa] = xval_  # strategy in state 2 for action 1
        Xs[xi, xc_, xa_] = 1 - Xs[xi, xc_, xa]  # strategy in state 2 for action 2
    if N > 1:  # more than 1 agent
        xi_ = tuple(set(range(N)) - set(xi))  # other agent
        Xs[xi_, xc, xa] = xval_  # other agent's strategy for action 1
        Xs[xi_, xc, xa_] = 1 - Xs[xi_, xc, xa]  # other agent's strat for action 2

    return Xs

# Compute X traj and reward traj
def reward_traj(aei,  # CRLD multi-agent environment object
                X:tuple,  # strategy
                i,  # agent
                traj_len:int=500,  # trajectory length
                ):

    r_traj = np.zeros((traj_len))  # will store rewards
    # Compute X trajectory
    traj, fpreached = aei.trajectory(X, traj_len)
    # Compute reward trajectory
    for t in range(traj_len):
        r_traj[t] = aei.Ri(traj[t,...])[i]

    return traj, r_traj

# Plot phase plot, X traj and reward traj for various levels of optimism/pessimism
def compare_levels(env,  # environment
                   level_list,  # list of levels
                   gamma,  # discount factor
                   beta,  # choice intensity
                   Rmin,  # min reward perceived
                   Rmax,  # max reward perceived
                   xinds:tuple,  # which phase space axes to plot along x axes
                   res,  # specify range & resolution
                   algo='SARSA',  # specify algorithm: 'SARSA' or 'AC'
                   nrb:int=29,  # number of neurons
                   discsig:float=0.05,  # discretization sigma
                   lr:float=0.1,  # learning rate
                   traj_len:int=500,  # trajectory length
                   otherXs=None,  # if 2nd condition: values of X along this
 ↪  condition
                   cmap='viridis',  # colormap
                   save=False,  # save fig?
                   savename=None,  # saved figure's name
                   axes=None,  # Axes to plot into
```

```
                ):

    # Fig and Axes
    # Style
    sns.set_style("whitegrid")
    sns.set_context("paper", font_scale=1.5, rc={"lines.linewidth": 2})

    # Subplots: phase plot, X traj, reward traj
    fig, (ax1, ax2, ax3) = plt.subplots(1, 3, figsize=(4.5 * 3.8, 3.8))

    # Colormap for the different lines
    cmap = mpl.colormaps[cmap]
    ext = np.linspace(0, 1, len(level_list))  # extent of colormap
    colors = cmap(tuple(ext))
    # Neutral agent is represented with a black line
    colors[int(len(level_list)/2)] = [0, 0, 0, 1]

    # Zero line
    zero = np.zeros((len(res)))

    # Loop over optimism/pessimism levels
    for j, level in enumerate(level_list):

        # Define agent-environment interface
        if algo=='SARSA':
            aei = DDRLSarsa(env, nr_reward_bins=nrb, discretization_sigma=discsig,
                            learning_rates=lr, discount_factors=gamma,
                            choice_intensities=beta, Rmin=Rmin, Rmax=Rmax,
                            method="weights", wU=level, use_prefactor=True)
        elif algo=='AC':
            aei = DDRLActorCritic(env, nr_reward_bins=nrb,
 ↪  discretization_sigma=discsig,
                                  learning_rates=lr, discount_factors=gamma,
                                  Rmin=Rmin, Rmax=Rmax,
                                  method="weights", wU=level, use_prefactor=True)

        # Obtain the dXs for y-axis of phase plot
        dXs = obtain_dXs(aei, res, xinds, traj_len, otherXs)

        # Phase plot
        ax1.plot(res, dXs[0, :], color=colors[j])
        ax1.plot(res, zero, color="grey", ls='dashed')  # plot zero line

        # Obtain X traj and reward traj
        X0 = np.ones((aei.N, aei.Z, aei.M)) * 0.5  # init random strategy
        X_traj, r_traj = reward_traj(aei, X0, 0, traj_len=traj_len)

        # X traj plot
        ax2.plot(np.arange(0, traj_len), X_traj[:,0,0,0],
                 color=colors[j], label=r'$w_{U}=$'+str(level))

        # Reward plot
```

```
        ax3.plot(np.arange(0, traj_len), r_traj,
                 color=colors[j], label=r'$w_{U}=$'+str(level))

    # Decorations
    ax1.set_ylabel(r"$dX$")
    ax1.set_xlabel("strategy "+r'$X$')
    ax1.set_title('phase plot')
    ax2.set_xlabel("time step "+r'$t$')
    ax2.set_ylabel("strategy "+r'$X^{0,0,0}(t)$')
    ax2.set_title('strategy over time')
    ax3.set_xlabel("time step "+r'$t$')
    ax3.set_ylabel("reward "+r'$R^{0}(t)$')
    ax3.set_title('mean reward over time')
    ax3.legend(bbox_to_anchor=(1.01, 1.01), title='weights '+r'$w_U$')
    fig.tight_layout()

    # Save figure
    if save == True:
        plt.savefig(savename, dpi=300, bbox_inches='tight')

# Plot phase plot, X traj and reward traj for various levels of optimism/pessimism
def compare_gammas(env,  # environment
                   level_list,  # list of levels
                   Rmin,  # min reward perceived
                   Rmax,  # max reward perceived
                   xinds:tuple,  # which phase space axes to plot along x axes
                   res,  # specify range & resolution
                   nrb:int=29,  # number of neurons
                   discsig:float=0.05,  # discretization sigma
                   lr:float=0.1,  # learning rate
                   traj_len:int=500,  # trajectory length
                   otherXs=None,  # if 2nd condition: values of X along this
 ↪  condition
                   cmap='viridis',  # colormap
                   save=False,  # save fig?
                   savename=None,  # saved figure's name
                   axes=None,  # Axes to plot into
                   ):

    # Fig and Axes
    # Style
    sns.set_style("whitegrid")
    sns.set_context("paper", font_scale=1.5, rc={"lines.linewidth": 2})

    # Subplots: phase plot, X traj, reward traj
    fig, (ax1, ax2) = plt.subplots(1, 2, figsize=(12, 3.8))

    # Colormap for the different lines
    cmap = mpl.colormaps[cmap]
    ext = np.linspace(0, 1, len(level_list))  # extent of colormap
    colors = cmap(tuple(ext))
```

```
    # Zero line
    zero = np.zeros((len(res)))

    # Loop over optimism/pessimism levels
    for j, level in enumerate(level_list):

        # Define agent-environment interface
        aei = DDRLActorCritic(env, nr_reward_bins=nrb, discretization_sigma=discsig,
↪
                              learning_rates=lr, discount_factors=level,
                              Rmin=Rmin, Rmax=Rmax,
                              method="weights", wU=1, use_prefactor=True)
        mae = stratAC(env, learning_rates=lr, discount_factors=level,
                      use_prefactor=True)

        # Obtain the dXs for y-axis of phase plot
        dXs_aei = obtain_dXs(aei, res, xinds, traj_len, otherXs)
        dXs_mae = obtain_dXs(mae, res, xinds, traj_len, otherXs)

        # Phase plot
        ax1.plot(res, dXs_mae[0, :], color=colors[j])
        ax1.plot(res, zero, color="grey", ls='dashed')  # plot zero line
        ax2.plot(res, dXs_aei[0, :], color=colors[j],
                 label=r'$\gamma=$'+str(level))
        ax2.plot(res, zero, color="grey", ls='dashed')  # plot zero line

    # Decorations
    ax1.set_ylabel(r"$dX^{safe}$", fontsize=18)
    ax1.set_xlabel("strategy "+r'$X^{safe}$', fontsize=18)
    ax1.set_title('phase plots for a non-distributional agent', fontsize=16)
    ax2.set_xlabel("strategy "+r'$X$', fontsize=18)
    ax2.set_title('phase plots for a distributional agent', fontsize=16)
    ax2.legend(bbox_to_anchor=(1.01, 1.01), title='discount factor '+r'$\gamma$',
↪   fontsize=16)
    fig.tight_layout()

    # Save figure
    if save == True:
        plt.savefig(savename, dpi=300, bbox_inches='tight')
```

##### 11.1.2 Comparison between asymmetric RL and DDRL

```
 #| export
## Functions for comparisons between DDRL biasing methods and asymmetric RL

# Compute Q-value gap at steady state
def deltaQinf(p0,  # reward prob of arm 1
              p1,  # reward prob of arm 2
```

```
            b):  # bias strength
    return ((p0 * b - (1 - p0)) / (p0 * b + (1 - p0))) - ((p1 * b - (1 - p1)) / (p1
     ↪  * b + (1 - p1)))

# Compute prob of choosing most rewarding arm at steady state
def Xinf(dQ,  # Q-value gap
         beta):  # choice intensity
    return 1 / (1 + np.exp(beta * (-dQ)))

# Compute final strategies for quantile, expectile, weights, and asymRL methods
def compute_comparisons(p0,  # reward prob of arm 1
                        p1,  # reward prob of arm 2
                        b_ratios,  # bias strengths for asymRL
                        q_list,  # quantiles
                        e_list,  # expectiles
                        w_list,  # weights
                        gamma,  # discount factor
                        beta,  # choice intensity
                        Rmin,  # min reward perceived
                        Rmax,  # max reward perceived
                        nrb:int=29,  # number of neurons
                        discsig:float=0.05,  # discretization sigma
                        lr:float=0.1,  # learning rate
                        axes=None,  # axes to plot into
                        ):

    # Create 2-armed bandit
    xs = np.array([[-1, 1],
                   [-1, 1]])
    ps = np.array([[1-p0, p0],
                   [1-p1, p1]])
    env = TwoArmedBandit(disttype='discrete', xs=xs, ps=ps)

    # Compute asymRL Xinf for each bias strength
    b_dQs = deltaQinf(p0, p1, b_ratios)  # Q-value gaps
    b_xinf = Xinf(b_dQs, beta)  # final strategies

    # Compute quantile Xinf for each quantile
    q_xinf = np.zeros((len(q_list)))
    for i, q in enumerate(q_list):
        # Define agent-env interface
        aei = DDRLSarsa(env, nr_reward_bins=nrb, discretization_sigma=discsig,
                        learning_rates=lr, discount_factors=gamma,
                        choice_intensities=beta, Rmin=Rmin, Rmax=Rmax,
                        method='quantile', tau=q, use_prefactor=False)
        # Initial strategy
        X = 0.5 * np.ones((aei.N, aei.Z, aei.M))
        # Compute strategy trajectory
        traj, fpreached = aei.trajectory(X, 150)
        # Take the last X^{0,0,0} of the trajectory
        q_xinf[i] = traj[-1, 0, 0, 0]

    # Compute expectile Xinf for each expectile
```

```
    e_xinf = np.zeros((len(e_list)))
    for i, e in enumerate(e_list):
        # Define agent-env interface
        aei = DDRLSarsa(env, nr_reward_bins=nrb, discretization_sigma=discsig,
                        learning_rates=lr, discount_factors=gamma,
                        choice_intensities=beta, Rmin=Rmin, Rmax=Rmax,
                        method='expectile', tau=e, use_prefactor=False)
        # Initial strategy
        X = 0.5 * np.ones((aei.N, aei.Z, aei.M))
        # Compute strategy trajectory
        traj, fpreached = aei.trajectory(X, 150)
        # Take the last X^{0,0,0} of the trajectory
        e_xinf[i] = traj[-1, 0, 0, 0]

    # Compute weights Xinf for each upper weight value
    w_xinf = np.zeros((len(w_list)))
    for i, w in enumerate(w_list):
        # Define agent-env interface
        aei = DDRLSarsa(env, nr_reward_bins=nrb, discretization_sigma=discsig,
                        learning_rates=lr, discount_factors=gamma,
                        choice_intensities=beta, Rmin=Rmin, Rmax=Rmax,
                        method='weights', wU=w, use_prefactor=False)
        # Initial strategy
        X = 0.5 * np.ones((aei.N, aei.Z, aei.M))
        # Compute strategy trajectory
        traj, fpreached = aei.trajectory(X, 150)
        # Take the last X^{0,0,0} of the trajectory
        w_xinf[i] = traj[-1, 0, 0, 0]

    return b_xinf, q_xinf, e_xinf, w_xinf

# Plot the final strategies
def plot_comparisons(b_ratios, q_list, e_list, w_list, b_xinf, q_xinf,
                     e_xinf, w_xinf, scarcity="poor", save=False,
                     savename=None):

    # Style
    sns.set_style("whitegrid")
    sns.set_context("paper", font_scale=1.5, rc={"lines.linewidth": 2})
    # Subplots: quantile, expectile, weights
    fig, (ax1, ax2, ax3) = plt.subplots(1, 3, figsize=(4.5 * 3.8, 4.5))
    axes = [ax1, ax2, ax3]
    # Twin axes and their properties
    ax11 = ax1.twiny()
    ax22 = ax2.twiny()
    ax33 = ax3.twiny()
    twin_axes = [ax11, ax22, ax33]
    ddrl_lists = [q_list, e_list, w_list]
    ddrl_xinfs = [q_xinf, e_xinf, w_xinf]
    colors = ['red', 'green', 'orange']
    labels = ['quantile', 'expectile', 'DDRL']
    xlabels = [r'$\tau$', r'$\tau$', 'weight '+r'$w_U$']
```

```python
# to find the bias strength of the neutral asymRL agent
ind = np.sum(b_ratios < 1)

# The plots
for i, ax in enumerate(axes):
    ax.set_xscale('log')  # log scale for x-axis
    # Plot Xinfs for asym RL
    ax.plot(b_ratios, b_xinf, '-+', mew=3, ms=8, color='blue', label="asym RL")
    # Plot neutral line
    ax.plot(b_ratios, np.repeat(b_xinf[ind], len(b_ratios)), '--', color='gray')
    # Plot Xinfs for DDRL method
    if i==1:
        axes[i].plot(b_ratios, ddrl_xinfs[i], '-o', color=colors[i],
                     label=labels[i])
    else:
        twin_axes[i].plot(ddrl_lists[i], ddrl_xinfs[i], '-o', color=colors[i],
                          label=labels[i])

# Instructions for specific axes
ax33.set_xscale('log')

# Decorations
for i, ax in enumerate(axes):
    if scarcity=="poor":
        ax.legend(loc='upper left')
    else:
        ax.legend(loc='upper right')
    ax.set_xlabel('bias strength '+r'$b$', color="blue")
    ax.xaxis.label.set_color('blue')
    ax.tick_params(axis='x', colors='blue')
    ax.spines['bottom'].set_color('blue')
    ax.spines['top'].set_color(colors[i])
    twin_axes[i].set_xlabel(xlabels[i], color=colors[i])
    twin_axes[i].xaxis.label.set_color(colors[i])
    twin_axes[i].tick_params(axis='x', colors=colors[i])
    twin_axes[i].spines['bottom'].set_color('blue')
    twin_axes[i].spines['top'].set_color(colors[i])

# Decorations for specific axes
ax1.set_ylabel('final strategy '+r'$X_{\infty}$')
# ax22.set_xticks(e_list)

if scarcity=="poor":
    ax11.legend(loc="center left")
    ax33.legend(loc='center left')
else:
    ax11.legend(loc='center right')
    ax33.legend(loc='center right')

ax1.set_title("asymmetric RL vs. quantiles")
ax2.set_title("asymmetric RL vs. expectiles")
ax3.set_title("asymmetric RL vs. DDRL in a poor environment")
```

```
    fig.tight_layout()

    # Save figure
    if save == True:
        plt.savefig(savename, dpi=300, bbox_inches='tight')
```

##### 11.1.3 Plot 3 flowplots with associated trajectories

```
 #|export
## Functions to plot 3 flowplots and associated trajectories

# Plot the flowplots together
def plot_3flowplots(aeis,  # list of aeis
                    x:tuple,  # which phase space axes to plot along x axes
                    y:tuple,  # which phase space axes to plot along y axes
                    NrRandom:int=4,  # number of random strategies to average over
                    Xinits=None,  # initial Xs to plot trajectories
                    decorate=False,  # whether to decorate the axes or not
                    titles=None,  # list of titles for 3 plots
                    xlabel=None,  # x-label
                    ylabel=None,  # y-label
                    cols=None,  # list of colors for trajectories
                    cmap='viridis',  # Colormap
                    axes=None,  # Axes to plot into
                    ):

    # Style
    sns.set_style("white")
    sns.set_context("paper", font_scale=1.25, rc={"lines.linewidth": 2})

    # Plot the flow plot
    for i, ax in enumerate(axes):
        aei = aeis[i]
        fp.plot_strategy_flow(aei, x, y, flowarrow_points = np.linspace(0.01 ,0.99,
↪   9),
                              use_RPEarrows=False, NrRandom=NrRandom, cmap=cmap,
↪   axes=ax)

        # Plot trajectory if initial strategies are specified
        if Xinits != None:
            trajs = []  # trajectories
            fprs = []  # are fixed points reached?
            for _, Xinit in enumerate(Xinits):
                trj, fpr = aei.trajectory(Xinit, Tmax=12000, tolerance=1e-6)
                trajs.append(trj)
                fprs.append(fpr)

            fp.plot_trajectories(trajs, x, y, fprs=fprs, axes=ax, cols=cols)
```

```
    # Add titles, xlabels and ylabels to subplots if needed
    if decorate==True:
        for i, ax in enumerate(axes):
            ax.set_xlabel(xlabel)
            ax.set_title(titles[i])

        axes[0].set_ylabel(ylabel)
```

##### 11.1.4 Plot separatrix on flowplots

```
#|export
def compile_strategy(p0c:float,  # cooperation probability of agent zero
                     p1c:float):  # cooperation probability of agent one
    strat = np.zeros((2, 1, 2))
    strat[0, 0, 0] = p0c
    strat[1, 0, 0] = p1c
    strat[0, 0, 1] = 1 - p0c
    strat[1, 0, 1] = 1 - p1c
    return strat

def plot_separatrix(aei, ax, x, y):
    # Add saddle node
    # by reversing the dynamics from two agents with identical strategies
    o = [0.5, 0.5]; o = compile_strategy(*o)
    for _ in range(1000): o, _ = aei.reverse_step(o)
    # Add separatrix
    o = o[:, 0, 0] - np.array([0, 0.0001])
    o1 = compile_strategy(*o)
    o2 = compile_strategy(*o[::-1])
    sep1=[]; sep2=[]
    for _ in range(1000): o1, _ = aei.reverse_step(o1); sep1.append(o1)
    for _ in range(1000): o2, _ = aei.reverse_step(o2); sep2.append(o2)
    fp.plot_trajectories([sep1, sep2], x=x, y=y, cols=['dimgray'], lws=[1],
                         lss=['--'], alphas=[0.95], axes=ax)
```

##### 11.1.5 Generate stability landscape

```
#|export
def fill_containers(env, Xrisk, Xsafe, discountfactors, weights):
    risky_optimal_data_container = np.zeros((discountfactors.size, weights.size, 2))
    risky_stable_container = np.zeros((discountfactors.size, weights.size, 2))
    safe_stable_container = np.zeros((discountfactors.size, weights.size, 2))
    for i, dcf in enumerate(discountfactors):
        for j, w in enumerate(weights):
            aei = DDRLActorCritic(env, nr_reward_bins=29, discretization_sigma=0.05,
↪
                                  learning_rates=0.1, discount_factors=dcf,
                                  Rmin=0, Rmax=1,
```

```
                                    method="weights", wU=w, use_prefactor=True)
            Visa_risk = aei.Visa(Xrisk)
            dX_risk = aei.step(Xrisk)[0] - Xrisk
            risky_stable_container[i, j, :] = dX_risk[0, :, 0] <= 0
            ag = 0; a = 1; s = 2
            Vis_risk = jnp.einsum(Visa_risk, [ag, s, a], Xrisk, [ag, s, a], [ag, s])
            Visa_safe = aei.Visa(Xsafe)
            dX_safe = aei.step(Xsafe)[0] - Xsafe
            safe_stable_container[i, j, :] = dX_safe[0, :, 0] >= 0
            Vis_safe = jnp.einsum(Visa_safe, [ag, s, a], Xsafe, [ag, s, a], [ag, s])
            risky_optimal_data_container[i, j, :] = Vis_risk[0,:] > Vis_safe[0,:]
            print("gamma = "+str(dcf)+", w = "+str(w)+" done")
    safe_stable_container = safe_stable_container * 2
    indicator_stable = safe_stable_container + risky_stable_container
    return risky_optimal_data_container, indicator_stable
```

```
 #| hide
import nbdev; nbdev.nbdev_export()
```

### Part IV

### Discretization calibration

In this section, we run error tests to determine which reward sensitivity $\sigma$ to choose in each environment. These results also give us insights into the effect of discretization on strategy. We run these tests on our three main environments: * two-armed bandit task; * risk-reward dilemma; * stag-hunt game;

First, we import what we need.

```
# general imports

import numpy as np

import jax.numpy as jnp

import matplotlib.pyplot as plt
import matplotlib as mpl

import pandas as pd
import seaborn as sns
```

```
# pyCRLD imports

# Agents
from pyCRLD.Agents.Base import abase
from pyCRLD.Agents.StrategySARSA import stratSARSA
from pyCRLD.Agents.StrategyActorCritic import stratAC

# Environments
from pyDDRL.Environments.SocialDilemma import SocialDilemma
from pyDDRL.Environments.RiskRewardDilemma import RiskRewardDilemma
from pyDDRL.Environments.TwoArmedBandit import TwoArmedBandit
```

```
# DDRL imports

from pyDDRL.Agents.DDRLBase import DDRLBase
from pyDDRL.Agents.DDRLSarsa import DDRLSarsa
from pyDDRL.Agents.DDRLActorCritic import DDRLActorCritic
```

#### Errors on a two-armed bandit task

Here, we compare a distributional and a non-distributional SARSA agent playing a "rich" two-armed bandit task ($p_1 = 0.9$, $p_2 = 0.7$) and a "poor" two-armed bandit task ($p_1 = 0.3$, $p_2 = 0.1$). The error at $X$ is defined as the norm of the difference between the temporal difference error of a distributional agent and that of a non-distributional agent for a given strategy $X$. We compute the error for different strategies $X$, and plot the mean error as a function of reward sensitivity $\sigma$. In **?@sec-results** , we will select the reward sensitivity that minimizes the error in order to compute our results.

First, we define a "poor" and a "rich" environment:

```
# Poor environment

xs = np.array([[-1, 1],
               [-1, 1]])
```

```
ps = np.array([[0.7, 0.3],
               [0.9, 0.1]])
env_poor = TwoArmedBandit(disttype='discrete', xs=xs, ps=ps)
```

```
# Rich environment

xs = np.array([[-1, 1],
               [-1, 1]])
ps = np.array([[0.1, 0.9],
               [0.3, 0.7]])
env_rich = TwoArmedBandit(disttype='discrete', xs=xs, ps=ps)
```

Now we define the reward sensitivities that we are going to compare, as well as the strategies for which we will compute the error:

```
# List of sigmas to compare
sigmas = np.linspace(0.0001, 1., 40)

# Sample strategies
Xs = np.linspace(0.01, 0.99, 11)
```

Here is the function that will allow us to compute the error:

```
# Function to compute mean error

def compute_mean_error(env, Xs, sigmas, path, version):
    errors = np.zeros((len(sigmas), len(Xs)))
    for j, sigma in enumerate(sigmas):  # loop over sigmas
        print("sigma = "+str(sigma))
        # non-distributional agent
        mae = stratSARSA(env=env, learning_rates=0.1,
                         discount_factors=0, choice_intensities=4.,
 ↪  use_prefactor=True)

        # distributional agent
        aei = DDRLSarsa(env, nr_reward_bins=29, discretization_sigma=sigma,
                        learning_rates=0.1, discount_factors=0,
                        choice_intensities=4, Rmin=-1, Rmax=1,
                        method="weights", wU=1, use_prefactor=True)

        for k, x in enumerate(Xs):  # loop over strategies
            # strategy X
            X = np.zeros((1, 1, 2))
            X[:, :, 0] = x
            X[:, :, 1] = 1 - x

            # TDe for distributional agent
            distRPE = aei.TDerror(X, norm=True)

            # TDe for non-distributional agent
```

```
            detRPE = mae.TDerror(X, norm=True)

            # compute the error
            error = np.linalg.norm(detRPE-distRPE)
            errors[j, k] = error

    # compute the mean error
    mean_error = np.mean(errors, axis=-1)

    # put results into df and save
    columns = ["X="+str(x) for x in Xs]
    index = ["sigma="+str(sigma) for sigma in sigmas]
    df = pd.DataFrame(errors, index=index, columns=columns)
    df.to_csv(path+"tab_error_df_"+version+".csv")
    return mean_error
```

```
# Path and version
path = "../data/error_calculation/"
version_poor = "poor_v1"
version_rich = "rich_v1"
```

```
# Compute mean error for poor and rich envs
mean_err_poor = compute_mean_error(env_poor, Xs, sigmas, path, version_poor)
mean_err_rich = compute_mean_error(env_rich, Xs, sigmas, path, version_rich)
```

```
# Extract the data
df_poor = pd.read_csv(path+"tab_error_df_"+version_poor+".csv")
df_rich = pd.read_csv(path+"tab_error_df_"+version_rich+".csv")

# Convert to np array
mean_err_poor = df_poor.to_numpy()[:, 1:]
mean_err_rich = df_rich.to_numpy()[:, 1:]
```

Now let’s plot the error:

```
# Plot
fig, (ax1, ax2) = plt.subplots(1, 2, figsize=(8, 3))
ax1.plot(sigmas, mean_err_poor)
ax2.plot(sigmas, mean_err_rich)
ax1.set_xlabel("discretization "+r'$\sigma$')
ax2.set_xlabel("discretization "+r'$\sigma$')
ax1.set_title("mean error in a poor environment")
ax2.set_title("mean error in a rich environment")
ax1.set_ylabel("mean error")
plt.show()
```

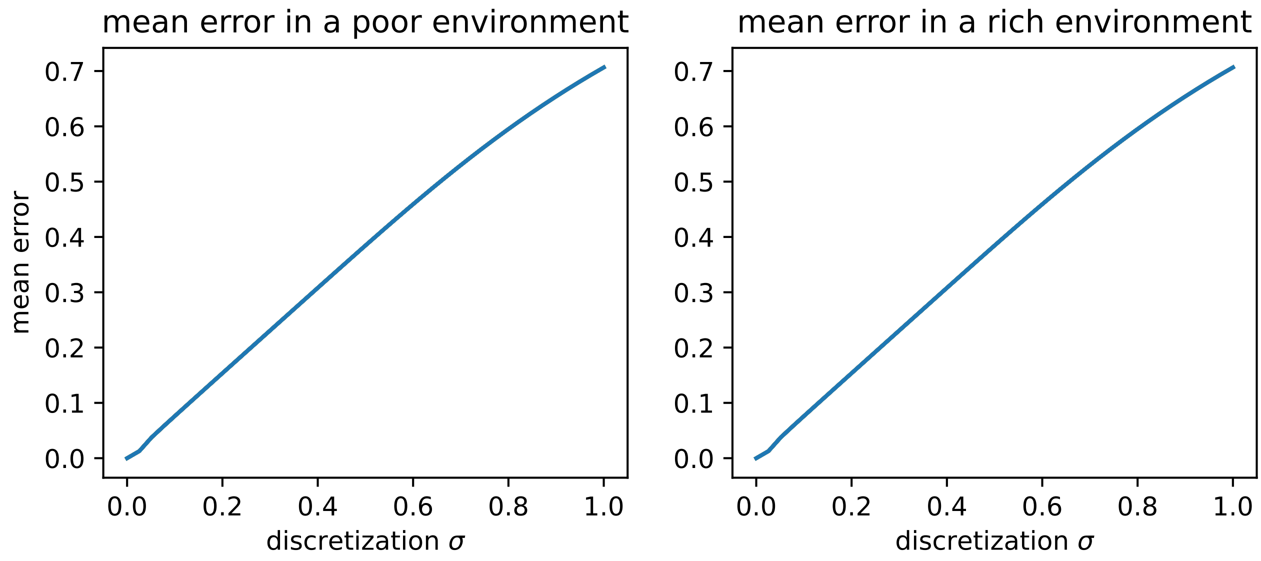

Figure 11.1: Mean error in a poor and in a rich environment as a function of reward sensitivity.

The reward sensitivity that minimizes the error is:

```
idx_rich, idx_poor = np.argmin(mean_err_rich), np.argmin(mean_err_poor)
sigmas[idx_rich], sigmas[idx_poor]
```

```
(np.float64(0.0001), np.float64(0.0001))
```

#### Errors in a stag-hunt game

Now we compare a distributional with a non-distributional agent playing the stag-hunt game defined Chapter 7 , with $R = 3$, $T = 1$, $P = 0$ and $S = -2$.

```
env = SocialDilemma(R=3, P=0, S=-2, T=1)
```

Once again, we compare different reward sensitivities $\sigma$ for the distributional agent. Since the game is symmetric in agents, we compute the error assuming that the second agent's strategy equals the first agent's strategy.

```
# List of sigmas to compare
sigmas = np.linspace(0.0001, 1., 40)

# Sample strategies
Xs = np.linspace(0.01, 0.99, 11)
```

```
# Function to compute mean error
def compute_mean_error(env, Xs, sigmas, gamma, path, version):
    errors = np.zeros((len(sigmas), len(Xs)))
    for j, sigma in enumerate(sigmas):  # loop over sigmas
        print("sigma = "+str(sigma))
```

```
        # Non-distributional agent
        mae = stratSARSA(env=env, learning_rates=0.1, choice_intensities=50,
                         discount_factors=gamma, use_prefactor=True)

        # Distributional agent
        aei = DDRLSarsa(env, nr_reward_bins=29, discretization_sigma=sigma,
                        learning_rates=0.1, discount_factors=gamma,
                        Rmin=-2, Rmax=3, choice_intensities=50,
                        method="weights", wU=1, use_prefactor=True)

        for k, x in enumerate(Xs):  # loop over strategies
            X = np.zeros((2, 1, 2))
            X[0, 0, 0] = x
            X[0, 0, 1] = 1 - x
            # Both agents have the same strategy
            X[1, 0, 0] = x
            X[1, 0, 1] = 1 - x

            # TDe for dist agent
            distRPE = aei.TDerror(X, norm=True)

            # TDe for non-dist agent
            detRPE = mae.TDerror(X, norm=True)

            # Compute the error
            error = np.linalg.norm(detRPE-distRPE)
            errors[j, k] = error

    # Compute mean error
    mean_error = np.mean(errors, axis=-1)

    # Format into df and save
    columns = ["X="+str(x) for x in Xs]
    index = ["sigma="+str(sigma) for sigma in sigmas]
    df = pd.DataFrame(errors, index=index, columns=columns)
    df.to_csv(path+"sh_error_df_"+version+".csv")
    return mean_error
```

```
# path and version
path = "../data/error_calculation/"
version = "v0"
```

We compute the error for a SARSA agent with discount factor $\gamma = 0.9$ and choice intensity $\beta = 50$, as in **?@sec-results** .

```
# Compute mean error
mean_err = compute_mean_error(env, Xs, sigmas, 0.9, path, version)
```

```
# Extract the data
df_error = pd.read_csv(path+"sh_error_df_"+version+".csv")
```

```
# Convert to np array
mean_err = np.mean(df_error.to_numpy()[:, 1:], axis=1)
```

Now we can plot the error as a function of $\sigma$:

```
# Plot
fig, ax1 = plt.subplots(1, 1, figsize=(4, 3))
ax1.plot(sigmas, mean_err)
ax1.set_xlabel("discretization "+r'$\sigma$')
ax1.set_title("mean error for "+r'$\gamma=0.9$')
ax1.set_ylabel("mean error")
plt.show()
```

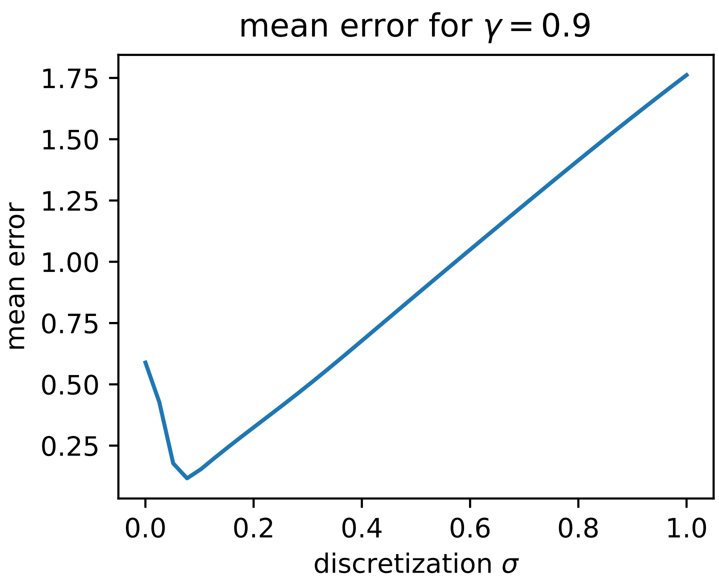

Figure 11.2: Mean error in a stag-hunt game as a function of reward sensitivity.

The reward sensitivity that minimizes the error is:

```
idx = np.argmin(mean_err)
sigmas[idx]
```

```
np.float64(0.07701538461538461)
```

#### Errors in a risk-reward dilemma

Here, we compute the error for a distributional Actor-Critic agent playing a risk-reward dilemma as defined in Chapter 8 , with $p_c = 0.2$, $p_r = 0.1$, $r_c = 0.5$, $r_r = 1$ and $r_d = 0$.

```
env = RiskRewardDilemma(pc=0.2, pr=0.1, rc=0.5, rr=1, rd=0)
```

Since the risk-reward dilemma contains 2 states, the discount factor $\gamma$ has a significant impact on the agent's strategy. Therefore, we will compute the error for an agent with a low discount factor ($\gamma = 0.4$), and for an agent with a high discount factor ($\gamma = 0.9$). In the degraded state, Actor-Critic agents converge to a maximally safe strategy since it is the only rational one (see **?@sec-results** ). Therefore, we compute the error for different strategies in the prosperous state, assuming the strategy in the degraded state is always $X^{safe} = 1$.

```
# Discount factors
gammas = [0.4, 0.9]

# List of sigmas to compare
sigmas = np.linspace(0.0001, 1., 40)

# Sample strategies
Xs = np.linspace(0.01, 0.99, 11)
```

```
# Function to compute mean error
def compute_mean_error(env, Xs, sigmas, gamma, path, version):
    errors = np.zeros((len(sigmas), len(Xs)))
    for j, sigma in enumerate(sigmas):  # loop over sigmas
        print("sigma = "+str(sigma))
        # Non-distributional agent
        mae = stratAC(env=env, learning_rates=0.1,
                      discount_factors=gamma, use_prefactor=True)

        # Distributional agent
        aei = DDRLActorCritic(env, nr_reward_bins=29, discretization_sigma=sigma,
                              learning_rates=0.1, discount_factors=gamma,
                              Rmin=0, Rmax=1,
                              method="weights", wU=1, use_prefactor=True)

        for k, x in enumerate(Xs):  # Loop over strategies
            X = np.zeros((1, 2, 2))
            X[:, 0, 0] = x
            X[:, 0, 1] = 1 - x
            X[:, 1, 0] = 1  # strategy in deg state

            # TDe for dist agent
            distRPE = aei.TDerror(X, norm=True)

            # TDe for non-dist agent
            detRPE = mae.TDerror(X, norm=True)

            # Compute error
            error = np.linalg.norm(detRPE-distRPE)
            errors[j, k] = error

    # Compute mean error
    mean_error = np.mean(errors, axis=-1)

    # Format into df and save
    columns = ["X="+str(x) for x in Xs]
    index = ["sigma="+str(sigma) for sigma in sigmas]
```

```
    df = pd.DataFrame(errors, index=index, columns=columns)
    df.to_csv(path+"rr_error_df_"+version+".csv")
    return mean_error
```

```
# Path and version
path = "../data/error_calculation/"
version_low = "low_v1"
version_high = "high_v1"
```

```
# Mean error for low gamma
mean_err_low = compute_mean_error(env, Xs, sigmas, gammas[0], path, version_low)

# Mean error for high gama
mean_err_high = compute_mean_error(env, Xs, sigmas, gammas[1], path, version_high)
```

```
# Extract the data
df_error_low = pd.read_csv(path+"rr_error_df_"+version_low+".csv")
df_error_high = pd.read_csv(path+"rr_error_df_"+version_high+".csv")

# Convert to np array
mean_err_low = np.mean(df_error_low.to_numpy()[:, 1:], axis=1)
mean_err_high = np.mean(df_error_high.to_numpy()[:, 1:], axis=1)
```

Now we can plot the error for both values of $\gamma$:

```
# Plot
fig, (ax1, ax2) = plt.subplots(1, 2, figsize=(8, 3))
ax1.plot(sigmas, mean_err_low)
ax2.plot(sigmas, mean_err_high)
ax1.set_xlabel("discretization "+r'$\sigma$')
ax2.set_xlabel("discretization "+r'$\sigma$')
ax1.set_title("mean error for "+r'$\gamma=0.4$')
ax2.set_title("mean error for "+r'$\gamma=0.9$')
ax1.set_ylabel("mean error")
plt.show()
```

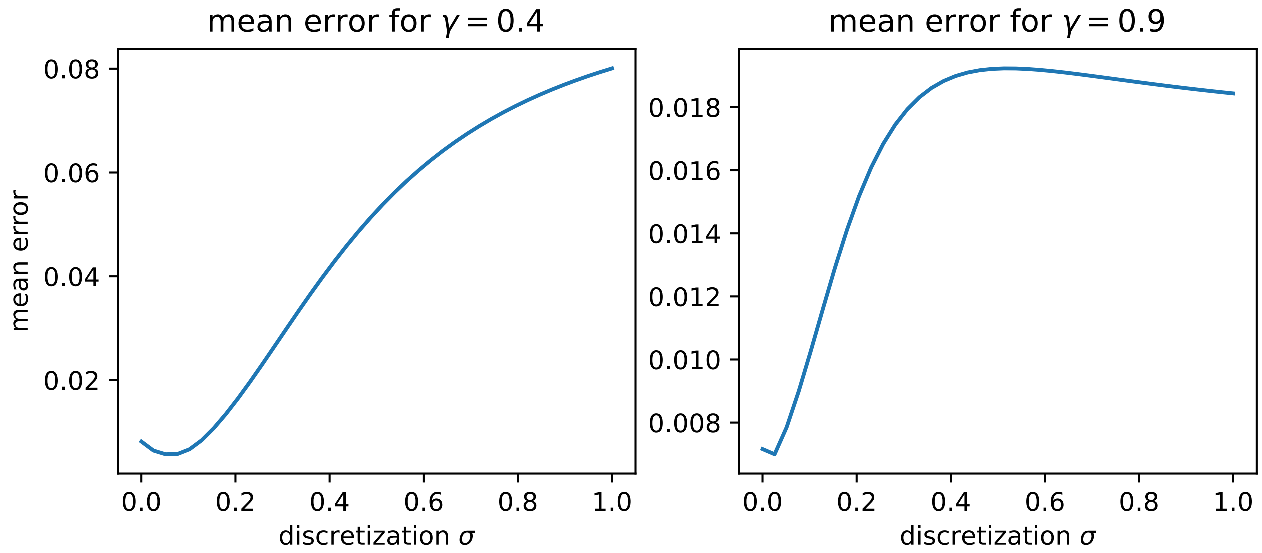

Figure 11.3: Mean error in a risk-reward dilemma as a function of reward sensitivity.

The reward sensitivity that minimizes the error for low discount factor is:

```
idx = np.argmin(mean_err_low)
sigmas[idx]
```

```
np.float64(0.05137692307692308)
```

And for high discount factor:

```
idx = np.argmin(mean_err_high)
sigmas[idx]
```

```
np.float64(0.025738461538461536)
```

### Part V

### Results

In this section, we show how we obtain our results. The figures shown here are the main and supplementary figures in our article. The key results can be summarized as follows:

- Chapter 12 : Exploration-Exploitation Challenge. The influence of optimism on performance in a two-armed bandit task is modulated by resource scarcity.
- Chapter 13 : Social Coordination Task. Optimism enhances cooperation in a stag-hunt game; optimism/pessimism has no effect on cooperation in a prisoner's dilemma.
- Chapter 14 : Intertemporal Risky Choice Problem. Optimism increases risk-seeking in a risk-reward dilemma.

### 12 Exploration-Exploitation Challenge

Cazé and Van der Meer (2013) showed that optimism/pessimism have an impact on exploration on a two-armed bandit task; importantly, this impact is modulated by resource scarcity (Cazé & Meer, 2013). Here, we see whether we can reproduce this effect. First, we implement optimism/pessimism with the three biasing methods defined in Chapter 5 . We find that the weights method closely approximates previous results on a two-armed bandit task. Then, we replicate Cazé and Van der Meer's findings, showing that optimism is beneficial in scarce environments, while pessimism is advantageous in the abundant version of the task.

First, we import what we need:

```
# general imports

import numpy as np
from scipy.stats import norm

import jax.numpy as jnp
from jax import jit
from functools import partial

import matplotlib.pyplot as plt
import matplotlib as mpl

import pandas as pd
import seaborn as sns
```

```
# pyCRLD imports

# Agents
from pyCRLD.Agents.Base import abase
from pyCRLD.Agents.StrategySARSA import stratSARSA

# Environment
from pyDDRL.Environments.TwoArmedBandit import TwoArmedBandit

# Utilities
from pyCRLD.Utils.Helpers import *
from pyDDRL.Utils.Trajectories1D import trajectories_1D
from pyDDRL.Utils.PhasePlot import phaseplot, _data_to_plot, _strategies, _dXisa_s
from pyDDRL.Utils.PlottingTools import *
```

```
# DDRL imports

from pyDDRL.Agents.DDRLBase import DDRLBase
from pyDDRL.Agents.DDRLSarsa import DDRLSarsa
```

#### 12.1 Comparison between the three DDRL biasing methods and asymmetric RL

Asymmetric RL was previously used to model optimism and pessimism in simple RL algorithms. It postulates a pair of asymmetric learning rates: a learning rate $\alpha^+$ that updates positive prediction errors, and a rate $\alpha^-$ that updates negative prediction errors. The bias strength $b$ is defined by the learning-rates ratio:

$$b = \frac{\alpha^+}{\alpha^-}$$

Cazé and Van der Meer (2013) (Cazé & Meer, 2013) have shown that, for an agent with bias strength $b$, the Q-value at steady-state of arm $a_i$ is:

$$Q_\infty^{a_i} = \frac{p_{a_i} \cdot b - (1 - p_{a_i})}{p_{a_i} \cdot b + (1 - p_{a_i})}$$

Below, we compare the final strategies obtained with the three DDRL biasing methods (quantile, expectile, weights) against the final strategies obtained with asymmetric RL.

Now, we create lists of bias strengths, quantiles, expectiles and weights.

The list of bias strengths is:

```
b_pes = np.linspace(0.1, 1., 10)  # pessimistic values
b_opt = np.linspace(1., 10., 10)  # optimistic values
b_ratios = np.zeros((19))
b_ratios[:9] = b_pes[:9]
b_ratios[9:] = b_opt
b_ratios
```

```
array([ 0.1,  0.2,  0.3,  0.4,  0.5,  0.6,  0.7,  0.8,  0.9,  1. ,  2. ,
        3. ,  4. ,  5. ,  6. ,  7. ,  8. ,  9. , 10. ])
```

Rowland *et al.* (2021) showed that, in stateless environments, expectile-based distributional RL is equivalent with asymmetric RL, when $\tau = \frac{b}{b+1}$ (Rowland et al., 2021).

We want to investigate whether this is the case on a two-armed bandit task, so we define the expectile list as:

```
e_list = b_ratios / (b_ratios + 1)
e_list
```

```
array([0.09090909, 0.16666667, 0.23076923, 0.28571429, 0.33333333,
       0.375     , 0.41176471, 0.44444444, 0.47368421, 0.5       ,
       0.66666667, 0.75      , 0.8       , 0.83333333, 0.85714286,
       0.875     , 0.88888889, 0.9       , 0.90909091])
```

Next, we define a quantile list with $\tau$ spanning the range of $[0, 1]$:

```
q_list = [0.05, 0.10, 0.15, 0.20, 0.25, 0.30,
          0.35, 0.40, 0.45, 0.50, 0.55, 0.60,
          0.65, 0.70, 0.75, 0.80, 0.85, 0.90,
          0.95]
```

Finally, we use a similar list for the upper weights $w_U$ as the one we used for the bias strengths $b$:

```
w_pes = np.linspace(0.1, 1., 10)
w_opt = np.linspace(1., 10., 10)
w_list = np.zeros((19))
w_list[:9] = w_pes[:9]
w_list[9:] = w_opt
w_list
```

```
array([ 0.1,  0.2,  0.3,  0.4,  0.5,  0.6,  0.7,  0.8,  0.9,  1. ,  2. ,
        3. ,  4. ,  5. ,  6. ,  7. ,  8. ,  9. , 10. ])
```

We can now compute and plot our comparisons on a poor two-armed bandit task. Here, we use a distributional SARSA agent with reward sensitivity $\sigma = 0.0001$, since we established in **?@sec-calibration** that this value of $\sigma$ minimizes the error on a two-armed bandit task.

```
# Compute comparisons in poor env
b_xinf, q_xinf, e_xinf, w_xinf = compute_comparisons(0.3, 0.1, b_ratios, q_list,
↪  e_list,
                                                     w_list, gamma=0, beta=4,
↪  Rmin=-1,
                                                     Rmax=1, nrb=29, discsig=0.0001,
↪  lr=0.1)
```

```
WARNING:2025-12-21 19:22:46,965:jax._src.xla_bridge:794: An NVIDIA GPU may be present on this 
enabled jaxlib is not installed. Falling back to cpu.
```

```
# Plot comparisons in poor env
plot_comparisons(b_ratios, q_list, e_list, w_list, b_xinf,
                 q_xinf, e_xinf, w_xinf, scarcity="poor",
                 save=True, savename='../figs/TwoArmedMethodComparisonPoor.png')

plt.show()
```

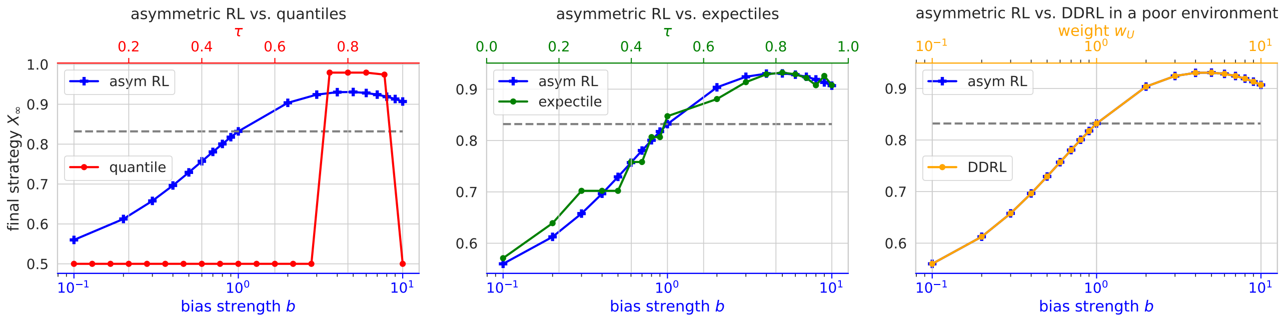

Figure 12.1: Final strategy of an agent playing a poor two-armed bandit task as a function of bias strength. The quantile, expectile, and weight methods are compared with asymmetric RL.

And now, in a rich environment:

```
# Compute comparisons in a rich environment
b_xinf, q_xinf, e_xinf, w_xinf = compute_comparisons(0.9, 0.7, b_ratios, q_list,
↪  e_list,
                                                   w_list, gamma=0, beta=4,
↪  Rmin=-1,
                                                   Rmax=1, nrb=29, discsig=0.0001,
↪  lr=0.1)
```

```
# Plot comparisons in a rich environment
plot_comparisons(b_ratios, q_list, e_list, w_list, b_xinf,
                 q_xinf, e_xinf, w_xinf, scarcity="rich",
                 save=True, savename='../figs/TwoArmedMethodComparisonRich.png')

plt.show()
```

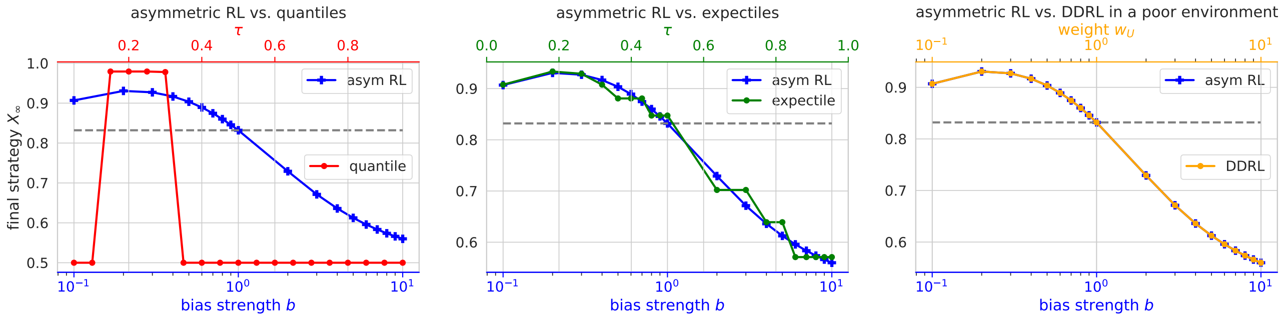

Figure 12.2: Final strategy of an agent playing a rich two-armed bandit task as a function of bias strength. The quantile, expectile, and weight methods are compared with asymmetric RL.

The figures show a close approximation of asymmetric RL by weights-DDRL. The equivalence between the two methods can be formally proven.

#### 12.2 Proof: Equivalence between weights-DDRL and asymmetric RL in a two-armed bandit task

We consider a multi-armed bandit with arms $a_i$, $i \in N$; for each $i$, $a_i$ yields a reward, $+1$, with probability $p_i$, and a punishment, $-1$, with probability $1 - p_i$.

An asymmetric RL agent is equipped with two different learning rates: $\alpha^+$ updates positive prediction errors, and $\alpha^-$ updates negative prediction errors. The agent's bias strength, $b$, is defined as the learning-rate ratio: $b = \frac{\alpha^+}{\alpha^-}$. Cazé and van der Meer (2013) have shown that, for such an agent, the Q-value at steady state for arm $a_i$ can be expressed as follows:

$$Q_\infty^{a_i} = \frac{bp_i - (1 - p_i)}{bp_i + (1 - p_i)} \tag{12.1}$$

Let us consider a weights-DDRL agent with number of neurons $n$, discretization sigma $\sigma$, discretization grid $x_j, j \in [1, n]$ with width $\Delta x$, and weight vector $w$ with elements $w_j$, $j \in [1, n]$. For each arm $a_i$, this agent computes a Q-value distribution $Q^{a_i,v}$, which quantifies the probability that the Q-value of arm $a_i$ equals a value $v$. Since the two-armed bandit is a stateless environment, the Q-value distribution $Q^{a_i,v}$ equals the reward distribution $R^{a_i,r}$, which quantifies the probability that arm $a_i$ yields a reward $r$.

Therefore, in the limit of an infinitely small discretization sigma $\sigma \to 0$, the Q-value distribution for arm $a_i$ can be described as follows:

- $Q^{a_i,v} = p_i$ when $v = 1$
- $Q^{a_i,v} = 1 - p_i$ when $v = -1$
- $Q^{a_i,v} = 0$ for all other values of $v$.

This is assuming that the agent is neutral – that is, all elements $w_j$ of vector $w$ have value 1. Now, let us consider an optimistic/pessimistic agent. For such an agent, $w_j = 1$ if $x_j < \bar{Q}^{a_i,v}$, and $w_j = w_U$ if $x_j \geq \bar{Q}^{a_i,v}$, where $\bar{Q}^{a_i,v}$ denotes the mean of the Q-value distribution.

The mean of the Q-value distribution is $\bar{Q}^{a_i,v} = 2p_i - 1$, hence $\bar{Q}^{a_i,v} \in [-1, 1]$. Excluding the extreme case in which there is an $i$, such that $p_i = 1$, we can formulate the weighted Q-value distribution $\tilde{Q}^{a_i,v}$:

- $\tilde{Q}^{a_i,v} = w_U p_i$ when $v = 1$
- $\tilde{Q}^{a_i,v} = 1 - p_i$ when $v = -1$
- $\tilde{Q}^{a_i,v} = 0$ for all other values of $v$.

We then normalize this distribution so that $\int_{x_1-\frac{\Delta x}{2}}^{x_n+\frac{\Delta x}{2}} \tilde{Q}^{a_i,v} dv = 1$. To do this, we divide the distribution by its integral, hence:

- $\tilde{Q}^{a_i,v} = \frac{w_U p_i}{w_U p_i + (1-p_i)}$ when $v = 1$
- $\tilde{Q}^{a_i,v} = \frac{1-p_i}{w_U p_i + (1-p_i)}$ when $v = -1$
- $\tilde{Q}^{a_i,v} = 0$ for all other values of $v$.

The mean of this new distribution is:

$$\overset{\simeq}{Q}^{a_i,v} = \frac{w_U p_i}{w_U p_i + (1-p_i)} - \frac{1-p_i}{w_U p_i + (1-p_i)} = \frac{w_U p_i - (1-p_i)}{w_U p_i + (1-p_i)}$$

Thus, on a multi-armed bandit task, an asymmetric-RL agent with bias strength $b$ learns the same Q-values at steady state as a weights-DDRL agent with upper weight value $w_U = b$.

#### 12.3 Influence of optimism/pessimism on performance

Here, we investigate how optimism/pessimism impact performance on a two-armed bandit task, when resources are scarce and when they are abundant.

##### 12.3.1 In a poor environment

In a “poor” two-armed bandit task, rewards are scarce. We thus define a two-armed bandit task with $p_1 = 0.3$, and $p_2 = 0.1$.

```
# Poor environment

xs = np.array([[-1, 1],
               [-1, 1]])
ps = np.array([[0.7, 0.3],
               [0.9, 0.1]])
env = TwoArmedBandit(disttype='discrete', xs=xs, ps=ps)
```

Now, we can plot the phase plot, strategy over time, and reward over time, of an agent which performs this task. We compare several levels of optimism and pessimism. These levels are defined by the upper weights $w_U$ - that is, the weights the agent applies to the upper part of the distribution.

```
# List of weight values to compare
weights = np.array([0.1, 0.2, 0.4, 0.8, 1., 2., 4., 8., 10.])

# i,s,a to plot along the x-axis
x = ([0], [0], [0])

# Range and resolution of X
res = np.linspace(0, 1., 21)
res[0] = 0.0001
res[-1] = 0.9999
```

In **?@sec-calibration** , we found that the discretization sigma yielding the lowest error for a neutral agent on a two-armed bandit task is $\sigma = 0.0001$. We therefore use it to compute our results.

```
compare_levels(env, weights, gamma=0., beta=4., Rmin=-1, Rmax=1,
               xinds=x, res=res, nrb=29, discsig=0.0001, traj_len=100,
               cmap='coolwarm', save=True,
               savename='../figs/TwoArmedPoorPhasePlot.png')

plt.show()
```

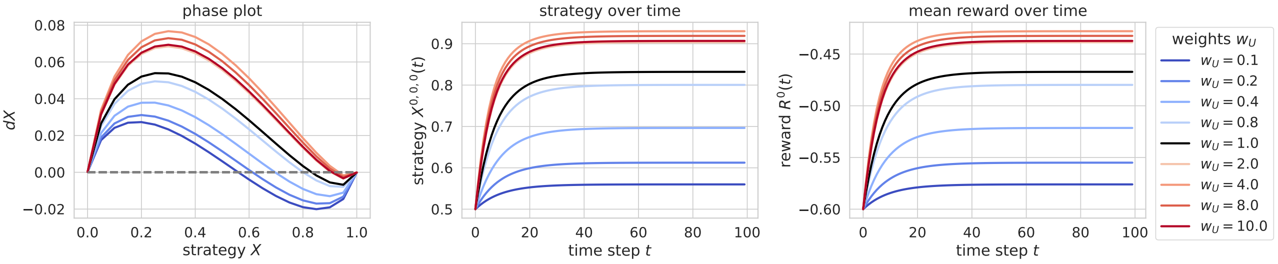

Figure 12.3: Phase plot, strategy plot and reward plot of a distributional agent playing a poor two-armed bandit task. Different levels of optimism/pessimism (quantified by the upper weight value $w_U$) are compared.

##### 12.3.2 In a rich environment

In a "rich" two-armed bandit task, rewards are abundant. We thus define a two-armed bandit task with $p_1 = 0.9$, and $p_2 = 0.7$.

```
# Rich environment

xs = np.array([[-1, 1],
               [-1, 1]])
ps = np.array([[0.1, 0.9],
               [0.3, 0.7]])
env = TwoArmedBandit(disttype='discrete', xs=xs, ps=ps)
```

Now, we can plot the phase plot, strategy over time, and reward over time, of an agent performing the rich task. We compare the same levels of optimism and pessimism as above.

```
compare_levels(env, weights, gamma=0., beta=4., Rmin=-1, Rmax=1,
               xinds=x, res=res, nrb=29, discsig=0.0001, traj_len=100,
               cmap='coolwarm', save=True,
               savename='../figs/TwoArmedRichPhasePlot.png')

plt.show()
```

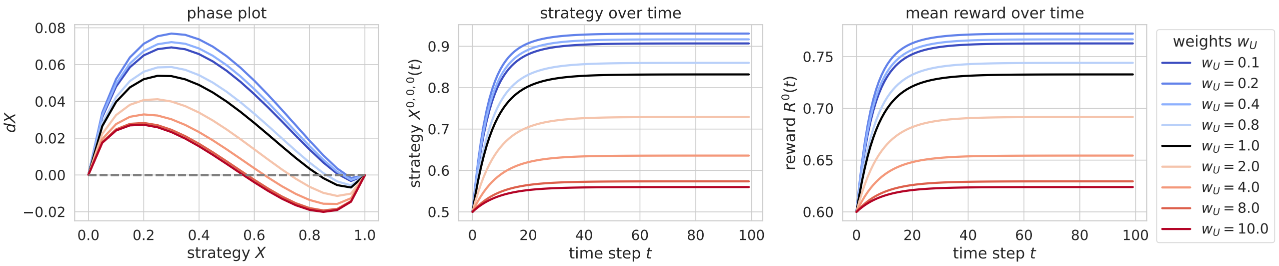

Figure 12.4: Phase plot, strategy plot and reward plot of a distributional agent playing a rich two-armed bandit task. Different levels of optimism/pessimism (quantified by the upper weight value $w_U$) are compared.

### 13 Social coordination task

In this section, we investigate the impact of optimism/pessimism in a social coordination task. To do this, we use the stag-hunt game defined in Chapter 7 . Additionally, we study the effect of optimism/pessimism on cooperation in a prisoner's dilemma.

In previous studies (such as (Matignon et al., 2007)), it has been suggested that optimism enhances cooperation on coordination tasks. This is what we replicate in this section.

First, we import what we need:

```
# general imports

import numpy as np
from scipy.stats import norm

import jax.numpy as jnp
from jax import jit
from functools import partial

import matplotlib.pyplot as plt
import matplotlib as mpl

import pandas as pd
import seaborn as sns
```

```
# pyCRLD imports

# Agents
from pyCRLD.Agents.Base import abase
from pyCRLD.Agents.StrategySARSA import stratSARSA

# Environment
from pyDDRL.Environments.SocialDilemma import SocialDilemma

# Utilities
from pyCRLD.Utils.Helpers import *
from pyDDRL.Utils.Trajectories1D import trajectories_1D
from pyDDRL.Utils.PhasePlot import phaseplot, _data_to_plot, _strategies, _dXisa_s
from pyDDRL.Utils.PlottingTools import *
```

```
# DDRL imports

from pyDDRL.Agents.DDRLBase import DDRLBase
from pyDDRL.Agents.DDRLSarsa import DDRLSarsa
```

Then, we define the environment:

```
env = SocialDilemma(R=3, P=0, S=-2, T=1)
```

#### 13.1 Sanity tests

We carry out sanity tests in order to check that the behavior of neutral DDRLSarsa agents imitate the behavior of regular SARSA agents. Once again, we compare the "mean", "expectile", and "weights" methods.

```
# Methods
methods = ["mean", "expectile", "weights"]

# Define agents to compare
aeis_ddrl = [] # DDRLSarsa agents
aeis_reg = []  # regular SARSA agents

for i, method in enumerate(methods):
    aei_ddrl = DDRLSarsa(env, nr_reward_bins=29, discretization_sigma=0.08,
                         learning_rates=0.1, discount_factors=0.9,
                         choice_intensities=50, Rmin=-2, Rmax=3,
                         method=method, use_prefactor=True)
    aeis_ddrl.append(aei_ddrl)

    aei_reg = stratSARSA(env=env, learning_rates=0.1, discount_factors=0.9,
                          choice_intensities=50, use_prefactor=True)
    aeis_reg.append(aei_reg)
```

```
WARNING:2025-12-21 19:38:18,828:jax._src.xla_bridge:794: An NVIDIA GPU may be present on this 
enabled jaxlib is not installed. Falling back to cpu.
```

Additionally, we plot trajectories with various initial conditions for better visualization.

```
# Phase space items to plot along the axes
x = ([0], [0], [0])  # agent 1
y = ([1], [0], [0])  # agent 2

# For trajectories
Xinits = []
xys = [(0.5, 0.5), (0.295, 0.205), (0.205, 0.695), (0.805, 0.305), (0.705, 0.795)]
cols = ['orange', 'purple', 'green', 'pink', 'brown']

for t, xy in enumerate(xys):
    Xinit = np.ones((2, 1, 2)) * 0.5
    Xinit[0, 0, 0] = xy[0]
    Xinit[1, 0, 0] = xy[1]
    Xinit[0, 0, 1] = 1 - Xinit[0, 0, 0]
    Xinit[1, 0, 1] = 1 - Xinit[1, 0, 0]
    Xinits.append(Xinit)

# Decorations
```

```
titles = ["method = mean", "method = expectile", "method = weights"]
xlabel = "prob. agent 1 cooperating"
ylabel = "prob. agent 2 cooperating"

# Figure and axes
fig, (ax1, ax2, ax3) = plt.subplots(1, 3, figsize=(3.2 * 3.8, 3.8))
axes = [ax1, ax2, ax3]

# Plot flow plots and trajectories
plot_3flowplots(aeis_reg, x, y, NrRandom=16, cmap='Reds', axes=axes)
plot_3flowplots(aeis_ddrl, x, y, NrRandom=16, cmap='Blues', decorate=True,
                titles=titles, xlabel=xlabel, ylabel=ylabel,
                Xinits=Xinits, cols=cols, axes=axes)

# Additional decorations
ax3.plot([], [], color='red', label='detSarsa')
ax3.plot([], [], color='blue', label='DDRLSarsa')
ax3.legend(bbox_to_anchor=(1.01, 1.01))

# Save the figure
plt.savefig('../figs/SHSanityTests.png', dpi=300, bbox_inches='tight')

plt.show()
```

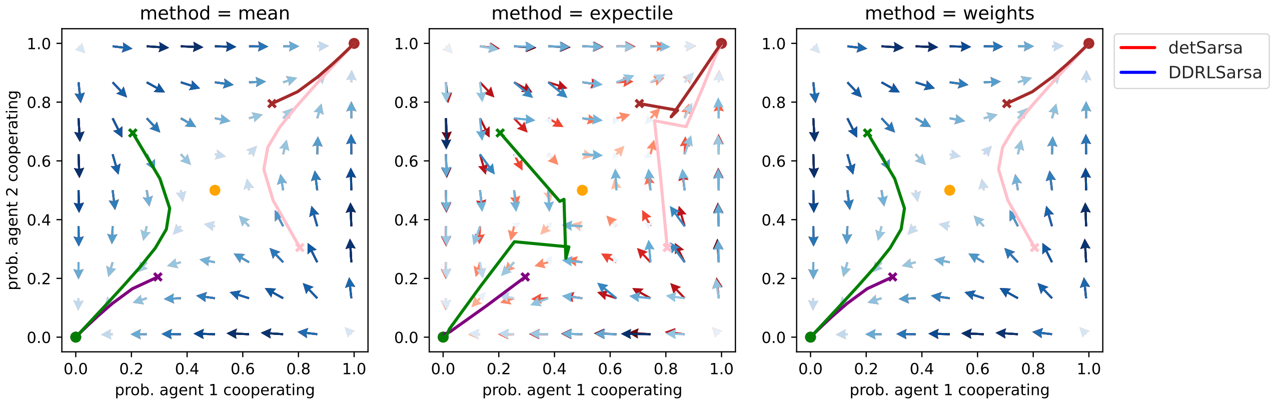

Figure 13.1: Comparison between a distributional and a non-distributional agent on a stag-hunt game. We compare distributional agents with a weights method ($w_U = 1$), with an expectile method ($ au=0.5$), and taking the mean of the Q-value distribution.

#### 13.2 Influence of optimism/pessimism

Once again, neutral weights-DDRLSarsa agents show less errors than neutral expectile-DDRLSarsa agents. Therefore, we use the weights method to explore the effects of optimism and pessimism on cooperation in the stag-hunt game. In particular, we look at how optimism/pessimism impacts the separatrix of the flowplot. This separatrix demarcates the domain where all initial strategies lead to mutual cooperation from the domain where all initial strategies lead to mutual defection.

We compare a pessimistic agent with $w_U = 0.25$, with a neutral agent ($w_U = 1$) and an optimistic agent ($w_U = 4$).

```
weights = [0.25, 1., 4.]

# Define agents to compare
aeis = [] # DDRLSarsa agents

for i, weight in enumerate(weights):
    aei = DDRLSarsa(env, nr_reward_bins=29, discretization_sigma=0.08,
                    learning_rates=0.1, discount_factors=0.9,
                    choice_intensities=50, Rmin=-2, Rmax=3,
                    method="weights", wU=weight, use_prefactor=True)
    aeis.append(aei)
```

Now plotting:

```
# Decorations
titles = [r'$w_U=0.25$', r'$w_U=1$', r'$w_U=4$']
xlabel = "prob. agent 1 cooperating"
ylabel = "prob. agent 2 cooperating"

# Figure and axes
fig, (ax1, ax2, ax3) = plt.subplots(1, 3, figsize=(3.2 * 3.8, 3.8))
axes = [ax1, ax2, ax3]

# Plot flow plots and trajectories
plot_3flowplots(aeis, x, y, NrRandom=16, cmap='Blues', decorate=True,
                titles=titles, xlabel=xlabel, ylabel=ylabel,
                Xinits=Xinits, cols=cols, axes=axes)

# Plot separatrices
for i, aei in enumerate(aeis):
    ax = axes[i]
    plot_separatrix(aei, ax, x, y)

# Save the figure
plt.savefig('../figs/SHFlowPlots.png', dpi=300, bbox_inches='tight')

plt.show()
```

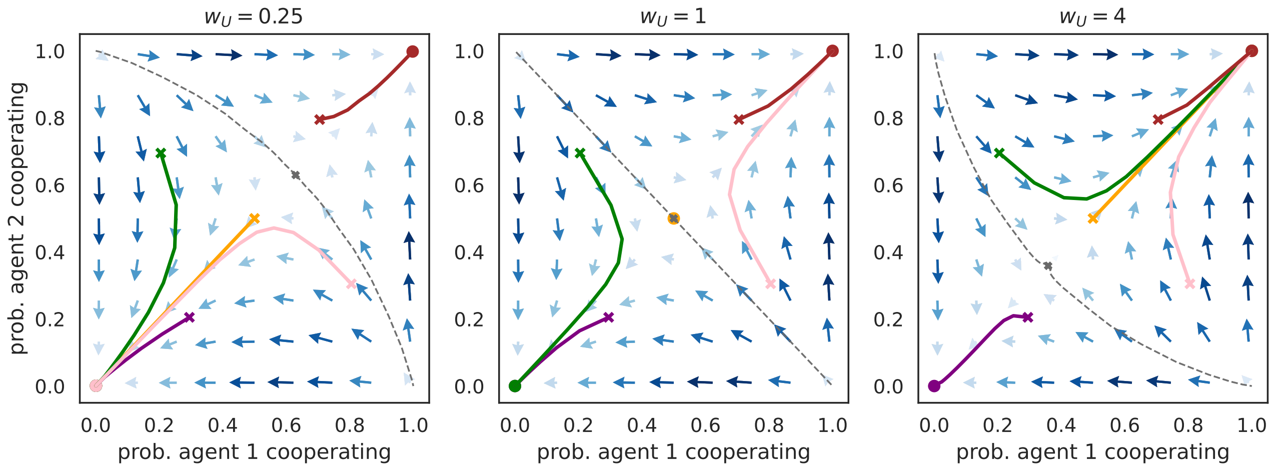

Figure 13.2: Flow plots on a stag-hunt game for a pessimistic, neutral and optimistic agent.

We can also plot the phase plot for one agent, assuming the other agent always plays the same strategy as the first one. This is equivalent with visualizing the diagonal of the above flowplots.

```
weights = np.array([0.1, 0.2, 0.4, 0.8, 1., 2., 4., 8., 10.])

x = ([0], [0], [0])
res = np.linspace(0, 1., 21)
res[0] = 0.0001
res[-1] = 0.9999
otherXs = np.linspace(0, 1., 21)
otherXs[0] = 0.0001
otherXs[-1] = 0.9999

compare_levels(env, weights, gamma=0.9, beta=50., Rmin=-2, Rmax=3,
               xinds=x, res=res, nrb=29, discsig=0.08, traj_len=30,
               otherXs=otherXs, cmap='coolwarm', save=True,
               savename='../figs/SHPhasePlots.png')

plt.show()
```

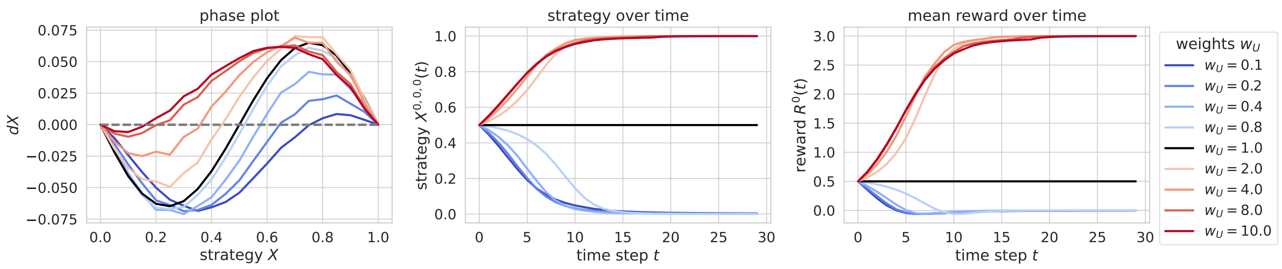

Figure 13.3: Phase plot, strategy plot and reward plot of an agent playing a stag-hunt game. Different levels of optimism/pessimism are compared.

#### 13.3 Optimism has no effect in a prisoner's dilemma

As a complement, we can turn the stag-hunt game into a prisoner's dilemma, which is another social dilemma with a different reward structure. In a prisoner's dilemma, typically, mutual cooperation is not an equilibrium; both agents are incentivized to defect. First, we define the environment:

```
env = SocialDilemma(R=1, P=0, S=-2, T=3)
```

And we carry the same sanity tests as above:

```
methods = ["mean", "expectile", "weights"]

# Define agents to compare
aeis_ddrl = [] # DDRLSarsa agents
aeis_reg = []  # regular SARSA agents

for i, method in enumerate(methods):
    aei_ddrl = DDRLSarsa(env, nr_reward_bins=29, discretization_sigma=0.08,
                         learning_rates=0.1, discount_factors=0.9,
                         choice_intensities=50, Rmin=-2, Rmax=3,
                         method=method, use_prefactor=True)
    aeis_ddrl.append(aei_ddrl)

    aei_reg = stratSARSA(env=env, learning_rates=0.1, discount_factors=0.9,
                         choice_intensities=50, use_prefactor=True)
    aeis_reg.append(aei_reg)
```

```
# Phase space items to plot on axes
x = ([0], [0], [0])  # agent 1
y = ([1], [0], [0])  # agent 2

# For trajectories
Xinits = []
xys = [(0.5, 0.5), (0.295, 0.205), (0.205, 0.695), (0.805, 0.305), (0.705, 0.795)]
cols = ['orange', 'purple', 'green', 'pink', 'brown']

for t, xy in enumerate(xys):
    Xinit = np.ones((2, 1, 2)) * 0.5
    Xinit[0, 0, 0] = xy[0]
    Xinit[1, 0, 0] = xy[1]
    Xinit[0, 0, 1] = 1 - Xinit[0, 0, 0]
    Xinit[1, 0, 1] = 1 - Xinit[1, 0, 0]
    Xinits.append(Xinit)

# Decorations
titles = ["method = mean", "method = expectile", "method = weights"]
xlabel = "prob. agent 1 cooperating"
ylabel = "prob. agent 2 cooperating"

# Figure and axes
fig, (ax1, ax2, ax3) = plt.subplots(1, 3, figsize=(3.2 * 3.8, 3.8))
```

```
axes = [ax1, ax2, ax3]

# Plot flow plots and trajectories
plot_3flowplots(aeis_reg, x, y, NrRandom=16, cmap='Reds', axes=axes)
plot_3flowplots(aeis_ddrl, x, y, NrRandom=16, cmap='Blues', decorate=True,
                titles=titles, xlabel=xlabel, ylabel=ylabel,
                Xinits=Xinits, cols=cols, axes=axes)

# Additional decorations
ax3.plot([], [], color='red', label='detSarsa')
ax3.plot([], [], color='blue', label='DDRLSarsa')
ax3.legend(bbox_to_anchor=(1.01, 1.01))

# Save the figure
plt.savefig('../figs/PDSanityTests.png', dpi=300, bbox_inches='tight')

plt.show()
```

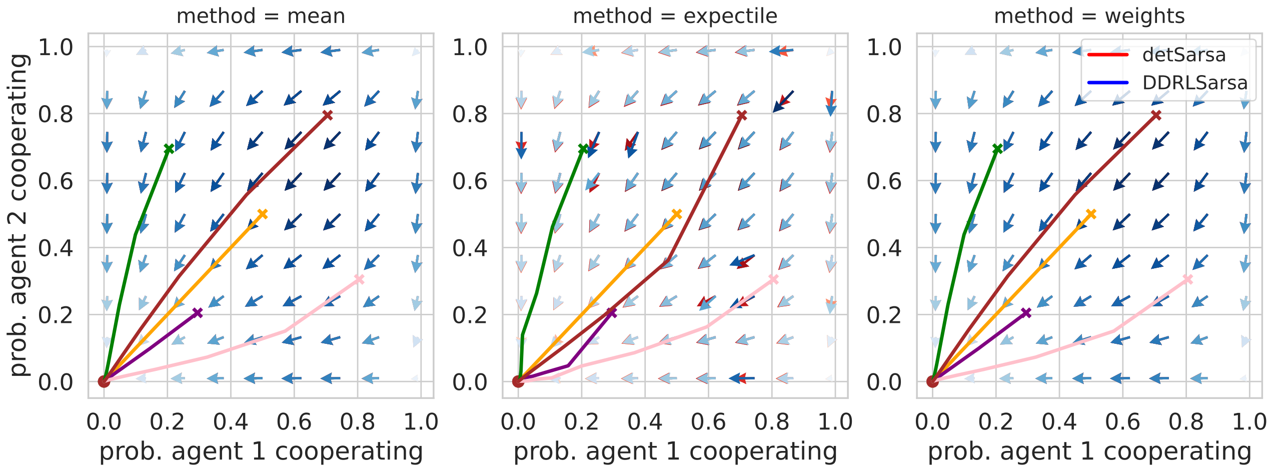

Figure 13.4: Comparison between a distributional and a non-distributional agent on a prisoner's dilemma. We compare distributional agents with a weights method, with an expectile method, and taking the mean of the Q-value distribution.

We can investigate the influence of optimism or pessimism by plotting the same phase plots and strategy plots as above, assuming the second agent always behaves like the first.

```
weights = np.array([0.1, 0.2, 0.4, 0.8, 1., 2., 4., 8., 10.])

x = ([0], [0], [0])
res = np.linspace(0, 1., 21)
res[0] = 0.0001
res[-1] = 0.9999
otherXs = np.linspace(0, 1., 21)
otherXs[0] = 0.0001
otherXs[-1] = 0.9999
```

We can prove that optimism has an effect on cooperation in a stag-hunt game, but not in a prisoner's dilemma.

#### 13.4 Proof: Effect of optimism on social dilemmas

We consider a social dilemma, in which two agents can cooperate ($c$) or defect ($d$).

Mutual cooperation yields a "reward" $R$ for both agents, while mutual defection yields a "punishment" $P$. If one agent cooperates and the other defects, the cooperator gets a "sucker" payoff $S$, while the defector gets a "temptation" payoff $T$. The stag-hunt game and the prisoner's dilemma differ in their reward structure:

- In a stag-hunt game, $R > T > P > S$;
- In a prisoner's dilemma, $T > R > P > S$.

We want to find the conditions under which the Q-value of cooperation for agent $i$, $Q^{i,c}$, is greater than the Q-value for defection for agent $i$, $Q^{i,d}$. To do this, we consider a weights-DDRL agent with number of neurons $n$, discretization sigma $\sigma$, discretization grid $x_j, j \in [1, n]$ with width $\Delta x$, and weight vector $w$ with elements $w_j$, $j \in [1, n]$. Moreover, we place ourselves in the limit of an infinitely small discretization sigma, $\sigma \to 0$, and we assume the agent's discount factor $\gamma$ is 0. As a consequence, the Q-value distributions $Q^{i,c,v}$ and $Q^{i,d,v}$ just depend on the payoff and on the other agent $j$'s strategy $X^{j,c}$.

- $Q^{i,c,v} = R$ with probability $X^{j,c}$;
- $Q^{i,c,v} = S$ with probability $1 - X^{j,c}$;
- $Q^{i,d,v} = P$ with probability $X^{j,d} = 1 - X^{j,c}$;
- $Q^{i,d,v} = T$ with probability $1 - X^{j,d} = X^{j,c}$.

The mean Q-value $\bar{Q}^{i,c,v}$ is therefore $X^{j,c} \cdot R + (1 - X^{j,c}) \cdot S = S + X^{j,c}(R - S)$. We consider $X^{j,c} \in ]0, 1[$. In both social dilemmas, $R > S$, from which it follows that $S < \bar{Q}^{i,c,v} < R$. As a consequence, the biased and normalized distribution $\tilde{Q}^{i,c,v}$ is described by:

- $Q^{i,c,v} = R$ with probability $\frac{w_U \cdot X^{j,c}}{w_U \cdot X^{j,c} + (1 - X^{j,c})}$;
- $Q^{i,c,v} = S$ with probability $\frac{1 - X^{j,c}}{w_U \cdot X^{j,c} + (1 - X^{j,c})}$.

A similar reasoning can be applied to the Q-value distribution $Q^{i,d,v}$, and we obtain:

- $Q^{i,d,v} = P$ with probability $\frac{1 - X^{j,c}}{w_U \cdot X^{j,c} + (1 - X^{j,c})}$;
- $Q^{i,d,v} = T$ with probability $\frac{w_U \cdot X^{j,c}}{w_U \cdot X^{j,c} + (1 - X^{j,c})}$.

Consequently, the means of these biased distributions are:

$$\overset{\simeq}{Q}^{i,c,v} = \frac{w_U \cdot X^{j,c} \cdot R + (1 - X^{j,c}) \cdot S}{w_U \cdot X^{j,c} + (1 - X^{j,c})} = \frac{X^{j,c} \cdot (w_U \cdot R - S) + S}{X^{j,c} \cdot (w_U - 1) + 1} \tag{13.1}$$

$$\overset{\simeq}{Q}^{i,d,v} = \frac{w_U \cdot X^{j,c} \cdot T + (1 - X^{j,c}) \cdot P}{w_U \cdot X^{j,c} + (1 - X^{j,c})} = \frac{X^{j,c} \cdot (w_U \cdot T - P) + P}{X^{j,c} \cdot (w_U - 1) + 1} \tag{13.2}$$

We have $\overset{\simeq}{Q}^{i,c,v} \geq \overset{\simeq}{Q}^{i,c,d}$

iff $X^{j,c} \cdot (w_U \cdot R - S) + S \geq X^{j,c} \cdot (w_U \cdot T - P) + P$

iff $X^{j,c} \cdot [w_U(R - T) + (P - S)] + (S - P) \geq 0$

iff $X^{j,c} \cdot [w_U(R - T)] + (S - P)[1 - X^{j,c}] \geq 0$.

In both social dilemmas, $P > S$ so $S - P < 0$. Therefore, $(S - P)[1 - X^{j,c}] < 0$.

In a prisoner's dilemma, $T > R$, so $R - T < 0$, and as a consequence, $X^{j,c} \cdot [w_U(R-T)] < 0$. Therefore, in a prisoner's dilemma, it is not possible to have $\overset{\simeq}{Q}^{i,c,v} \geq \overset{\simeq}{Q}^{i,c,d}$. In other terms, the agents always mutually defect.

In a stag-hunt game, however, $R > T$, so $R - T > 0$, and as a consequence, $X^{j,c} \cdot [w_U(R-T)] > 0$. So we have $\overset{\simeq}{Q}^{i,c,v} \geq \overset{\simeq}{Q}^{i,c,d}$

iff $w_U(R-T) \geq \frac{(P-S)(1-X^{j,c})}{X^{j,c}}$

iff $w_U \geq \frac{(P-S)(1-X^{j,c})}{(R-T)X^{j,c}}$.

Therefore, in a stag-hunt game, optimism/pessimism has an effect on cooperation.

We can plot $w_U$ as a function of $X^{j,c}$ for the stag-hunt game we investigate in our paper, that is: $R = 3$, $T = 1$, $P = 0$, and $S = -2$.

```
compare_levels(env, weights, gamma=0.9, beta=50., Rmin=-2, Rmax=3,
               xinds=x, res=res, nrb=29, discsig=0.08, traj_len=30,
               otherXs=otherXs, cmap='coolwarm', save=False,
               savename='figs/PDPhasePlots.png')

plt.show()
```

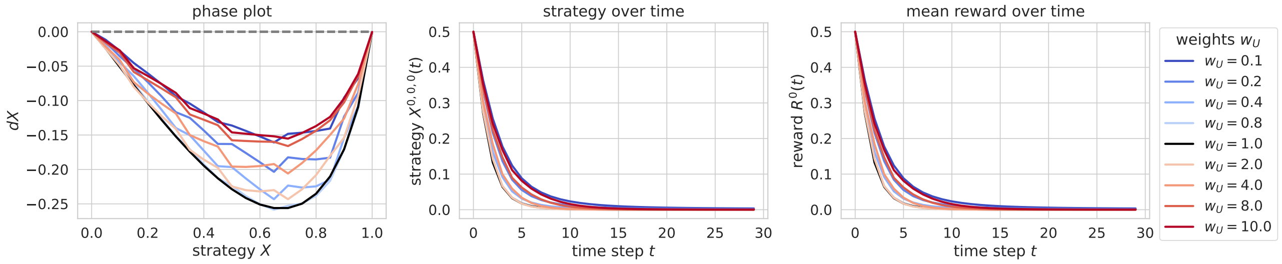

Figure 13.5: Phase plot, strategy plot and reward plot of an agent playing a prisoner's dilemma. Different levels of optimism/pessimism are compared.

```
def w_coop(X, R=3, T=1, P=0, S=-2):
    return (P-S) * (1-X) / ((R-T) * X)
```

```
X = np.linspace(0.05, 1, 20)
ws = w_coop(X)
ones = np.ones((len(X)))

plt.plot(X, ws, 'o-')
plt.plot(X, ones, '--', color="gray", label=r'$w_U = 1$')
plt.xlabel("probability of agent 2 cooperating")
plt.ylabel("minimum "+r'$w_U$'+" for agent 1 to cooperate")
plt.legend()

plt.show()
```

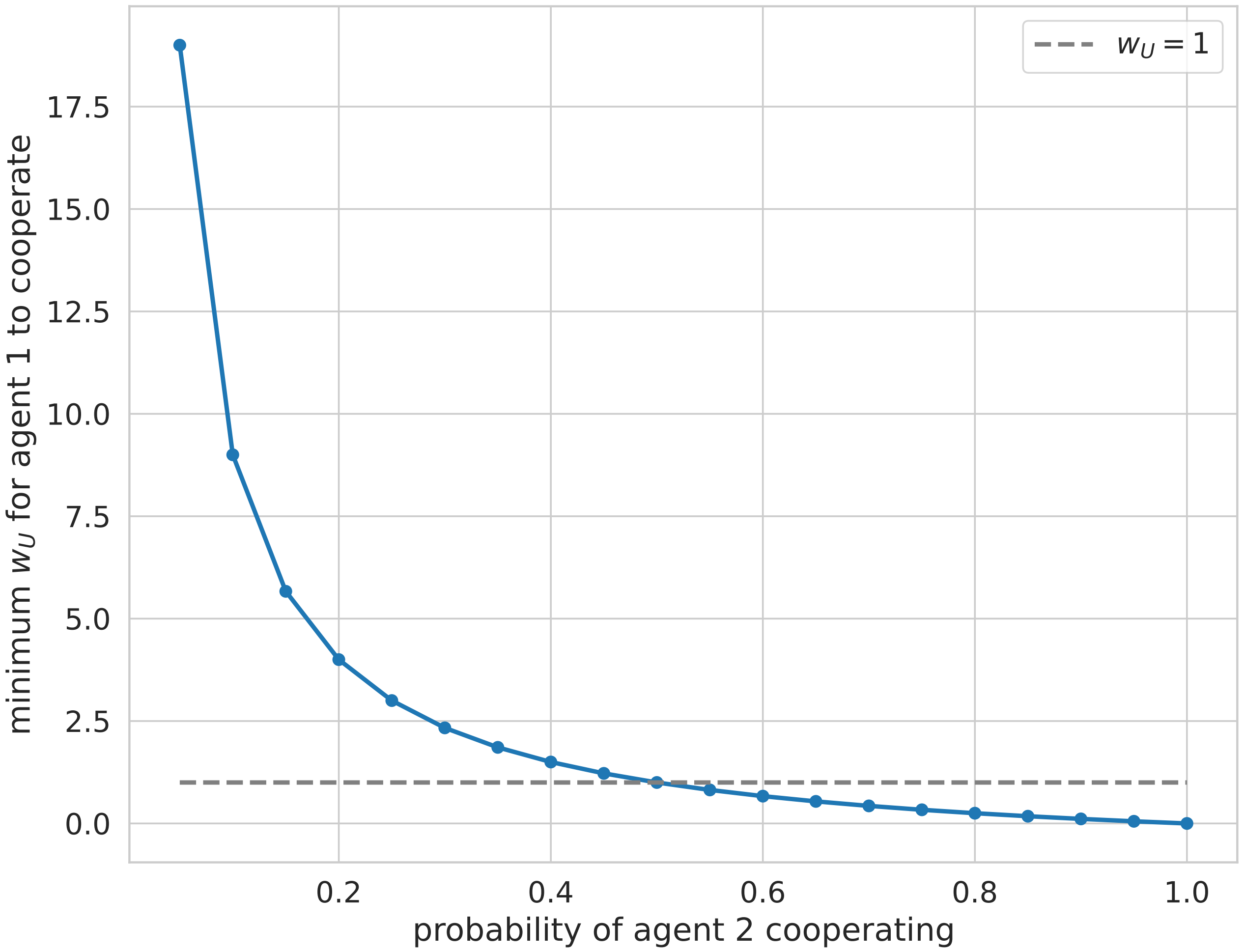

$w_U = 1$
minimum $w_U$ for agent 1 to cooperate
probability of agent 2 cooperating
0.0
2.5
5.0
7.5
10.0
12.5
15.0
17.5
0.2
0.4
0.6
0.8
1.0

### 14 Intertemporal risky choice problem

In this section, we study a risk-reward dilemma in which an agent can choose between a high-risk, high-gain option, and a low-risk, low-gain option (see Section Chapter 8 ).

Previous studies (such as (Mihatsch & Neuneier, 2002) and (Shen et al., 2014)) have framed optimism/pessimism in terms of risk-sensitivity. Therefore, we investigate whether optimism increases risk-seeking in this intertemporal risky choice problem. We find that, for different values of the discount factor $\gamma$, optimism does increase risk-seeking in this environment. Furthermore, we uncover the existence of bistable regimes and "incoherent choices" for certain values of $\gamma$ and $w_U$.

First, we import what we need:

```
# general imports

import numpy as np
from scipy.stats import norm

import jax.numpy as jnp
from jax import jit
from functools import partial

import matplotlib.pyplot as plt
import matplotlib as mpl

import pandas as pd
import seaborn as sns
```

```
# pyCRLD imports

# Agents
from pyCRLD.Agents.Base import abase
from pyCRLD.Agents.StrategyActorCritic import stratAC

# Environment
from pyDDRL.Environments.RiskRewardDilemma import RiskRewardDilemma

# Utilities
from pyCRLD.Utils.Helpers import *
from pyCRLD.Utils import FlowPlot as fp
from pyDDRL.Utils.Trajectories1D import trajectories_1D
from pyDDRL.Utils.PhasePlot import phaseplot, _data_to_plot, _strategies, _dXisa_s
from pyDDRL.Utils.PlottingTools import *
```

```
# DDRL imports

from pyDDRL.Agents.DDRLBase import DDRLBase
from pyDDRL.Agents.DDRLActorCritic import DDRLActorCritic
```

Then, we define the environment:

```
# Define environment
env = RiskRewardDilemma(pc=0.2, pr=0.1, rc=0.5, rr=1, rd=0)
```

In this section, we use Actor-Critic agents (defined in Section Chapter 4 ). The reason we use such agents is that, unlike SARSA agents, they have no exploratory parameter (such as the inverse temperature $\beta$). Therefore, we can investigate the effects of optimism and pessimism on the Actor-Critic agents' trajectories towards a safe or risky strategy, without considering the effects of the heuristics on exploration in the degraded state.

#### 14.1 Sanity tests

First, we carry out sanity tests in order to check that DDRLActorCritic agents yield the same flow plots as a regular AC agent. We compare different DDRL biasing methods in order to choose the most appropriate. To this end, we compare DDRLActorCritic agents and regular ActorCritic agents across 3 values of the discount factor $\gamma$: $\gamma = 0$, $\gamma = 0.4$, and $\gamma = 0.9$. First, we create neutral DDRLActorCritic agents which take the mean of their value distribution as their value estimate.

```
# Values of gamma
gammas = [0., 0.4, 0.9]
```

In Section **?@sec-calibration** , we found that the discretization sigma that yields the minimal error is $\sigma = 0.05$ for $\gamma = 0.4$, and $\sigma = 0.03$ for $\gamma = 0.9$. For these sanity tests, we use $\sigma = 0.05$, because it does not increase the error when $\gamma = 0.9$ by much.

```
# Define agents to compare
aeis_reg = []  # regular SARSA agents
aeis_ddrl = []  # DDRLSarsa agents

for i, gamma in enumerate(gammas):
    aei_reg = stratAC(env=env, learning_rates=0.1, discount_factors=gamma,
                      use_prefactor=True)
    aeis_reg.append(aei_reg)

    aei_ddrl = DDRLActorCritic(env, nr_reward_bins=29, discretization_sigma=0.05,
                               learning_rates=0.1, discount_factors=gamma,
                               Rmin=0, Rmax=1,
                               method="mean", use_prefactor=True)
    aeis_ddrl.append(aei_ddrl)
```

```
WARNING:2025-12-21 19:48:20,771:jax._src.xla_bridge:794: An NVIDIA GPU may be present on this 
enabled jaxlib is not installed. Falling back to cpu.
```

Now, we can visualize the flowplots:

```
# Phase space items to plot along the x- and y- axes
x = ([0], [0], [0])  # prosperous state
y = ([0], [1], [0])  # degraded state

# For trajectories
Xinits = [np.ones((1, 2, 2)) * 0.5]  # initial strategy
cols = ["orange"]  # trajectory color

# Decorations
titles = [r'$\gamma = 0.0$', r'$\gamma = 0.4$', r'$\gamma = 0.9$']
xlabel = "prob. safe action in prosp. state"
ylabel = "prob. safe action in deg. state"

# Figure and axes
fig, (ax1, ax2, ax3) = plt.subplots(1, 3, figsize=(3.2 * 3.8, 3.8))
axes = [ax1, ax2, ax3]

# Plot flow plots and trajectories
plot_3flowplots(aeis_reg, x, y, NrRandom=16, cmap='Reds', axes=axes)
plot_3flowplots(aeis_ddrl, x, y, NrRandom=16, cmap='Blues', decorate=True,
                titles=titles, xlabel=xlabel, ylabel=ylabel,
                Xinits=Xinits, cols=cols, axes=axes)

# Additional decorations
ax3.plot([], [], color='red', label='detAC')
ax3.plot([], [], color='blue', label='DDRL AC')
ax3.legend(bbox_to_anchor=(1.01, 1.01))

# Save the figure
plt.savefig('../figs/RRSanityTestMean.png', dpi=300, bbox_inches='tight')

plt.show()
```

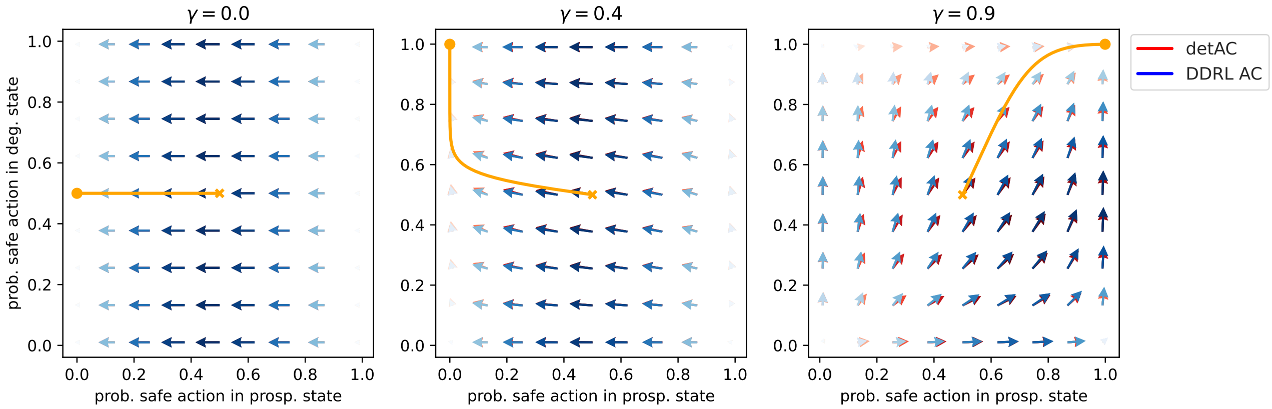

Figure 14.1: Comparison between a distributional and a non-distributional agent in a risk-reward dilemma, for several values of the discount factor. For the distributional agent, the mean of the Q-value distribution was taken.

Now, we compare the regular AC agents with an expectile-DDRLActorCritic agent ($\tau = 0.5$, which should be equivalent with taking the mean of the Q-value distribution). Defining the expectile agents:

```
# Define agents to compare
aeis_ddrl = []  # DDRLSarsa agents

for i, gamma in enumerate(gammas):
    aei_ddrl = DDRLActorCritic(env, nr_reward_bins=29, discretization_sigma=0.05,
                               learning_rates=0.1, discount_factors=gamma,
                               Rmin=0, Rmax=1,
                               method="expectile", tau=0.5, use_prefactor=True)
    aeis_ddrl.append(aei_ddrl)
```

And visualizing the flowplots:

```
# Figure and axes
fig, (ax1, ax2, ax3) = plt.subplots(1, 3, figsize=(3.2 * 3.8, 3.8))
axes = [ax1, ax2, ax3]

# Plot flow plots and trajectories
plot_3flowplots(aeis_reg, x, y, NrRandom=16, cmap='Reds', axes=axes)
plot_3flowplots(aeis_ddrl, x, y, NrRandom=16, cmap='Blues', decorate=True,
                titles=titles, xlabel=xlabel, ylabel=ylabel,
                Xinits=Xinits, cols=cols, axes=axes)

# Additional decorations
ax3.plot([], [], color='red', label='detAC')
ax3.plot([], [], color='blue', label='DDRL AC')
ax3.legend(bbox_to_anchor=(1.01, 1.01))

# Save the figure
plt.savefig('../figs/RRSanityTestExpectile.png', dpi=300, bbox_inches='tight')

plt.show()
```

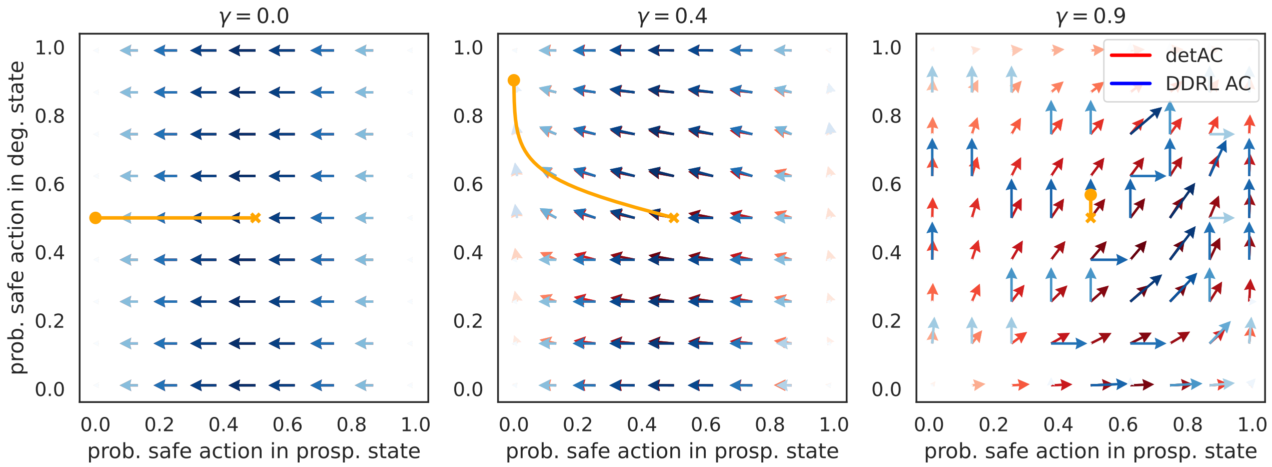

Figure 14.2: Comparison between a distributional and a non-distributional agent in a risk-reward dilemma, for several values of the discount factor. For the distributional agent, the expectile method was used.

Finally, we compare the regular AC agents with a weights-ActorCritic agent ($w_U = 1$, which should be equivalent with taking the mean of the Q-value distribution). Defining the weights agents:

```
# Define agents to compare
aeis_ddrl = []  # DDRLSarsa agents

for i, gamma in enumerate(gammas):
    aei_ddrl = DDRLActorCritic(env, nr_reward_bins=29, discretization_sigma=0.05,
                               learning_rates=0.1, discount_factors=gamma,
                               Rmin=0, Rmax=1,
                               method="weights", wU=1., use_prefactor=True)
    aeis_ddrl.append(aei_ddrl)
```

Now visualizing:

```
# Figure and axes
fig, (ax1, ax2, ax3) = plt.subplots(1, 3, figsize=(3.2 * 3.8, 3.8))
axes = [ax1, ax2, ax3]

# Plot flow plots and trajectories
plot_3flowplots(aeis_reg, x, y, NrRandom=16, cmap='Reds', axes=axes)
plot_3flowplots(aeis_ddrl, x, y, NrRandom=16, cmap='Blues', decorate=True,
                titles=titles, xlabel=xlabel, ylabel=ylabel,
                Xinits=Xinits, cols=cols, axes=axes)

# Additional decorations
ax3.plot([], [], color='red', label='detAC')
ax3.plot([], [], color='blue', label='DDRL AC')
ax3.legend(bbox_to_anchor=(1.01, 1.01))

# Save the figure
plt.savefig('../figs/RRSanityTestWeights.png', dpi=300, bbox_inches='tight')

plt.show()
```

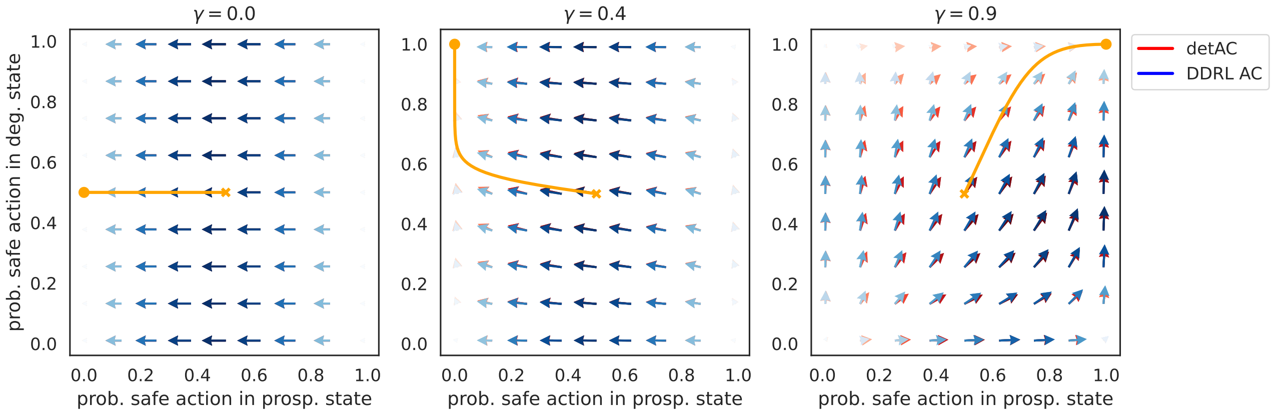

Figure 14.3: Comparison between a distributional and a non-distributional agent in a risk-reward dilemma, for several values of the discount factor. For the distributional agent, the weights method was used.

#### 14.2 Influence of optimism/pessimism

Since the flowplots obtained with the weights method are closest to the regular AC agents' flowplots, we use this method to implement optimism/pessimism. We study the influence of optimism for a low discount factor $\gamma = 0.4$, and for a high discount factor $\gamma = 0.9$. We assume that, in the degraded state, the agent always chooses the safe action (since it is the only one which can take it back to the prosperous state).

```
#  Levels of optimism/pessimism (defined by upper weight values w_U)
weights = np.array([0.1, 0.2, 0.4, 0.8, 1., 2., 4., 8., 10.])

# Phase space items to plot along the x-axis
x = ([0], [0], [0])

# Range and resolution
res = np.linspace(0, 1., 21)
res[0] = 0.0001
res[-1] = 0.9999

# Value of X^{0,1,0}
otherXs = np.ones((len(res)))
```

For a low discount factor $\gamma = 0.4$:

```
# Plot and save - gamma = 0.4
compare_levels(env, weights, gamma=0.4, beta=50, Rmin=0, Rmax=1, xinds=x,
               res=res, algo='AC', nrb=29, discsig=0.05,
               traj_len=800, otherXs=otherXs, cmap='coolwarm',
               save=True, savename='../figs/RRPhasePlotGamma04.png')

plt.show()
```

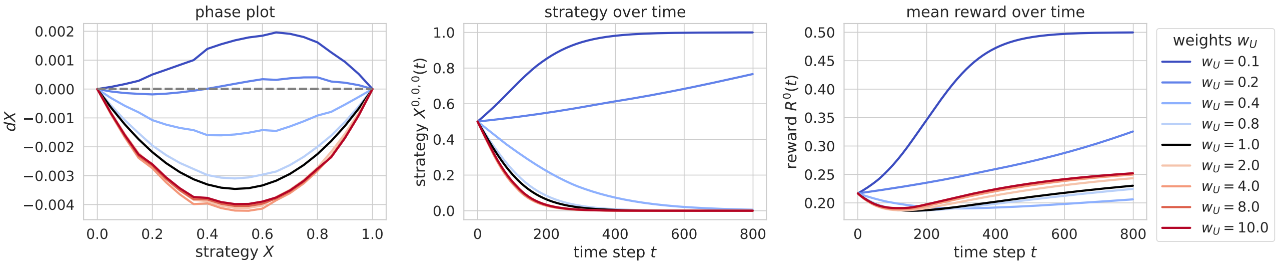

Figure 14.4: Phase plot, strategy plot and reward plot of an agent playing a risk-reward dilemma with low discount factor. Different levels of optimism/pessimism are compared.

For a high discount factor $\gamma = 0.9$:

```
# Plot and save - gamma = 0.9
compare_levels(env, weights, gamma=0.9, beta=50., Rmin=0, Rmax=1,
               xinds=x, algo='AC', res=res, nrb=29, discsig=0.05,
               traj_len=1800, otherXs=otherXs, cmap='coolwarm',
               save=True, savename='../figs/RRPhasePlotGamma09.png')
```

```
plt.show()
```

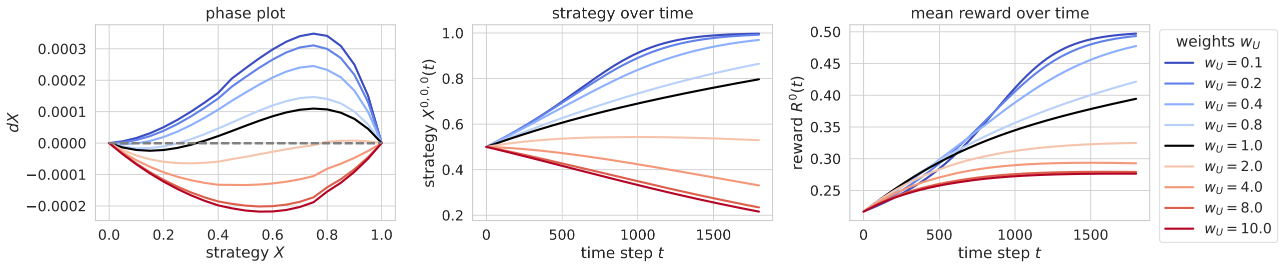

Figure 14.5: Phase plot, strategy plot and reward plot of an agent playing a risk-reward dilemma with high discount factor. Different levels of optimism/pessimism are compared.

We can visualize the impact of optimism/pessimism on the agent's strategy in both states, by using flowplots. Here, we show flowplots for a pessimistic agent ($w_U = 0.2$), a neutral agent ($w_U = 1$), and an optimistic agent ($w_U = 5$). First, we study how optimism/pessimism impacts the strategy of an agent with low discount factor $\gamma = 0.4$.

```
x = ([0], [0], [0])  # prosperous state
y = ([0], [1], [0])  # degraded state
w_list = [0.2, 1, 5]  # list of weights
```

```
# Define agents to compare
aeis = [] # DDRLActorCritic agents

for i, w in enumerate(w_list):
    aei = DDRLActorCritic(env, nr_reward_bins=29, discretization_sigma=0.05,
                          learning_rates=0.1, discount_factors=0.4,
                          Rmin=0, Rmax=1,
                          method="weights", wU=w, use_prefactor=True)
    aeis.append(aei)
```

```
# For trajectories
Xinits = [np.ones((1, 2, 2)) * 0.5]  # initial strategy
cols = ["orange"]  # trajectory color

# Decorations
titles = [r'$w = 0.2$', r'$w = 1$', r'$w = 5$']
xlabel = "prob. safe action in prosp. state"
ylabel = "prob. safe action in deg. state"

# Figure and axes
fig, (ax1, ax2, ax3) = plt.subplots(1, 3, figsize=(3.2 * 3.8, 3.8))
axes = [ax1, ax2, ax3]

# Plot flow plots and trajectories
plot_3flowplots(aeis, x, y, NrRandom=16, cmap='Blues', decorate=True,
                titles=titles, xlabel=xlabel, ylabel=ylabel,
```

```
                 Xinits=Xinits, cols=cols, axes=axes)

# Save the figure
plt.savefig('../figs/RRFlowplotsGamma04.png', dpi=300, bbox_inches='tight')

plt.show()
```

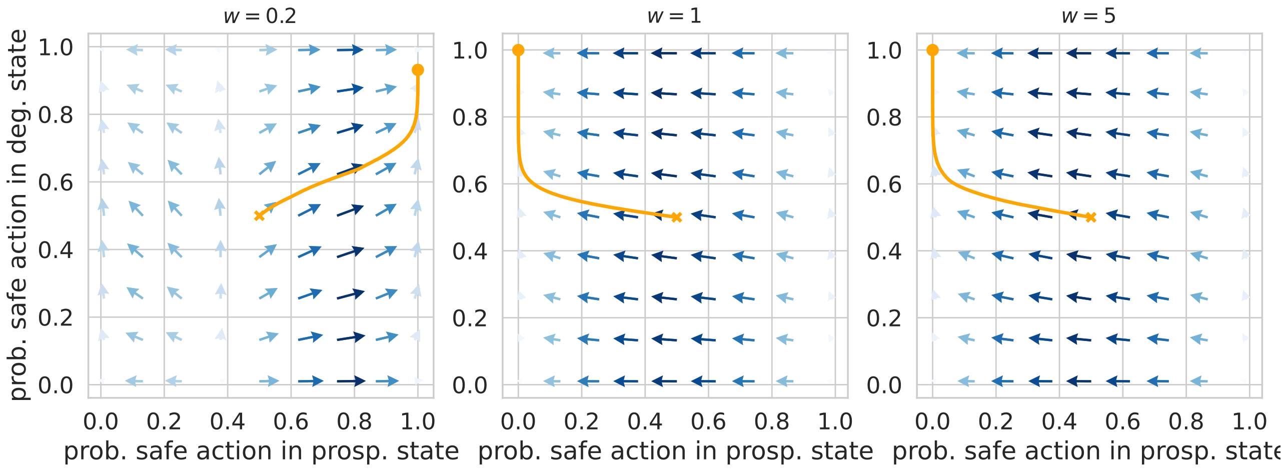

Figure 14.6: Flowplots for an agent playing a risk-reward dilemma with low discount factor. Different levels of optimism/pessimism are compared: pessimistic, neutral and optimistic.

We then turn to agents with a high discount factor $\gamma = 0.9$.

```
# Define agents to compare
aeis = [] # DDRLActorCritic agents

for i, w in enumerate(w_list):
    aei = DDRLActorCritic(env, nr_reward_bins=29, discretization_sigma=0.05,
                          learning_rates=0.1, discount_factors=0.9,
                          Rmin=0, Rmax=1,
                          method="weights", wU=w, use_prefactor=True)
    aeis.append(aei)
```

```
# For trajectories
Xinits = [np.ones((1, 2, 2)) * 0.5]  # initial strategy
cols = ["orange"]  # trajectory color

# Decorations
titles = [r'$w = 0.2$', r'$w = 1$', r'$w = 5$']
xlabel = "prob. safe action in prosp. state"
ylabel = "prob. safe action in deg. state"

# Figure and axes
fig, (ax1, ax2, ax3) = plt.subplots(1, 3, figsize=(3.2 * 3.8, 3.8))
axes = [ax1, ax2, ax3]

# Plot flow plots and trajectories
```

```
plot_3flowplots(aeis, x, y, NrRandom=16, cmap='Blues', decorate=True,
                titles=titles, xlabel=xlabel, ylabel=ylabel,
                Xinits=Xinits, cols=cols, axes=axes)

# Save the figure
plt.savefig('../figs/RRFlowplotsGamma09.png', dpi=300, bbox_inches='tight')

plt.show()
```

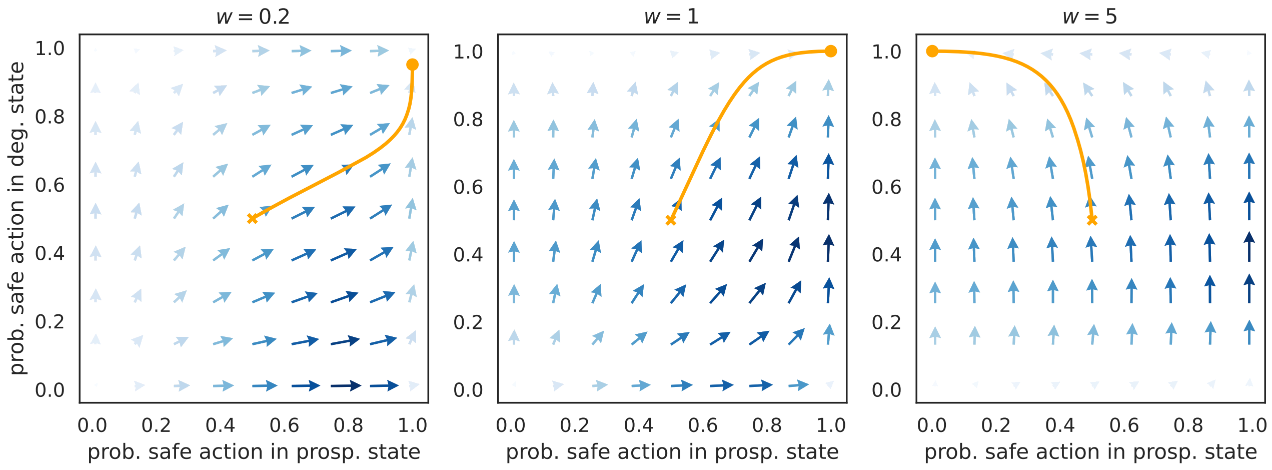

Figure 14.7: Flowplots for an agent playing a risk-reward dilemma with high discount factor. Different levels of optimism/pessimism are compared: pessimistic, neutral and optimistic.

#### 14.3 Bistability regimes and individual dilemma

In this section, we dive deeper into the particularities of our distributional model when combined to intertemporal choice. Indeed, Figure 14.6 shows the emergence of a bistability regime for distributional agents when $\gamma = 0.4$ and $w = 0.2$. In fact, we show that a bistability regime emerges even for neutral ($w = 1$) distributional agents, whereas neutral, non-distributional agents, do not exhibit bistability. This can be visualized in the Figure below:

```
# Discount factors
gammas = np.array([0.80, 0.84, 0.88, 0.92, 0.96, 1.])

# Phase space items to plot along the x-axis
x = ([0], [0], [0])

# Range and resolution
res = np.linspace(0, 1., 21)
res[0] = 0.0001
res[-1] = 0.9999

# Value of X^{0,1,0}
otherXs = np.ones((len(res)))
```

```
# Plot
compare_gammas(env, gammas, Rmin=0, Rmax=1, xinds=x,
               res=res, nrb=29, discsig=0.05,
               traj_len=800, otherXs=otherXs, cmap='viridis',
               save=True, savename='../figs/DistDetPhasePlots.png')

plt.show()
```

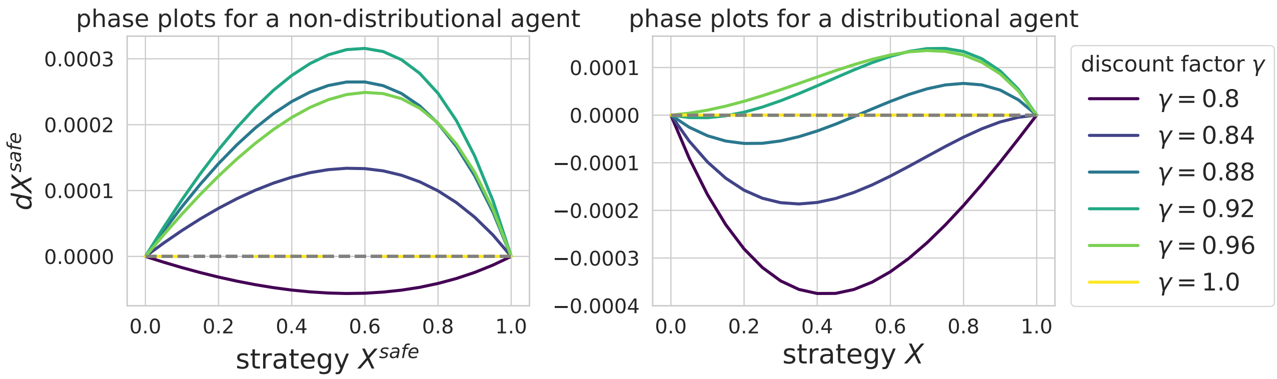

Figure 14.8: Phaseplots for a distributional vs. non-distributional agent playing a risk-reward dilemma. Different values of the discount factor are compared.

We can get a broader overview of the different stability regimes for various values of $w$ and $\gamma$:

```
# Safe and risky policies
Xsafe = np.array([[[0.999,0.001],[1,0]]])
Xrisk = np.array([[[0.001,0.999],[1,0]]])
```

```
discountfactors = np.linspace(0.01, 0.99, 21)
weights = np.array([0.1, 0.2, 0.3, 0.4, 0.5,
                    0.6, 0.7, 0.8,
                    0.9, 1., 2., 3., 4.,
                    5., 6., 7., 8., 9., 10.])
```

```
# risky_optimal_data_container, indicator_stable = fill_containers(env, Xrisk,
 ↪ Xsafe, discountfactors, weights)
```

```
# Extract data
df_optimal = pd.read_csv("../data/risky_optimal.csv")
df_indicator = pd.read_csv("../data/indicator_stable.csv")
```

```
# Convert to numpy array
risky_optimal_data_container = df_optimal.to_numpy()[:, 1:]
indicator_stable = df_indicator.to_numpy()[:, 1:]
```

```
# For drawing the black lines
```

```
risky_optimal_reverse = np.array(1 - risky_optimal_data_container).astype(int)
risky_bottom = np.sum(risky_optimal_reverse[:8, :], axis=0).astype(int)
risky_top = len(discountfactors) - np.sum(risky_optimal_reverse[8:, :], axis=0)
line_bottom = np.repeat(discountfactors[risky_bottom], 3)
line_top = np.repeat(discountfactors[risky_top], 3)
height = discountfactors[1] - discountfactors[0]
line_bottom = line_bottom - height/2
line_top = line_top - height/2
```

```
# Plot

from matplotlib.colors import ListedColormap, BoundaryNorm
from matplotlib.patches import Rectangle

hatches = {0.: '///', 1.: "\\\\\\"}
labels = {1.: r'$V_{risky} > V_{safe}$', 0.: r'$V_{safe} \geq V_{risky}$'}
x_vals = weights
y_vals = discountfactors
nx = len(x_vals)
ny = len(y_vals)

fig, ax = plt.subplots(1,1, figsize=(7,7))

vals = {10}
x_points = np.zeros((len(weights)*3))
x_points[0] = 0.09
x_points[-1] = 10.5
x_points[1::3] = weights

for (i, j), val in np.ndenumerate(risky_optimal_data_container):
    if j < nx - 1:
        width = x_vals[j+1] - x_vals[j]
        xp1 = x_vals[j+1]-width/2-0.0001
        xp2 = x_vals[j+1]-width/2+0.0001
        x_points[3*j+2] = xp1
        x_points[3*j+3] = xp2
    else:
        width = x_vals[j] - x_vals[j-1]

    if i < ny - 1:
        height = y_vals[i+1] - y_vals[i]
    else:
        height = y_vals[i] - y_vals[i-1]

    if val not in vals:
        label = labels[val]
        vals.add(val)
        # Draw rectangle with no fill but black edge and hatch pattern
        rect = Rectangle((x_vals[j] - width/2, y_vals[i] - height/2), width, height,
                         fill=False, edgecolor='black', hatch=hatches[val],
                         linewidth=0, label=label)  # no visible border line
    else:
```

```
        rect = Rectangle((x_vals[j] - width/2, y_vals[i] - height/2), width, height,
                          fill=False, edgecolor='black', hatch=hatches[val],
                          linewidth=0)  # no visible border line

    if x_vals[j]==1:  # Special case
        width=0.55
        xp1 = x_vals[j+1]-0.5-0.0001
        xp2 = x_vals[j+1]-0.5+0.0001
        x_points[3*j+2] = xp1
        x_points[3*j+3] = xp2
        rect = Rectangle((x_vals[j] - 0.05, y_vals[i] - height/2), width, height,
                          fill=False, edgecolor='black', hatch=hatches[val],
                          linewidth=0)  # no visible border line

    ax.add_patch(rect)
    # if (indicator_highlight[i, j] == 2) or (indicator_highlight[i, j] == 5):
        # text = ax.text(x_vals[j], y_vals[i], '*',
                       # ha="center", va="center", color="w")

# Create a discrete colormap with colors for 1, 2, 3
cmap = ListedColormap(['red', 'blue', 'purple'])

# Define boundaries between your discrete values
bounds = [0.5, 1.5, 2.5, 3.5]
norm = BoundaryNorm(bounds, cmap.N)

ax.pcolormesh(weights, discountfactors,
              indicator_stable, cmap=cmap, norm=norm, alpha=.6)
ax.plot(x_points, line_bottom, color='k')
ax.plot(x_points, line_top, color='k')

ax.set_xlabel('Weight '+r'$w_U$', fontsize=16)
ax.set_xscale('log')
ax.set_xlim([0.09, 10.5])
ax.set_ylim([0.01, 0.99])

ax.plot([], [], color='red', alpha=.6, label="risky stable")
ax.plot([], [], color='blue', alpha=.6, label="safe stable")
ax.plot([], [], color='purple', alpha=.6, label="both stable")
ax.legend(bbox_to_anchor=(1.01, 1.01), fontsize=14)

ax.tick_params(axis='both', which='major', labelsize=12)
ax.tick_params(axis='both', which='minor', labelsize=12)

ax.set_ylabel('Discount factor '+r'$\gamma$', fontsize=16)
ax.set_title('Stability landscape', fontsize=18)

# Save the figure
plt.savefig('../figs/StabilityLandscape.png', dpi=300, bbox_inches='tight')

plt.show()
```

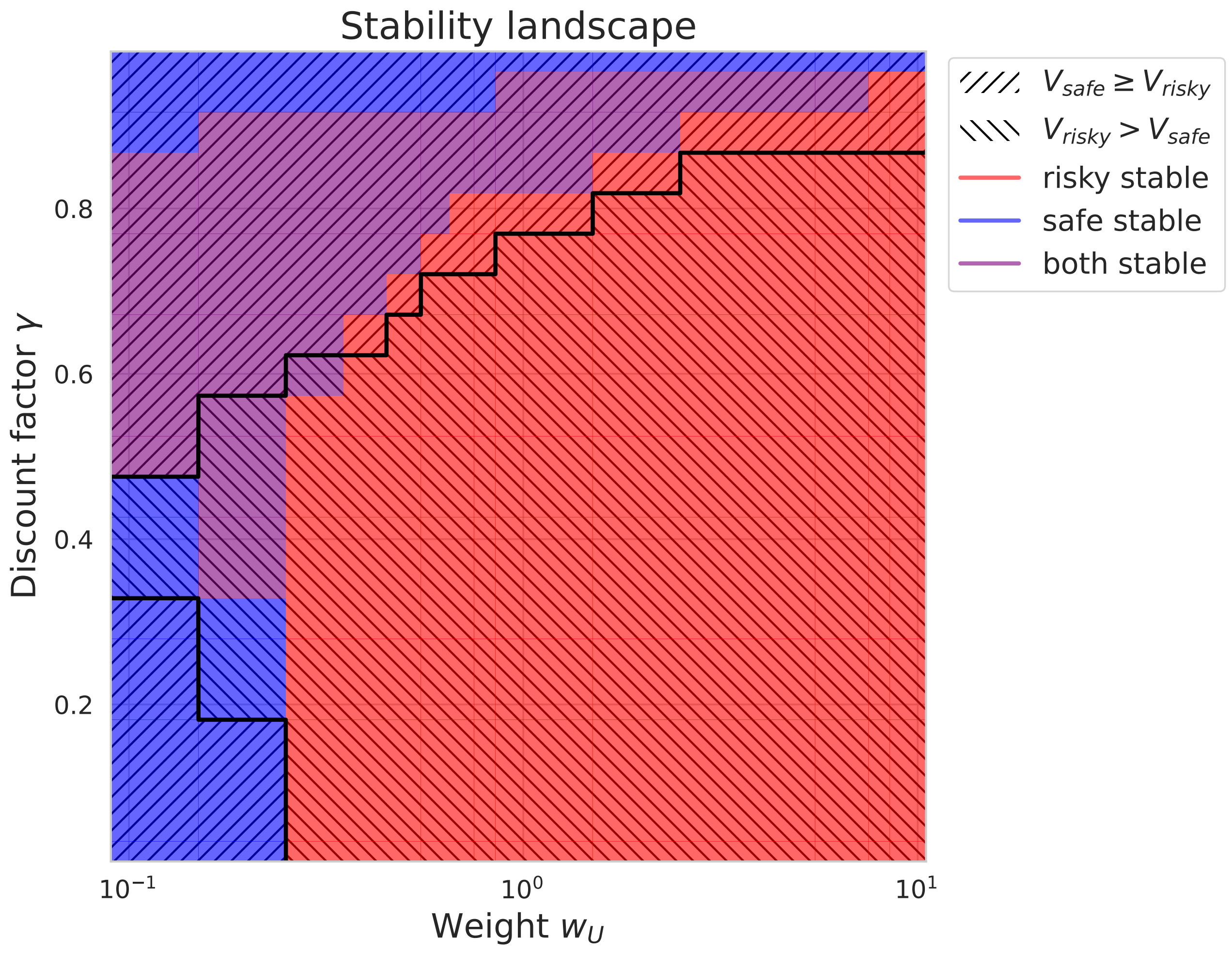

Figure 14.9: Stability landscape for an agent playing a risk-reward dilemma, for different values of the optimism weight and the discount factor.

Interestingly, Figure 14.9 shows zones where the safe strategy is stable, while the risky strategy yields a higher value overall (and vice-versa). This occurs even for neutral agents ($w_U = 1$). We coin this kind of incoherent choice an "individual dilemma". Below, we visualize the flowplots showing this individual dilemma at $w_U = 1$, $\gamma = 0.794$. Note that this individual dilemma does not occur for non-distributional agents.

```
# Individual dilemma at w=1, gamma=0.794

aei = DDRLActorCritic(env, nr_reward_bins=29, discretization_sigma=0.05,
                      learning_rates=0.1, discount_factors=0.794,
                      Rmin=0, Rmax=1,
                      method="weights", wU=1, use_prefactor=True)

mae = stratAC(env, learning_rates=0.1, discount_factors=0.794, use_prefactor=True)
```

```
# Flowplots

x = ([0], [0], [0])  # prosperous state
y = ([0], [1], [0])  # degraded state
```

```
fig, (ax1, ax2) = plt.subplots(1, 2, figsize=(6,3))
fp.plot_strategy_flow(aei, x, y, flowarrow_points = np.linspace(0.01 ,0.99, 9),
                      NrRandom=16, axes=ax1)
fp.plot_strategy_flow(mae, x, y, flowarrow_points = np.linspace(0.01 ,0.99, 9),
                      NrRandom=16, axes=ax2)
ax1.set_title("Distributional")
ax2.set_title("Non-distributional")
ax1.set_ylabel("prob. safe in deg. state")
ax1.set_xlabel("prob. safe in prosp. state")
ax2.set_xlabel("prob. safe in prosp. state")

plt.show()
```

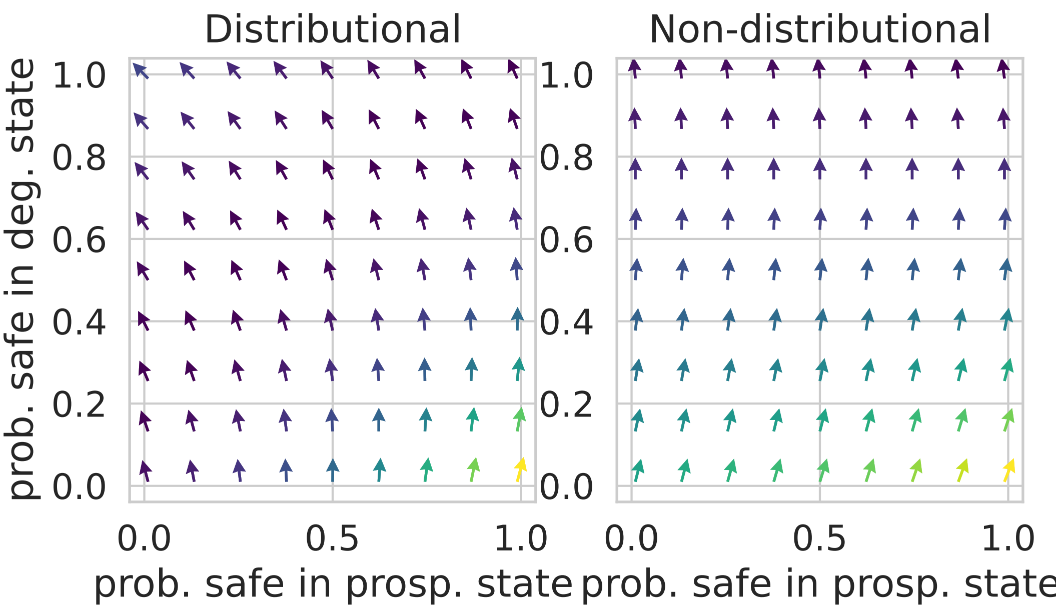

We can also plot the distributional agent's state-value and action-state-value distributions at this individual dilemma. Figure 14.10 shows that, although the risky policy is stable, the safe policy yields a higher state value.

```
# Value distributions

Visv_risk = aei.Visv(Xrisk)
Visv_safe = aei.Visv(Xsafe)

# Means

vis_risk = np.einsum('ijk,k->ij', Visv_risk, aei.gridpoints)
vis_safe = np.einsum('ijk,k->ij', Visv_safe, aei.gridpoints)

# State-Action-value distributions

Visav_risk = aei.Visav(Xrisk)
Visav_safe = aei.Visav(Xsafe)

# Means
```

```python
visa_risk = np.einsum('ijkl,l->ijk', Visav_risk, aei.gridpoints)
visa_safe = np.einsum('ijkl,l->ijk', Visav_safe, aei.gridpoints)
```

```python
# Plot

fig, ((ax1, ax2), (ax3, ax4)) = plt.subplots(2, 2, figsize = (9, 12))

ax1.bar(aei.gridpoints, Visv_safe[0,0,:], width=0.9*aei.gridwidth, color="blue",
 ↪  alpha=0.4)
ax2.bar(aei.gridpoints, Visv_risk[0,0,:], width=0.9*aei.gridwidth, color="red",
 ↪  alpha=0.4)
ax1.plot([vis_safe[0, 0], vis_safe[0, 0]], [0, 0.02], color="blue")
ax2.plot([vis_risk[0, 0], vis_risk[0, 0]], [0, 0.005], color="red")
ax2.plot([vis_safe[0, 0], vis_safe[0, 0]], [0, 0.005], color="blue", ls='--')
ax1.plot([vis_risk[0, 0], vis_risk[0, 0]], [0, 0.02], color="red", ls='--')
ax1.set_title("Value distribution for safe policy")
ax2.set_title("Value distribution for risky policy")
ax1.set_xlabel(r'$v$')
ax2.set_xlabel(r'$v$')
ax1.set_ylabel(r'$V^{i,prosp,v}$')
ax3.bar(aei.gridpoints, Visav_safe[0,0,0,:],
        width=0.9*aei.gridwidth, color="blue", alpha=0.4)
ax3.bar(aei.gridpoints, Visav_safe[0,0,1,:],
        width=0.9*aei.gridwidth, color="red", alpha=0.4)
ax4.bar(aei.gridpoints, Visav_risk[0,0,0,:],
        width=0.9*aei.gridwidth, color="blue", alpha=0.4, label="safe action")
ax4.bar(aei.gridpoints, Visav_risk[0,0,1,:],
        width=0.9*aei.gridwidth, color="red", alpha=0.4, label="risky action")
ax3.plot([visa_safe[0, 0, 0], visa_safe[0, 0, 0]], [0, 0.02], color="blue")
ax3.plot([visa_safe[0, 0, 1], visa_safe[0, 0, 1]], [0, 0.02], color="red")
ax4.plot([visa_risk[0, 0, 0], visa_risk[0, 0, 0]], [0, 0.005], color="blue")
ax4.plot([visa_risk[0, 0, 1], visa_risk[0, 0, 1]], [0, 0.005], color="red")
ax3.set_title(r'$V^{i,s,a,v}(X^{safe})$')
ax4.set_title(r'$V^{i,s,a,v}(X^{risk})$')
ax4.legend(bbox_to_anchor=(1.01, 1.01))
ax3.set_xlabel(r'$v$')

plt.show()
```

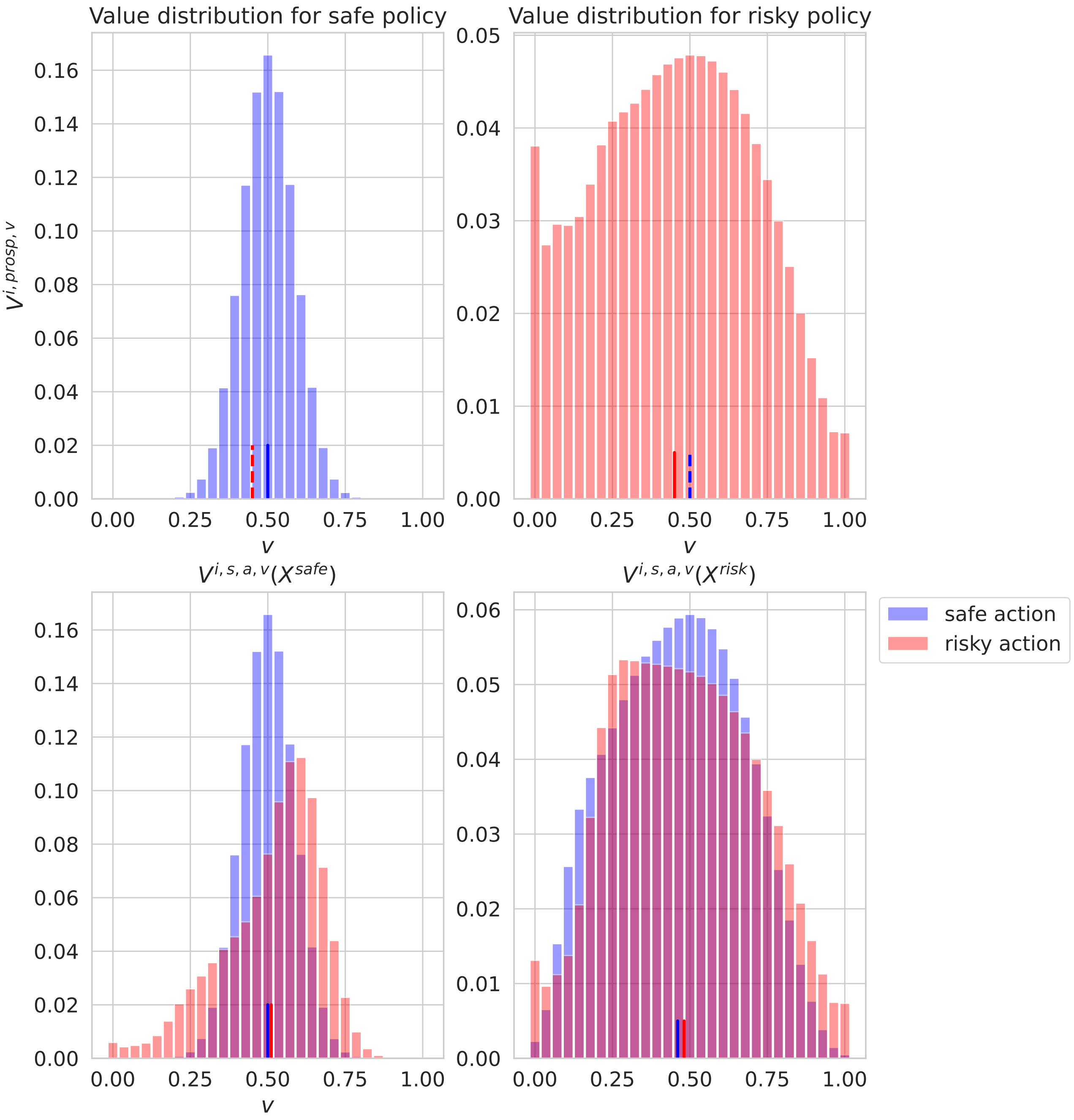

Figure 14.10: State-value and action-state-value distributions for a distributional agent playing a risk-reward dilemma.
